# Meisoindigo: An Effective Inhibitor of SARS-CoV-2 Main Protease Revealed by Yeast System

**DOI:** 10.1101/2023.09.03.555867

**Authors:** Wojciech Grabiński, Anna Kicińska, Ewa Kosicka, Martyna Baranek-Grabińska, Ewelina D Hejenkowska, Joanna Budzik, Paulina Śliska, Weronika Śliwińska, Andonis Karachitos

## Abstract

The COVID-19 pandemic caused by SARS-CoV-2 has had a significant impact on global health and the global economy. Despite the availability of vaccines, limited accessibility and vaccine hesitancy pose challenges in controlling the spread of the disease. Effective therapeutic strategies, including antiviral drugs, are needed to combat the future spread of new SARS-CoV-2 virus variants. The main protease (Mpro) is a critical therapeutic target for COVID-19 medicines, as its inhibition impairs viral replication. However, the use of substances that inhibit Mpro may induce selection pressure. Thus, it is vital to monitor viral resistance to known drugs and to develop new drugs. Here, we have developed a yeast system for the identification of Mpro inhibitors as an alternative to costly and demanding high-biosecurity procedures. The system is based on stable expression of Mpro and does not require selection media. Yeast can be cultured on a rich carbon source, providing rapid growth and screening results. The designed tool was subsequently used to screen the FDA-Approved Drug Library. Several chemicals with Mpro inhibitory properties were identified. We found that meisoindigo, which was not previously known to have the potential to inhibit Mpro, was highly effective. Our results may promote the development of new derivatives with therapeutic properties against SARS-CoV-2 and other beta-coronaviruses.

## 1. Introduction

The COVID-19 pandemic, caused by the SARS-CoV-2 coronavirus, has had an unprecedented global impact on public health, the economy, and everyday life. Despite the rapid development and deployment of vaccines against SARS-CoV-2, which have proven effective in improving patient outcomes and reducing the transmission of the disease, vaccine availability remains limited in certain regions. Additionally, vaccine hesitancy further complicates global vaccination efforts. Thus, controlling the spread of the virus and mitigating the impact of COVID-19 worldwide remain ongoing challenges (Troiano and Nardi, 2021).

In addition, SARS-CoV-2 continues to undergo genetic mutations, particularly in the spike protein (Yuan et al., 2021). These changes may result in a reduced immune response, thereby compromising the efficacy of vaccines and the effect of previous infections. Thus, there is an urgent need for effective therapeutic strategies to combat the spread of new variants of SARS-CoV-2. The development of antiviral substances that target viral replication is essential to mitigate the impact of emerging variants. These drugs can play a critical role in controlling infection, reducing disease severity, and preventing further transmission of SARS-CoV-2.

The main protease (M^pro^), also named 3-chymotrypsin-like protease (3CLpro), is considered to be the key target for COVID-19 medicines, as it is highly conserved among coronaviruses and plays an important role in the viral life cycle. M^pro^ cleaves the polyproteins that are translated from viral RNA (Amin et al., 2021). Inhibition of M^pro^ function can disrupt viral replication and could be highly effective not only against the current pandemic but also against other fatal coronavirus-caused diseases (Ma et al., 2020).

In fact, multiple strategies have been used to screen for effective M^pro^ inhibitors, including drug repurposing, virtual screening and structure-based drug design (Hu et al., 2022). These studies resulted in numerous compounds with M^pro^ inhibitory potential, one of which (paxlovid) is already used in clinical practice, and a few others are strong candidates (Hu et al., 2022).

However, the use of drugs targeting M^pro^ can create selection pressure and contribute to the development of resistance (Krishnamoorthy and Fakhro, 2021). Therefore, it is essential to continuously monitor and test for virus resistance to known substances and better understand the mechanisms of enzyme action.

In fact, recent studies investigating M^pro^ activity primarily involve *in vitro* assays with isolated proteins that allow the determination of enzymatic kinetics and functional assays aimed at understanding viral replication in the presence of inhibitors (De Castro et al., 2022; Kitamura et al., 2022). Moreover, it has recently been proposed to use yeast as a system for M^pro^ expression (Alalam et al., 2021; Flynn et al., 2022; Ou et al., 2023). The yeast system provides an alternative to techniques that measure M^pro^ activity in mammalian cells and addresses their several limitations. It reduces associated costs and eliminates the need for a high level of biosecurity.

The use of yeast as an expression system for M^pro^ also offers several other advantages. Yeast cells can be easily cultured and manipulated in the laboratory, enabling efficient expression of the viral protease. This system allows researchers to study the enzymatic activity and functional effects of M^pro^ in living cells.

The yeast system provides rapid access to the effects of potential inhibitors in a cellular context, providing valuable insight into compound entry, stability, and efficacy. This approach offers flexibility, as rapid adjustment of the sensitivity and potential modification of the amino acid sequence of M^pro^ to accommodate future mutant variants or enzymes from other betacoronaviruses is straightforward.

In this study, we present a novel yeast system specifically designed for the identification of M^pro^ inhibitors. The key feature of this system is the stable expression of M^pro^, which is achieved by integrating the Mpro gene into the *GAL1* region of yeast chromosome 2. This integration allows precise control of M^pro^ expression by galactose induction (Peng et al., 2015). Notably, this system does not require the use of selection media, and cells can be cultured on a rich carbon source such as sucrose, allowing for rapid growth and accelerated screening results.

In our M^pro^-expressing strain, a unique design for the N-terminal fragment of the protein was implemented. This design not only reconstitutes the native primary structure of M^pro^, where the first amino acid is serine but also leads of the release of an N-terminal EGFP fusion protein. This enhances the enzymatic activity of M^pro^, resulting in a strong phenotypic effect characterized by toxicity and inhibition of yeast growth. In addition, the fusion of EGFP to M^pro^ provides a sensitive analytical tool for assessing M^pro^ activity.

Our yeast system provides a comprehensive and sensitive approach for the identification of M^pro^ inhibitors. The controlled expression of M^pro^, the phenotypic effects induced by its activity, and the analytical tool of EGFP fusion provide valuable tools for screening and characterizing potential inhibitors. The system was validated with the use of known M^pro^ inhibitors and allowed us to discover a new inhibitor, meisoindigo.

Thus, our yeast-based screening approach provides a valuable tool for the discovery and evaluation of M^pro^ inhibitors, which can ultimately lead to the development of novel drugs and enhance our ability to combat COVID-19 and other related viral infections.

## 2. Results and Discussion

### 2.1. The yeast system

In the development of our yeast system, our primary goal was to establish stable expression of the target protein, specifically M^pro^, which was intentionally designed to be toxic to cells. The use of classical vector-based expression systems would require the implementation of special selection pressure conditions, such as minimal media, which cannot adequately support robust culture growth. To overcome this challenge, we chose to use the CRISPR/Cas9 system to integrate the M^pro^-expressing gene directly into the yeast chromosome (Fig 1A). This system consists of a vector containing Cas9, sgRNA, and a separate vector containing the repair DNA. An important advantage of our system was its ability to facilitate rapid modification and redesign of the yeast system, allowing us to efficiently express mutant variants of M^pro^.

**Fig 1.**
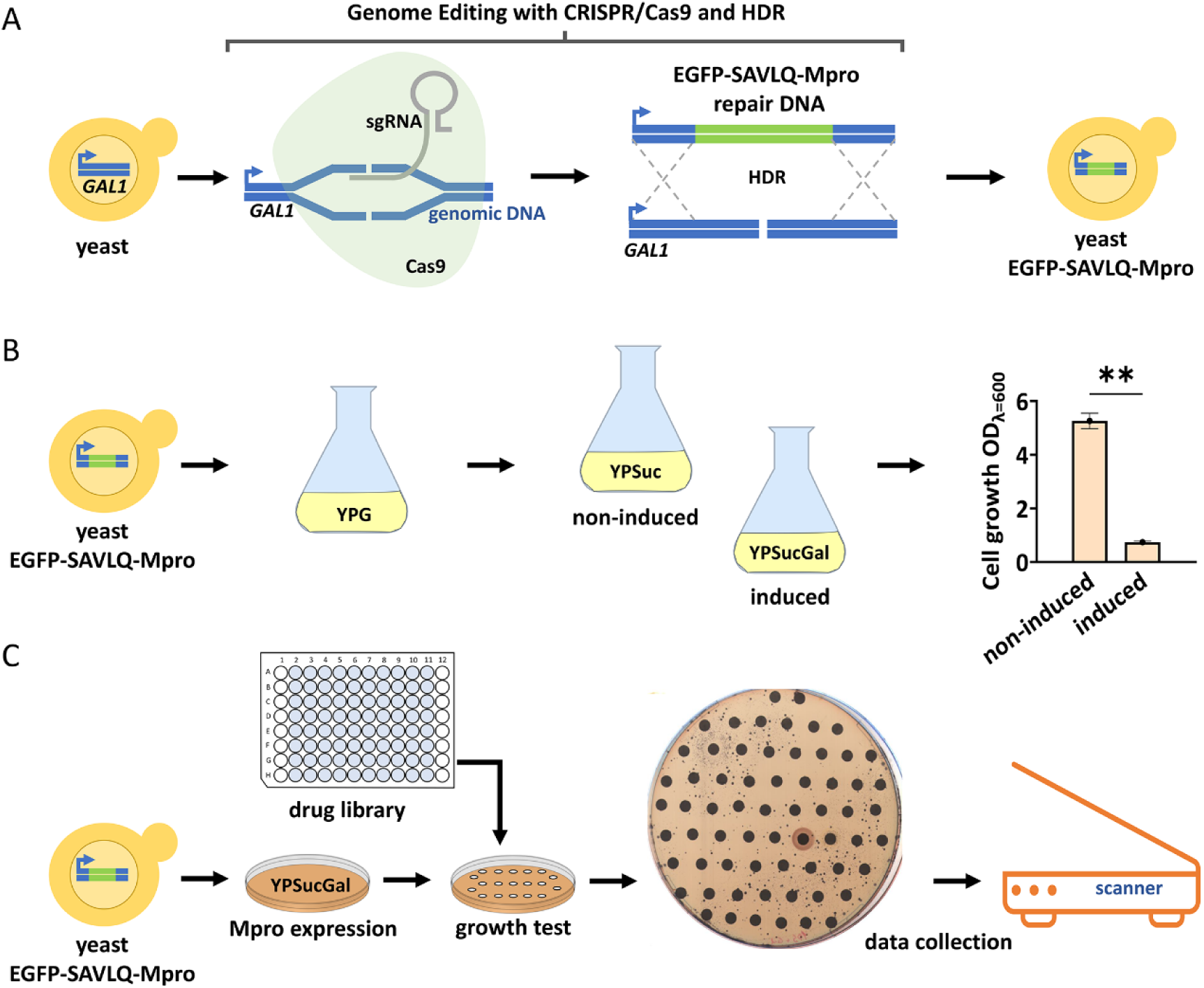
The yeast system designed for the identification of M^pro^ inhibitors. **A)** Illustration showing the method of EGFP-SAVLQ-Mpro encoding gene integration into the yeast genome. The CRISPR/Cas9 system recognizes a specific site within *GAL1* that causes a double-strand break. By flanking homology to regions downstream and upstream of *GAL1* within the repair DNA, homology-directed repair (HDR) and the integration of a new gene at the *GAL1* site take place. By using this technique, the EGFP-SAVLQ-Mpro coding gene is placed under the control of the GAL1 promoter. **B)** Differences in yeast culture growth depending on M^pro^ expression. The asterisks (**) indicate that the p value is less than 0.01 (n=3). Error bars represent the standard deviation (SD). **C)** Utilizing a yeast system to screen a drug library for M^pro^ inhibitors. Yeast culture is spread homogeneously on a plate with solid medium, which leads to the induction of M^pro^ expression and induces growth suppression. The application of filters containing drugs from the library allows high-throughput screening of potential M^pro^ inhibitors based on differences in culture growth.

By integrating the *M^pro^* gene into the *GAL1* locus, we achieved precise control over its expression by galactose addition to the growth medium. However, it is important to note that the gene knock-in procedure resulted in the removal of the *GAL1* gene from the yeast strain. As a result, the yeast strain no longer possesses the ability to consume galactose as a carbon source (Bhat et al., 1990). Thus, small concentrations of galactose in the medium effectively induce M^pro^ expression. In addition, in the context of drug screening, glucose-rich medium cannot be used due to the phenomenon of catabolic repression. High glucose concentrations inhibit the induction of gene expression by galactose (Gancedo, 1998). Consequently, the *GAL1* promoter remains inactive in the absence of galactose or in the presence of repressive carbon sources (glucose), resulting in minimal or no gene expression.

The next goal in designing the system was to eliminate yeast multidrug resistance, which could potentially interfere with screening results. We chose to delete three specific genes: *PDR5* and *SNQ2,* encoding multidrug resistance transporters, and *PDR1*, a transcription factor that regulates pleiotropic drug response (Decottignies et al., 1998). By removing these genes from the yeast genome, we aimed to minimize the confounding effects of multidrug resistance and improve the accuracy of our drug screening results. The deletion of the three targeted genes significantly changed the phenotype of the yeast strain expressing M^pro^ - EGFP-SAVLQ-Mpro. However, this change rendered the screening process infeasible. Interestingly, the mutant strains (EGFP-SAVLQ-Mpro Δpdr5, Δsnq2 and EGFP-SAVLQ-Mpro Δpdr5, Δsnq2, Δpdr1) showed significantly enhanced growth capabilities compared to the original strain with only one *PDR5* gene deleted, -EGFP-SAVLQ-Mpro (see Fig S2). This unexpected improvement in the growth ability of the mutants underscored the complex interplay between multidrug resistance and yeast physiological responses. While it posed a challenge to our drug screening efforts, it also shed light on the intricate mechanisms underlying drug resistance in yeast. The observed phenomenon of enhanced growth in our mutant strains, despite the deletion of the three selected genes, remains unexplained. Interestingly, a previously reported yeast system in which M^pro^ is expressed in a strain lacking *PDR1, SNQ2* and *PDR3* resulted in significantly weaker cytotoxicity induced by M^pro^ expression compared to our variant that had only a *PDR5* deletion (Alalam et al., 2021). The different results observed between the two systems highlight the complexity of the factors influencing M^pro^ expression-induced cytotoxicity. These results underscore the importance of further investigations to elucidate the underlying mechanisms. In our case, the culture conditions and yeast phenotype required the use of the *pdr5*Δ mutant for optimal results in terms of drug screening.

One of the primary challenges we faced was selecting the optimal culture conditions to accelerate the drug screening process. Our initial approach was to test drugs on a glycerol-supplemented medium. Although we observed a significant change in the yeast phenotype upon the induction of *M^pro^* expression (Fig 2C), our attempts to screen the drug library using a nonfermentable carbon source were unsuccessful. None of the drugs showed any effect on the yeast phenotype (see Fig S3). In addition, the screening process was approximately one week long, which was too long for our needs. Galactose can strongly activate the *GAL1* gene when yeast are grown on glycerol media. However, glycerol itself is a weak carbon source, which may have contributed to the limited success of our drug screening approach (Hashimoto et al., 1983). To address the limitations encountered, the same authors proposed an alternative carbon source: sucrose (Hashimoto et al., 1983). This choice offered several advantages, including significantly enhanced cell growth compared to glycerol and minimal catabolic repression. We found that under optimized growth conditions, yeast cells exhibited significant cytotoxicity following the induction of M^pro^ with galactose. This cytotoxic effect, believed to be a result of the overexpression of M^pro^, led to a clear reduction in the growth rates of the yeast (Fig 1B). By using sucrose as the carbon source in our system, drug screening could be performed in a much shorter time interval. Remarkably, screening results were visible after only 24 hours, and the entire experiment was completed within a maximum of 48 hours. This timeline greatly accelerated the drug discovery process (Fig 1C).

**Fig 2.**
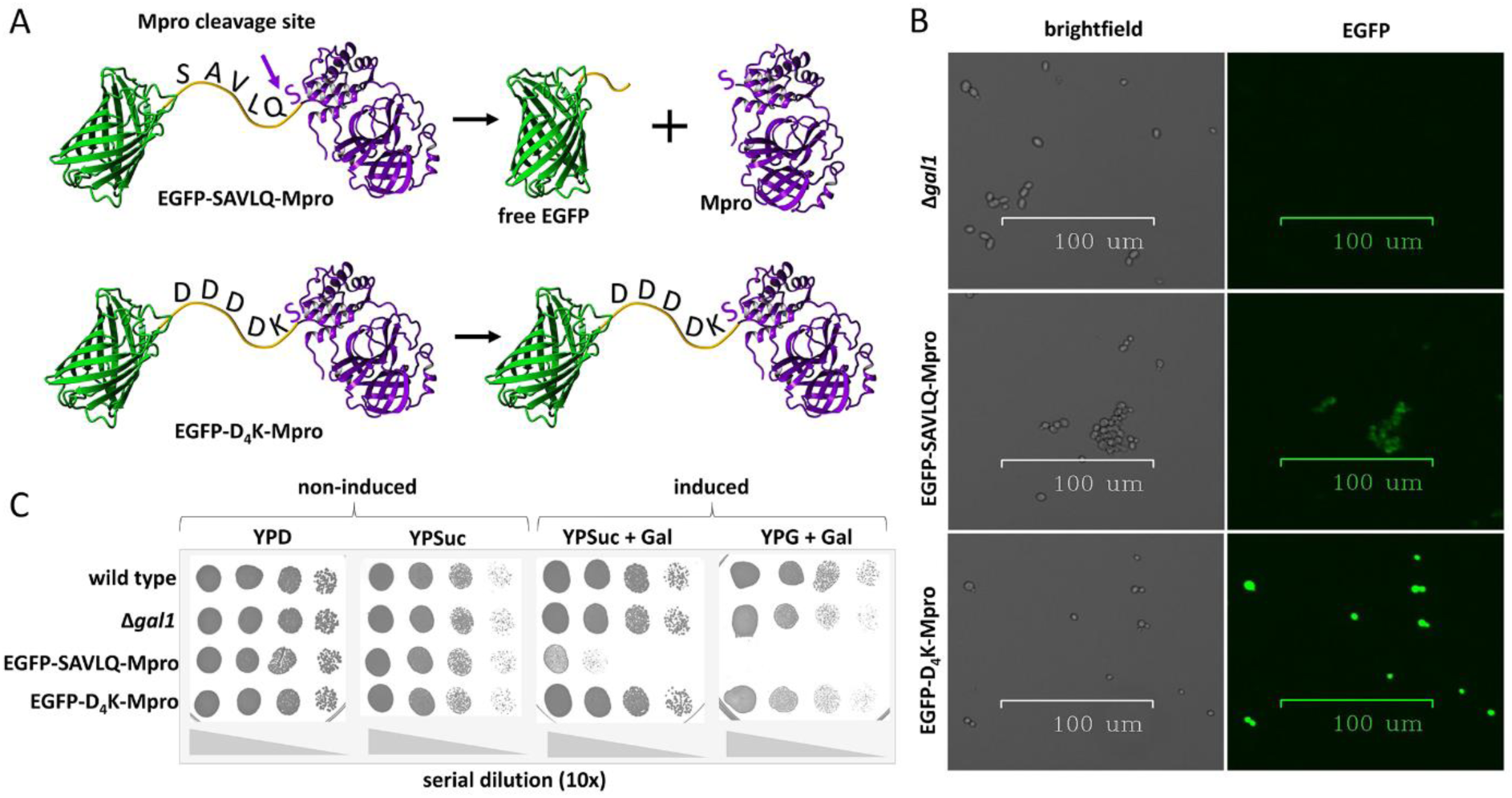
EGFP-M^pro^ linkers and their effect on M^pro^ function and expression. **A)** An illustration showing the structure of the fusion proteins expressed in the yeast used in this work. EGFP-SAVLQ-Mpro contains EGFP at the N-terminus, followed by a sequence (SAVLQ) that is recognized and autocatalytically cleaved by M^pro^. As a result, EGFP is released, and M^pro^ reconstitutes the native structure and is fully active. Replacing the sequence recognized by M^pro^ with the sequence recognized by enterokinase (D_4_K) provides the same linker length but prevents M^pro^ from cleaving off the N-terminal EGFP *in vivo* and consequently inhibits Mpro’s enzymatic activity. YASARA was used to visualize and render the final Fig of the fused EGFP (PDB: 6YLQ) and M^pro^ (PDB: 7QT5) protein structures. **B)** Differences in green fluorescence of Δgal1 (no EGFP) and EGFP-SAVLQ-Mpro or EGFP- D_4_K-Mpro (EGFP expression) strains. The images were made with the ZOE Fluorescent Cell Imager (Bio-Rad). **C)** Differences in the growth of cultures of different strains depending on the medium used and the type of linker used between EGFP and M^pro^.

A notable feature of our yeast system is the fusion of EGFP to the N-terminus of M^pro^ with SAVLQ linker. This allows autocatalytic cleavage by M^pro^ itself (as shown in Fig 2A). As a result, M^pro^ becomes fully active and subsequently inhibits yeast cell growth. The autocatalytic cleavage of M^pro^ mimics the process of M^pro^ expression in SARS-CoV. In viral life cycle, the polyproteins pp1a and pp1ab undergo autocatalytic cleavage to generate functional M^pro^ (V’kovski et al., 2021). The enzymatic activity of M^pro^ is inhibited by the presence of additional N- and C-terminal amino acids (Xue et al., 2007). We introduced a modified gene with a D_4_K linker between EGFP and M^pro^ to serve as a control strain without cytotoxicity upon M^pro^ expression. This linker prevents the enzymatic activity of M^pro^. As shown in Fig 2B, there was a marked difference in fluorescence between the EGFP-SAVLQ-Mpro and EGFP-D_4_K-Mpro strains. The EGFP-D_4_K-Mpro strain, which lacks the M^pro^ cleavage site, exhibits significantly higher green fluorescence than the EGFP-SAVLQ-Mpro strain. Consistent with this observation, the yeast phenotype (as shown in Fig 2C) confirms that the EGFP-D_4_K-Mpro strain does not exhibit growth inhibition, further supporting the role of the D_4_K linker in preventing M^pro^-induced cytotoxicity.

### 2.2. Screening the drug library

As described, the yeast system we developed is characterized by growth limitations induced by M^pro^ expression. The presence of M^pro^ exerts a cytotoxic effect on yeast cells, resulting in inhibited growth and compromised viability. This phenomenon is clearly demonstrated in Figure 1B, which provides the data on the impact of M^pro^ expression on yeast growth. Based on these results, the Z factor was calculated to assess the quality of our screening bioassay. The positive control (yeast cells expressing M^pro^ without inhibitor) showed significantly inhibited growth, with a mean optical density (OD_600_) of 0.74 ± 0.04. In contrast, the negative control (yeast cells without M^pro^ expression) exhibited normal growth, with a mean of 5. 26 ± 0.29. A Z factor of 0.782 indicates that our assay demonstrates a high level of assay quality, with sufficient dynamic range and low variability between the controls, suitable for high-throughput screening.

Prior to screening over 1,800 FDA-approved compounds from our library, we conducted an initial investigation to assess the system’s ability to effectively identify M^pro^ inhibitors. We used GC376, a control compound known for its activity against the 3CLpro of several coronaviruses, including SARS-CoV and SARS-CoV-2 (Shi et al., 2021).

The use of GC376 in our system yielded a positive result. The yeast strains expressing M^pro^ showed increased growth specifically around the filter containing the drug (Fig 3A, induced). This finding validated the efficacy of GC376 as an M^pro^ inhibitor in our system. However, during the screening phase, we also observed that certain drugs, such as ebselen, which had shown promise as an M^pro^ inhibitor (Jin et al., 2020), had no detectable effect on our system (Fig 3B). This lack of response was attributed to the general toxicity of ebselen, which affected overall yeast growth.

**Fig 3.**
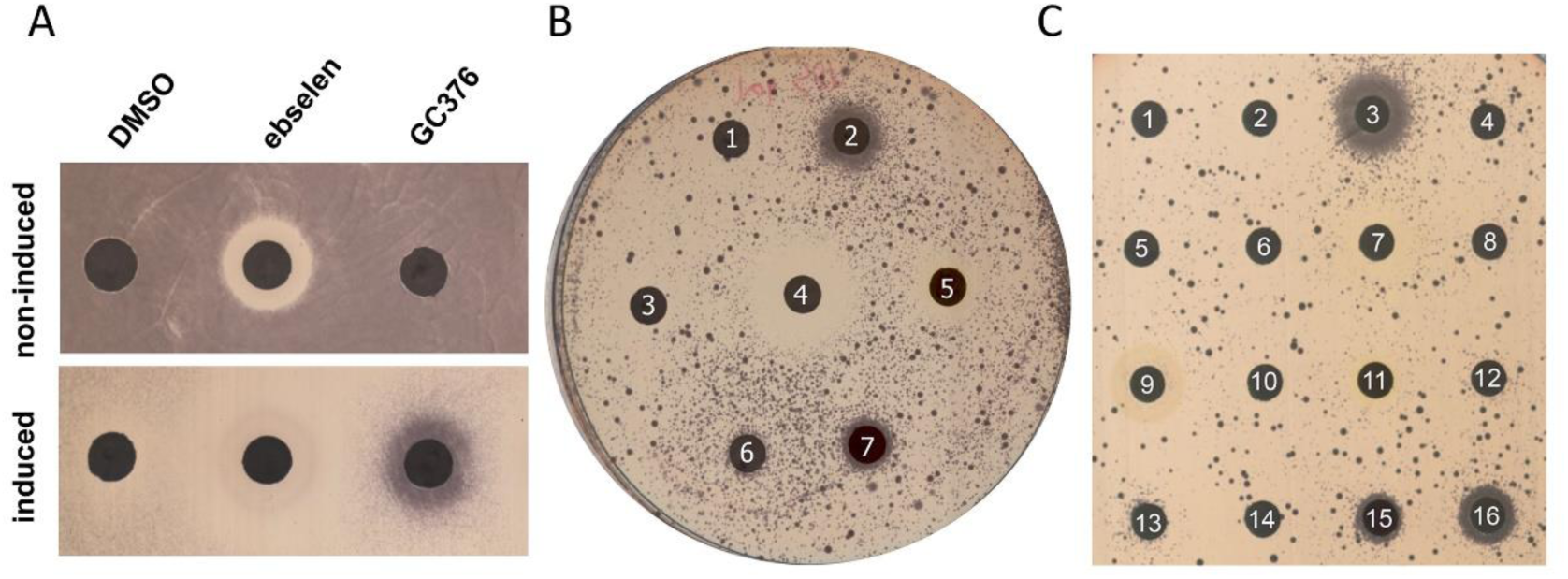
Effect of M^pro^ expression and potential inhibitors on yeast growth. **A)** Effects of a potential M^pro^ inhibitor (ebselen) and a known 3Cl protease inhibitor (GC376) on the growth of Mpro-expressing yeast. Limited growth of cultures with noninduced M^pro^ expression may indicate general cytotoxicity of the drug within its diffusion circle (ebselen), while increased growth of cultures with induced M^pro^ expression suggests an inhibitory effect on M^pro^ activity (GC376). **B)** The effect of the drugs selected in this work after the first screening on the growth of the yeast system for induced culture; 1 – DMSO or 10 mM drug: 2 – GC376, 3 – bedaquiline, 4 – berberine, 5 – ethacridine, 6 – ixazomib, 7- meisoindigo. **C)** The effect of selected drugs on the growth of yeast with M^pro^ expression depending on the drug concentration; 1 – empty filter, 2 – DMSO, 3 – 50 mM GC376, 4 – 5 mM GC376, 5 – 10 mM bedaquiline, 6 – 1 mM bedaquiline, 7 – 10 mM berberine, 8 – 1 mM berberine, 9 – 10 mM ciclopirox, 10 – 1 mM ciclopirox, 11 – 10 mM ethacridine, 12 – 1 mM ethacridine, 13 – 10 mM ixazomib, 14 – 1 mM ixazomib, 15 – 10 mM meisoindigo, 16 – 1 mM meisoindigo.

A comprehensive search of the drug library led to the identification of candidates with potential inhibitory effects on M^pro^: 9-aminoacridine, bedaquiline, berberine, ciclopirox, ethacridine, ixazomib, meisoindigo, menadione, pomalidomide and tedizolid (Supplementary file 2).

Notably, two of these identified drugs (ixazomib and meisoindigo) induced significant changes in the phenotype of the yeast strain expressing M^pro^ (Fig 3B, C). Some of the identified drugs have been previously reported as potential M^pro^ inhibitors and candidates for therapeutic agents. Inhibitory properties of ixazomib and pomalidomide have been suggested in in silico studies (Elzupir, 2020; Vázquez-Mendoza et al., 2022). Ethacridine, another drug identified in our screening, has shown inhibitory effects on SARS-CoV-2 virus particles (Li et al., 2021). The inhibitory effects observed in previous studies provide encouraging evidence for the potential of ethacridine as a therapeutic agent against SARS-CoV-2.

Despite the initial promising screening results, ixazomib, ethacridine and some of the other drugs were found to be only slightly effective in subsequent screening on liquid medium (Fig 4 presents ixazomib as a representative example). However, it is important to note that the possibility that ixazomib possesses M^pro^ inhibitory activity cannot be completely excluded. In the downstream steps, ixazomib serves as a control, representing a weak inhibitor compared to GC376 (Fig 4).

**Fig. 4.**
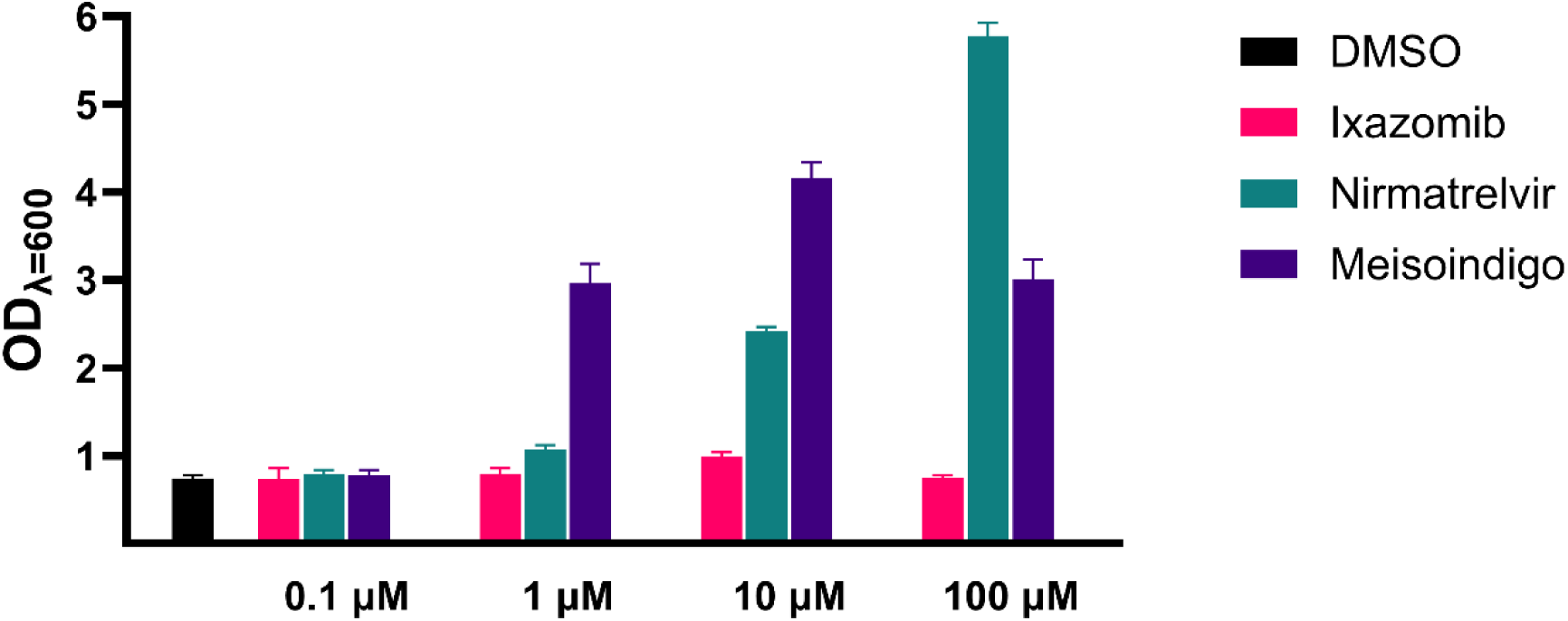
Concentration dependent effects of compounds on the growth of Mpro- expressing yeast. The OD_600_ of cultures 24 hours after induction, in the presence of tested compounds is shown. Thee control group was treated with vehicle DMSO. Error bars indicate the standard deviation (SD), n=3.

In addition to drugs that have been previously studied and reported, we identified several substances that have not previously been associated with M^pro^ inhibition. These drugs represent novel findings and were identified as potential M^pro^ inhibitors for the first time in our screening. Their identification in this study highlights their previously unexplored potential in targeting M^pro^ and suggests avenues for further investigation to determine their efficacy and mechanism of action.

Of particular importance is meisoindigo (for the chemical structure, see Fig S4), a derivative of indigo naturalis, an active compound in a Chinese antileukemia medicine with proven efficacy in the treatment of chronic myelogenous leukemia (CML) (Lee et al., 2010; Ye et al., 2019). In our studies, the performance of meisoindigo was particularly noteworthy (Fig 4). At a concentration of 10 µM, after 24 hours, the OD_600_ of yeast culture was 4.16 ± 0.17 vs. 0.74 ± 0.04 in the control culture (DMSO). Interestingly, at a higher concentration of 100 µM, meisoindigo did not protect yeast against M^pro^ toxicity that well (OD_600_ = 3.01 ± 0.23). This reduced efficacy at higher concentrations underscores the challenges posed by meisoindigo’s solubility in water. Meisoindigo, a second-generation derivative of indirubin, exhibits enhanced water solubility compared to its precursor (Ye et al., 2019). While this enhanced solubility facilitates its application in various media, it also seems to limit its efficacy at higher concentrations. The calculated IC_50_ values for GC376 and meisoindigo in yeast system are respectively µM 96.84 ± 14.75 and 0.597 ± 0.208 µM (Fig.5A). In contrast, nirmatrelvir, a component of the FDA-approved drug paxlovid used to treat COVID-19, demonstrated a mean yeast growth value of 2.42 ± 0.041 at 24 hours at a concentration of 10 µM (Fig 4). At the 100 µM concentration, the mean value of nirmatrelvir was significantly higher, 5.77 ± 0.15.

**Figure 5.**
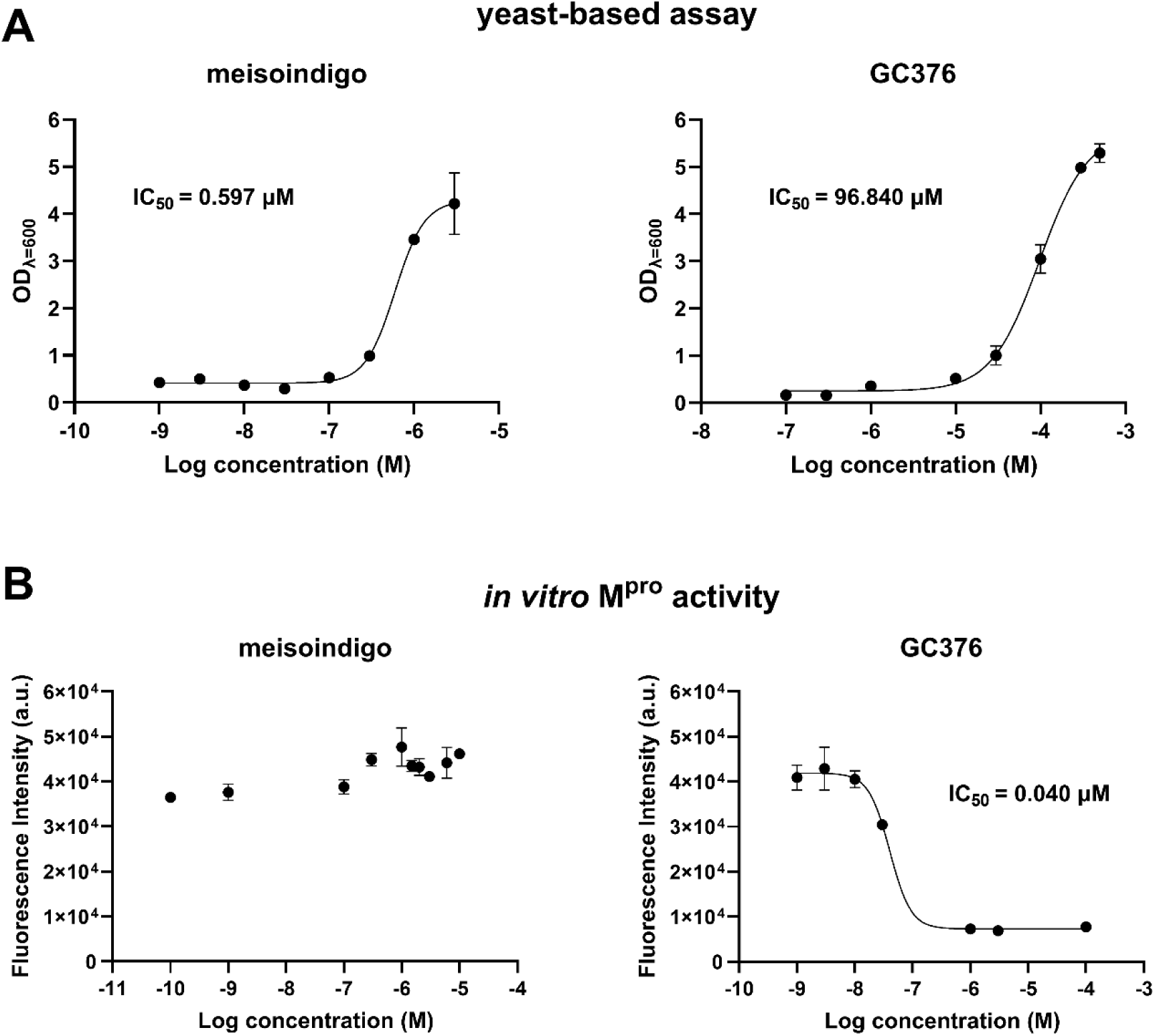
Comparison of inhibitor efficacy using yeast-based system and FRET *in vitro* assay. Evaluation of two drugs, GC376 and meisoindigo, as M^pro^ inhibitors using two distinct methods at various concentrations. In the yeast-based system, the efficacy of inhibitors was assessed based on yeast growth measurements, recorded as OD_600_ values. Lower OD_600_ values indicate higher M^pro^ activity. In the FRET-based in vitro assay, the efficacy of inhibitors was assessed through fluorescence measurements. Higher fluorescence values indicate higher M^pro^ activity. The IC_50_ values, which represent the concentration of the inhibitor required to reduce the M^pro^ activity by 50%, were calculated using nonlinear regression.

The solubility of meisoindigo in water, while presenting certain limitations, also offers intriguing possibilities for further research. Its enhanced solubility compared to that of indirubin suggests that there might be ways to further optimize its formulation for better results. Enhancing its solubility or stabilizing it at higher concentrations could potentially amplify the efficacy, making it a more potent agent in future applications.

The efficacy of meisoindigo as an inhibitor of the main protease of SARS-CoV-2 can be contextualized within the framework established by the research of Narkhede et al. (2020) (Narkhede et al., 2020). Their study highlighted the potential of indirubin, a compound structurally similar to meisoindigo, as an Mpro inhibitor. This structural similarity suggests a possible shared mechanism of action. Indirubin compounds, including indigo, have been also suggested as potential inhibitors of SARS coronavirus M^pro^ (Khalifa et al., 2021; Lin et al., 2005). Therefore, the inclusion of meisoindigo in our assays is consistent with existing knowledge regarding the potential anti-SARS coronavirus activity of indirubin compounds. The impressive results observed with meisoindigo further support its potential as an M^pro^ inhibitor and warrant further investigation for its therapeutic applications, particularly in the context of COVID-19.

### 2.3. *In vitro* M^pro^ inhibition

The results of our *in vivo* tests were however not confirmed through *in vitro* studies of M^pro^ inhibition with meisoindigo. The enzymatic activity of purified M^pro^ protein was studied. As shown in Fig 5B, meisoindigo did not show significant, dose-dependent inhibitory activity toward isolated M^pro^. The effect of GC376 on enzymatic activity of M^pro^ *in vitro* is also shown in Fig 5B, with IC_50_ value of 0.040 µM. This suggests that either meisoindigo interacts with the FRET assay used to measure M^pro^ activity or the inhibition of M^pro^ by this substance occurs *in vivo* but not *in vitro*. Additionally, the observation highlights the important advantage of our screening system. The yeast system facilities detection of the potentially effective M^pro^ inhibitors which have been discounted in large *in vitro* screenings as they interact with commonly used FRET assays. Thus our data still suggest that meisoindigo may be a strong candidate M^pro^ inhibitor in a biologically relevant context.

### 2.4. EGFP fluorescence assay

As mentioned above, the primary reason for expressing M^pro^ as a fusion protein with EGFP at the N-terminus was to ensure that M^pro^ is active and would autocatalytically cleave upon recognition of the linker sequence (SAVLQ). Additionally, this approach enabled us to monitor gene expression in real time by following green fluorescence. This method provides immediate visual feedback on the activity of the *GAL1* promoter.

Based on the difference observed in the growth of the EGFP-SAVLQ-Mpro and EGFP-D4K- Mpro strains (Fig 2C), which was correlated with a significant difference in EGFP fluorescence levels (Fig 2B and 6A), we decided to investigate whether fluorescence could serve as an additional bioindicator of the efficacy of M^pro^ inhibitors. During our experiment, the fluorescence intensity of yeast culture in the presence of different drugs was evaluated. Within just 3 hours of drug exposure, the fluorescence was significantly increased by the studied inhibitors, with the respective recorded values of 4.66 ± 0.13 x 10^4^ (10 µM ixazomib), 4.89 ± 0,19 x 10^4^ (1 µM meisoindigo), 5.02 ± 0.12 x 10^4^ (500 µM GC376), and· 4.49 ± 0.05 x 10^4^ (100 µM nirmatrelvir), compared to 4.34 ± 0.12 x 10^4^ for the control (DMSO) (Fig 6B). By the 24-hour mark, the differences were even more pronounced, reaching 2.79 ± 0,04 x 10^4^ (1 µM meisoindigo) and 5.17 ± 0.19 x 10^4^ (500 µM GC376) in comparison to 1.79 ± 0.03 x 10^4^ for the control (DMSO) (Fig 6C).

**Fig 6.**
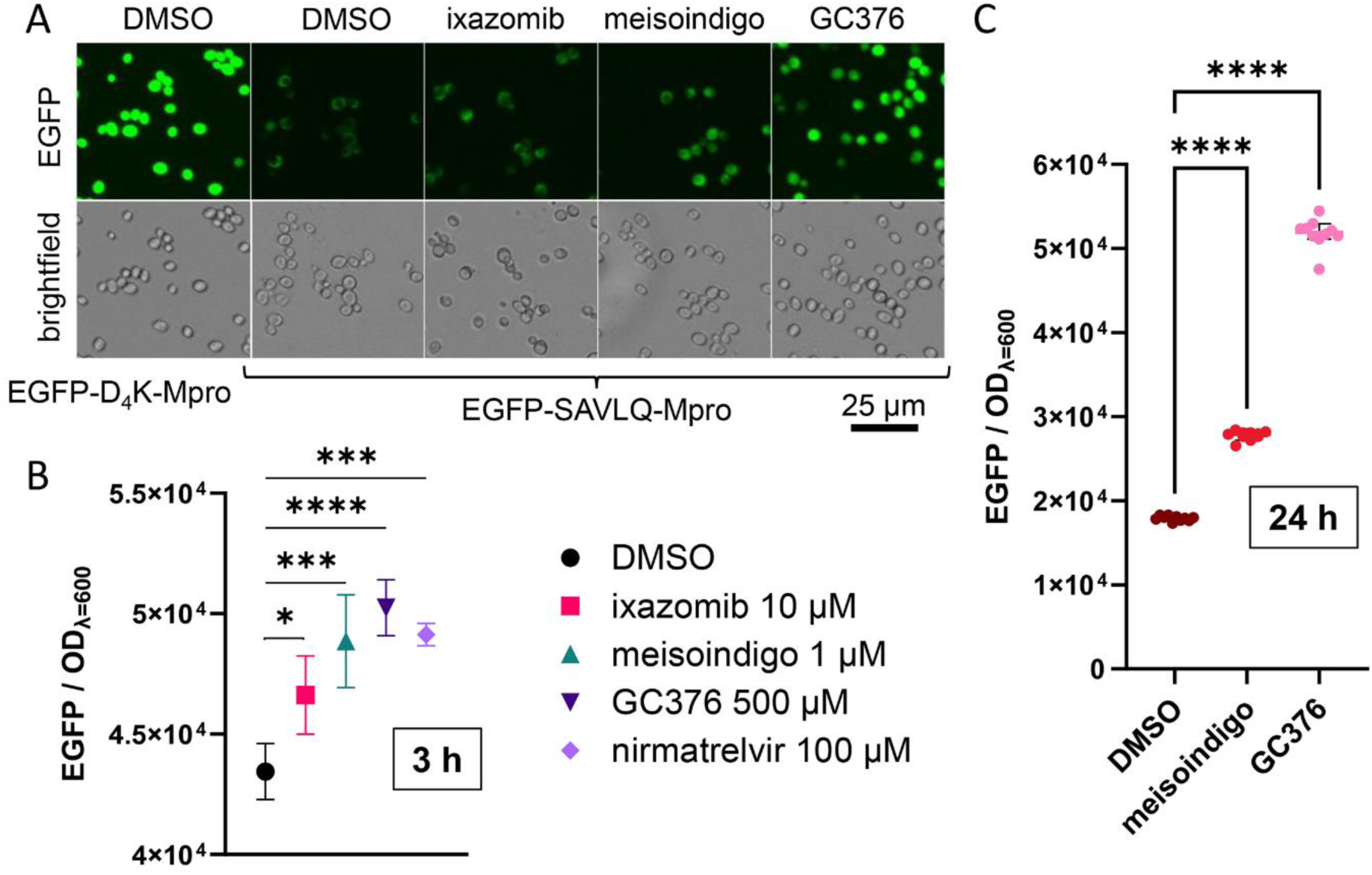
Quantitative analysis of green fluorescence of EGFP-SAVLQ-Mpro cells. **(A)** Microscopy images of EGFP-SAVLQ-Mpro yeast cells treated with various drugs, including an additional control of the EGFP-D4K-Mpro strain. **(B)** Quantitative analysis of green fluorescence levels in EGFP-SAVLQ-Mpro cells 3 hours after induction, with (*) indicating p < 0.05, (***) indicating p < 0.001, and (****) indicating p < 0.0001 based on n = 3 measurements. **(C)** The fluorescence levels 24 hours post-induction, with (****) indicating a p value less than 0.0001 for n = 10 measurements. Across all sections, the drug concentrations are consistent and are indicated in B. Error bars represent standard deviation (SD), and images were captured using the ZOE Fluorescent Cell Imager (Bio-Rad).

The use of GFP as a biosensor for cytotoxic compounds has been previously recognized. Transgenic *Leishmania infantum* promastigotes that continuously express GFP were used to monitor the effects of antileishmanial compounds (Kamau et al., 2001). The GFP-based assay served as a reliable measure of the inhibitory effects on protein expression, providing a dynamic representation of the response of leishmanial promastigotes to tested compounds. Another example of these assays is the GreenScreen genotoxicity assay, which simultaneously measures toxicity and genotoxicity in yeast (Cahill et al., 2004). The first description of the use of GFP-expressing mammalian cells for cytotoxic screening involved the use of the inducible Tet-On system to drive the expression of EGFP in HeLa cells to assess the cytotoxicity of cisplatin (Sandman et al., 1999). A strong correlation between the decline in GFP fluorescence and cytotoxicity was observed in HeLa cells treated with cisplatin and other platinum complexes.

Similarly, in our system, the decrease in EGFP fluorescence intensity due to the presence of active M^pro^ serves as an indicator of the cytotoxicity of the expressed enzyme. However, as shown in Fig S5, the fluorescence level can also be reduced due to the general cytotoxicity of the test drug.

Consequently, we found that meisoindigo, an anticancer drug, could exhibit toxicity at higher concentrations (especially those above 3 µM), leading to a decrease in the fluorescence of the fluorescent protein (EGFP in fusion with inactive M^pro^ or mCherry alone). Furthermore, the toxicity of meisoindigo reduced yeast growth and was strongly dependent on the genetic background. Strains with *pdr5*, *pdr1*, and *snq2* deletions were understandably more sensitive to 3 µM meisoindigo than the *pdr5*Δ variant (Fig S6). Interestingly, the toxicity was not associated with the *GAL1* promoter, as it also occurred on YPD, causing *GAL1* repression.

## 3. Materials & Methods

### 3.1. Tested compounds

MedChemExpress: FDA-Approved Drug Library Mini (Catalog # HY-L022M). Selleckchem: GC376 (Catalog # S0475). Merck: Tolperisone hydrochloride (Catalog # T3577); Ebselen (Catalog # E3520); Disulfiram (Catalog # PHR1690); Tideglusib (Catalog # SML0339); Carmofur (Catalog # C1494). WuhanChemNorm Biotech: Nirmatrelvir (Catalog # TBW03242).

### 3.2. Construction of plasmids

All oligonucleotides are shown in Table S1 (Supplementary file 1). The vectors for CRISPR/Cas9 were based on the pML104 vector (a gift from John Wyrick; Addgene plasmid # 676380), and the guide sequences were introduced by site-directed mutagenesis: the pML104-GAL1 vector has a guide sequence 5’-CTCTTAAATTATAGTTGGTT-3’ introduced by the Q5® Site-Directed Mutagenesis Kit (New England Biolabs) (Hu et al., 2018), while the pML104-PDR1 vector has a guide sequence, 5’-CTGGATAAACGTCGCTCCAC-3’, introduced by Q5 polymerase PCR (New England Biolabs) and In-Fusion Snap Assembly (Takara) site-directed mutagenesis as described by the manufacturer. A codon-optimized gene encoding the EGFP-M^pro^ fusion protein with a linker encoding a cleavage sequence recognized by M^pro^ (SAVLQ) flanked by sequences downstream (249 nt) and upstream (253 nt) of *GAL1* was synthesized in the pBSK(+) Simple-Amp vector by Biomatik (Ontario, Canada). The amino acid sequence of M^pro^ was derived from the ORF1a polyprotein (GenBank: UCQ02319.1). The pBSK(+)-Mpro(SAVLQ) vector (Fig S1) was used as a template for an enterokinase- recognized linker variant (D_4_K) made by using In-Fusion Snap Assembly (Takara) site-directed mutagenesis. The *mCherry* gene, sourced from the pMitoLoc template vector (Addgene # 58980), was integrated into the pBSK(+)-Mpro(SAVLQ) vector. Supplementary file 1 contains the DNA sequences.

### 3.3. Bacterial and yeast cultures

Bacteria were cultured in LB liquid medium (1% tryptone, 0.5% yeast extract, 1% sodium chloride) with 100 μg/ml ampicillin. Yeast cells were grown in YPD (1% yeast extract, 2% peptone, 2% D-glucose), supplemented with 300 μg/ml hygromycin B or 200 µg/ml G418 when necessary: YPG (1% yeast extract, 2% peptone, 3% glycerol, pH = 5.5); YPGGal (1% yeast extract, 2% peptone, 3% glycerol, pH = 5.5 and 0.5% galactose); YPSuc (1% yeast extract, 2% peptone, 2% sucrose); YPSucGal (1% yeast extract, 2% peptone, 2% sucrose, 0.5% galactose); SD-ura (0.67% yeast nitrogen base without amino acids, yeast synthetic drop-out medium supplement without uracil, and 2% D-glucose) or SDC + 5-FOA (0.67% yeast nitrogen base without amino acids, yeast complete synthetic drop-out medium supplement, 1 mg/ml 5- fluoroorotic acid,2% D-glucose). Agar (2%) was added to the solid media.

### 3.4. Strains

NEB® 5-alpha competent *E. coli* was ordered from New England Biolabs (Catalog # C2987H). Yeast *S. cerevisiae* strains were derived from BY4741 and are listed in Table S2 (Supplementary file 1). The Δgal1 strain was generated by CRISPR/Cas9 using pML104- GAL1, and DNA repair was performed by hybridizing two complementary oligonucleotides containing a sequence below (35 nt) and above (35 nt) the *GAL1* open reading frame. EGFP- SAVLQ-M^pro^, EGFP-D4K-M^pro^, and mCherry strains were obtained by CRISPR/Cas9 using repair DNA produced by PCR. The *PDR5* gene was deleted via a KanMX6 deletion cassette. The *PDR1* gene was deleted by CRISPR/Cas9 using the KanMX6 cassette as repair DNA. The *SNQ2* gene was deleted via the HphMX6 cassette. The procedure of yeast transformation was according to the user manual of Yeastmaker™ Yeast Transformation System 2 (Takara) with small scale transformation. The transformed cells were grown on SD-ura medium at 28°C.

Each application of CRISPR/Cas9 involved removal of the pML104 vector from yeast cells by culturing in SDC + 5-FOA medium (Laughery and Wyrick, 2019).

### 3.5. Yeast-based drug screening assay

After overnight culture on YPG medium, the yeast were centrifuged and adjusted to OD_600_ = 10. The suspension (0.8 ml) was spread homogeneously with stainless steel beads (3 mm diameter) on a Petri dish (140 mm diameter) containing YPSucGal solid medium. Individual compounds from the chemical library were prepared at a concentration of 10 mM and applied in a volume of 3 μl to a sterile filter (6 mm diameter). A filter with DMSO was used as a vehicle control. Filters were placed on the agar surface at a 10 mm distance (64 filters per plate). The plates were incubated at 28 °C for 48 h and scanned.

The criteria for evaluating the compounds focused on identifying those that promoted increased yeast growth, indicating a positive hit. Compounds were selected based on their ability to significantly enhance yeast colony size or number compared to the DMSO control. Preference was given to compounds displaying symmetrical growth enhancement around the filter, similar to the reference inhibitor GC376. Compounds with potential neighboring drug effects or inconsistent results (e.g., no effect at 24 hours but strong effect at 48 hours) were excluded.

### 3.6. Growth and fluorescence assay on liquid medium

The yeast were cultured in a flask on YPG liquid medium. The culture was then inoculated into 1 ml YPSucGal medium in 24-well plates, at OD_600_ 0,05. A drug or DMSO as a vehicle was added to individual wells. The final concentration of DMSO in each well was 1%. The culture plate was incubated using a microplate shaker (PSU-2T, BioSan) at 28°C, 700 rpm. After 24h the OD_600_ was measured.

For fluorescence intensity, 100 μl of the undiluted culture was transferred to a 96-well flat bottom plates. EGFP (ex. 485 nm, em. 532 nm) and mCherry (ex. 580 nm, em. 620 nm) fluorescence was measured with The Spark® multimode microplate reader (TECAN) at RT.

### 3.7. Calculation of the Z’ Factor

The calculation was based on the means of OD_600_ readings from liquid cultures after 24 hours of induction. The Z’ factor was calculated using the formula:

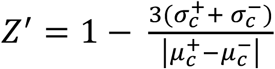

where σ^+^_*c*_ and σ^−^_*c*_ are the standard deviations, and μ^+^_*c*_ and μ^−^*c* are the means of the positive and negative controls, respectively. The positive control involved yeast cells expressing M^pro^ without inhibitors, manifesting inhibited growth due to the enzyme’s toxicity; the negative control involved yeast cells without M^pro^ expression (non-induced), showing normal growth.

### 3.8. Measurement of M^pro^ inhibition

The inhibition of SARS-CoV-2 Mpro in vitro enzymatic activity was measured using 3CL Protease, Untagged (SARS-CoV-2) Assay Kit (BPS Bioscience), according to the manufacturer’s instructions. Briefly, tested compounds were diluted in DMSO at 100x the desired concentration. Further dilutions were made in Assay buffer. The concentration of DMSO used in a reaction did not exceed 1%. Mpro (15 ng per reaction) diluted in assay buffer (containing 1 mM DTT) was introduced into wells containing tested compounds or DMSO. The enzyme with tested compounds was incubated 30 min at RT with slow shaking. The reaction was initiated by the addition of 40 µM the 3CL Protease substrate solution to each well. The sealed plate was incubated for 4 h at RT with slow shaking. Finally, the fluorescence intensity was measured using The Spark® multimode microplate reader (TECAN) at an excitation wavelength of 360 nm and detection of emission at a wavelength of 460 nm.

### 3.9. Statistics

Statistical analysis was performed using GraphPad Prism software v.9.5.1 (GraphPad Software, San Diego, CA, USA). Data are presented as the mean ± standard deviation (SD) from at least three independent experiments. Statistical significance was assessed by ANOVA for three or more groups or Welch’s t test for two groups. P values less than 0.05 were considered statistically significant.

## 4. Conclusions

A yeast system characterized by growth restriction induced by M^pro^ expression was developed and used to evaluate the effects of various compounds, including potential M^pro^ inhibitors. An initial screen using the known M^pro^ inhibitor GC376 validated the system’s ability to identify M^pro^ inhibitors. A comprehensive search of a drug library identified several candidates with potential inhibitory effects on M^pro^, including ixazomib and ethacridine, which had been suggested as potential M^pro^ inhibitors in previous studies. However, these drugs were found to be ineffective in subsequent liquid medium assays. The study also identified several drugs not previously associated with M^pro^ inhibition, including meisoindigo, a derivative of indigo, which showed significant inhibitory activity against Mpro in *in vivo* assays but not *in vitro*. This discrepancy suggests that some drugs might interfere with biochemical methods and highlights the advantage of the developed bioassay.

In this study, EGFP fusion was used to track M^pro^ protein activity and gene expression, serving as a potential indicator for M^pro^ inhibitor efficacy. The decrease in EGFP intensity signaled the cytotoxicity of the active M^pro^ enzyme, although it could also reflect the cytotoxicity of the test drug. The observed increase in fluorescence intensity was associated with the inhibition of M^pro^ activity while ensuring that the drug concentration did not induce cytotoxicity to the cell.

This makes the developed system highly valuable for screening potential inhibitors, both at the level of the survival assay and by a highly sensitive method based on EGFP fluorescence. Using these methods, meisoindigo was identified as a potent M^pro^ inhibitor. The discovery of this new compound with potential M^pro^ inhibitory activity could open new avenues of research, including the creation of new derivatives that could potentially serve as drugs for SARS-CoV- 2 and other coronavirus infections.

