## Supplementary File 1 for "Meisoindigo: An Effective Inhibitor of SARS-CoV-2 Main Protease Revealed by Yeast System"

**Title**: Meisoindigo: An Effective Inhibitor of SARS-CoV-2 Main Protease Revealed by Yeast System

**Authors**: Wojciech Grabiński, Anna Kicińska, Ewa Kosicka, Martyna Baranek-Grabińska, Ewelina Hejenkowska, Joanna Budzik, Paulina Śliska, Weronika Śliwińska, Andonis Karachitos*

**Description**: This file contains 6 figures and 2 tables and DNA sequences that supplement the content of the main article.

A

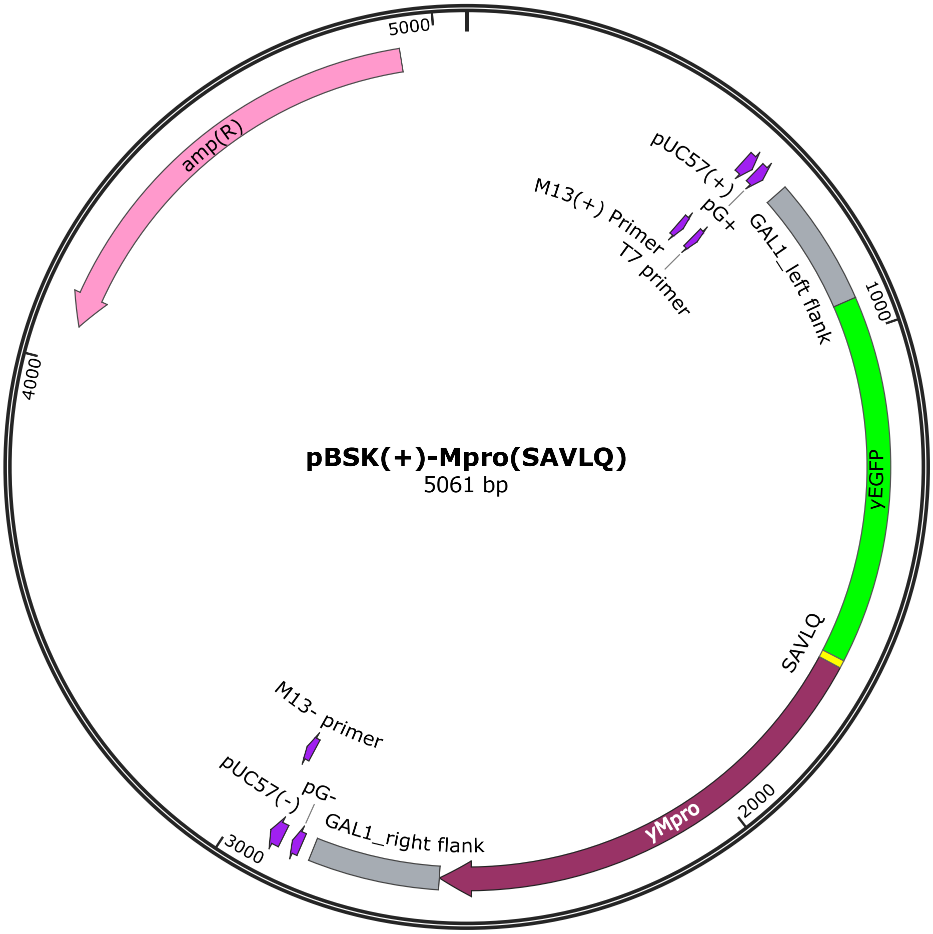


B

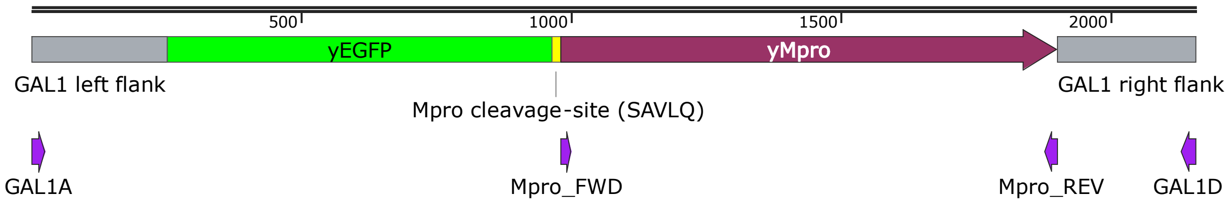


**Fig S1. (A)** Map showing the vector pBSK(+)-Mpro(SAVLQ). **(B)** Linear DNA repair template with the location of primers used in the work (purple arrows). Flanking sequences of *GAL1* are shown in gray. The figure was created using SnapGene® software.

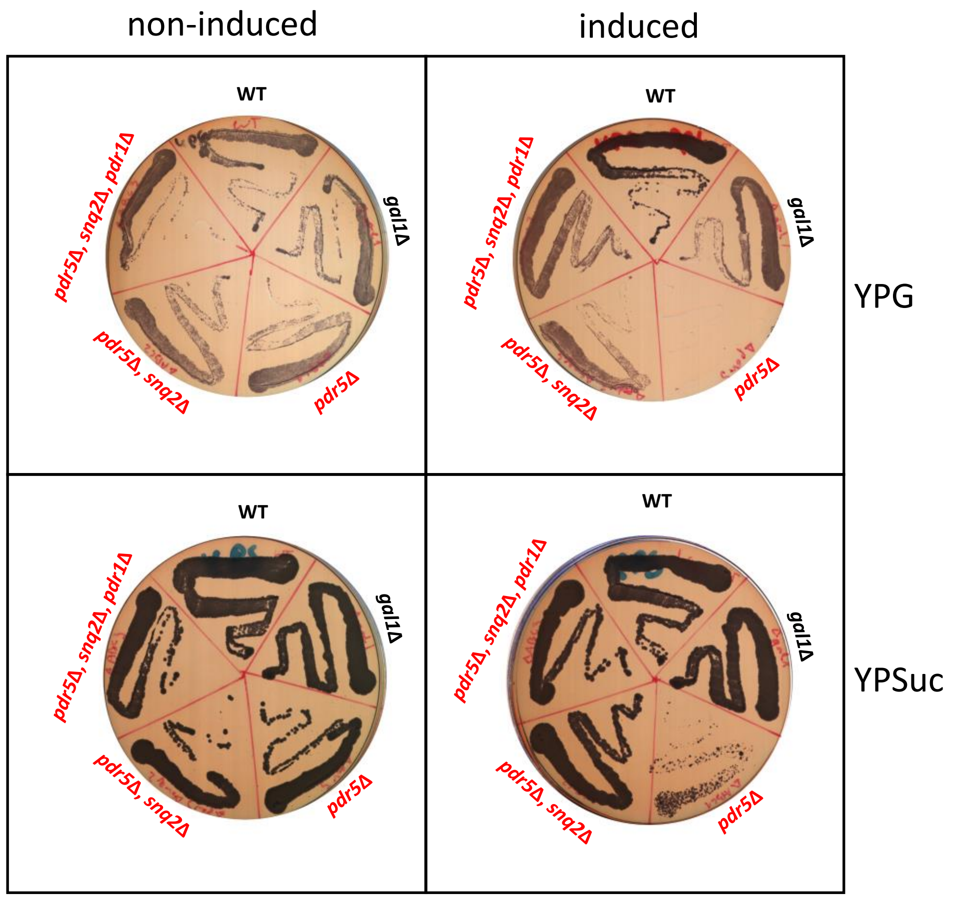


**Fig S2.** Effect of deletion of gene(s) (PDR1, PDR5 and/or SNQ2) responsible for multidrug resistance on toxicity induced by Mpro expression in EGFP-SAVLQ-Mpro strain (labeled in red), compared to a wild-type strain (WT) and a strain lacking the *GAL1* gene (*gal1*Δ). Yeast was transferred from liquid culture and spread on solid medium containing glycerol (YPG) or sucrose (YPSuc) as a carbon source. Mpro expression was induced by galactose supply.

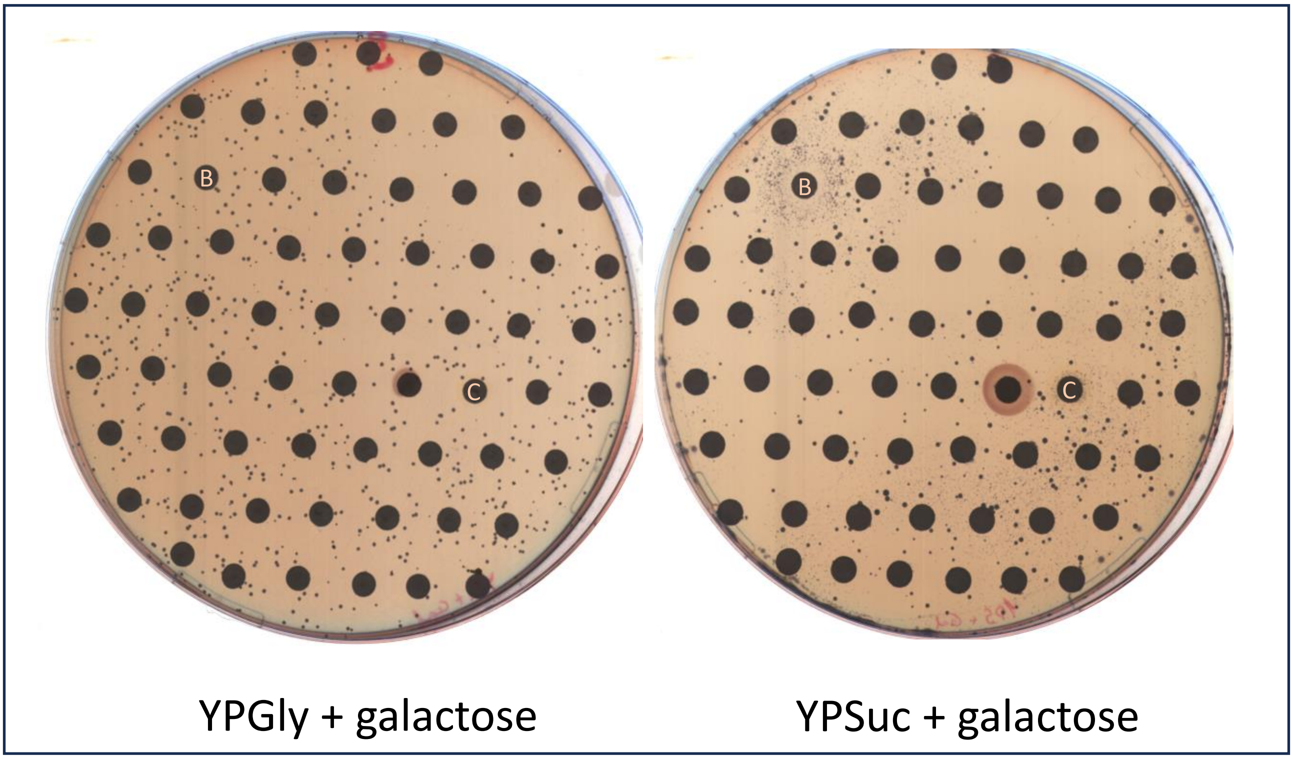


**Fig S3.** The difference in growth of EGFP-SAVLQ-Mpro yeast during screening, depending on the medium used: YPG containing glycerol as a carbon source or YPSuc containing sucrose. Both media include the inducing agent galactose. Drug filters are spread on both media according to the same pattern. Filters marked with a letter correspond to drugs that gave a visible growth effect (halo) during screening on YPSuc medium, where (B) is berberine and (C) is ciclopirox.

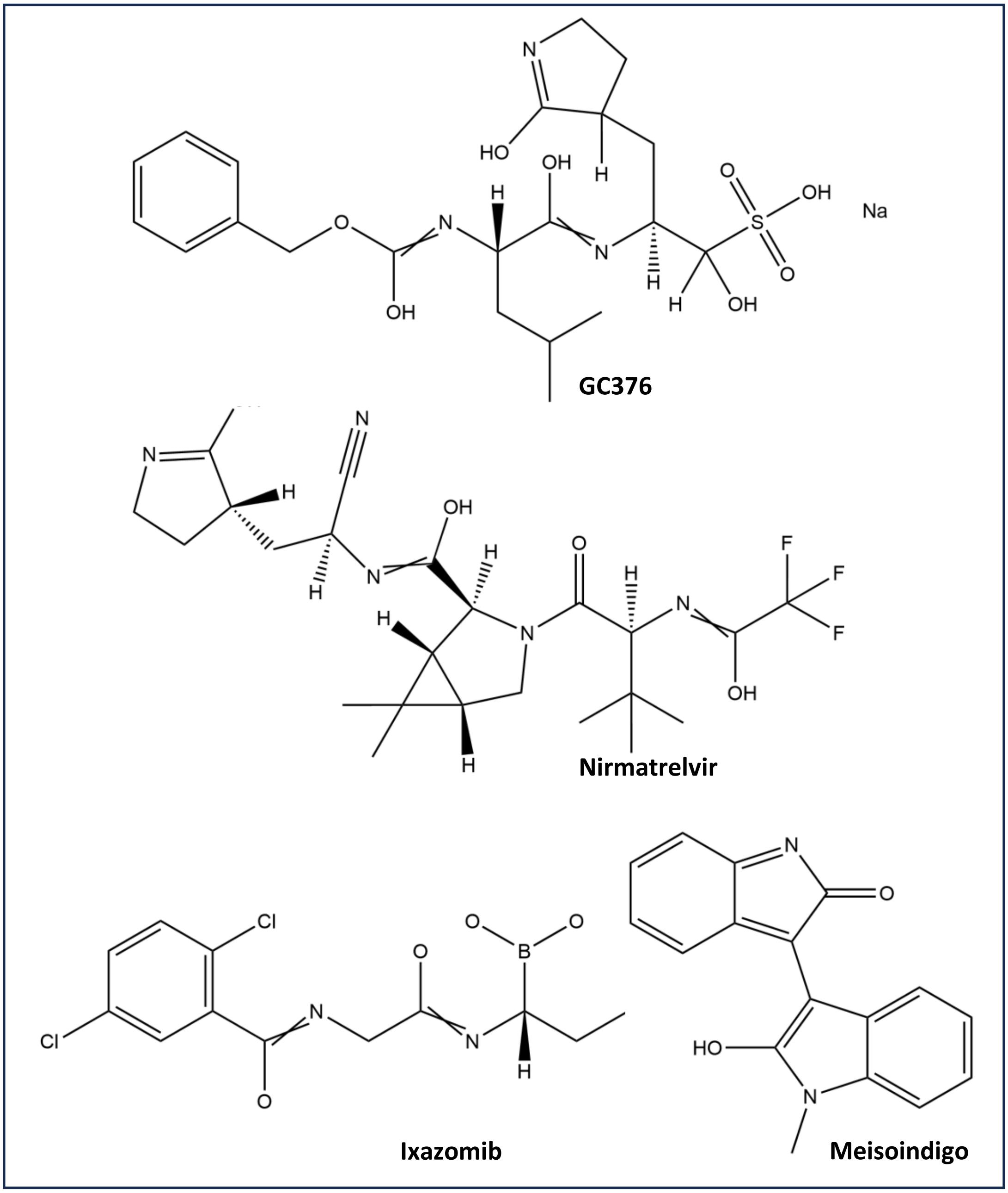


**Fig S4.** Chemical structures of the drugs used in the present study of Mpro activity inhibition.

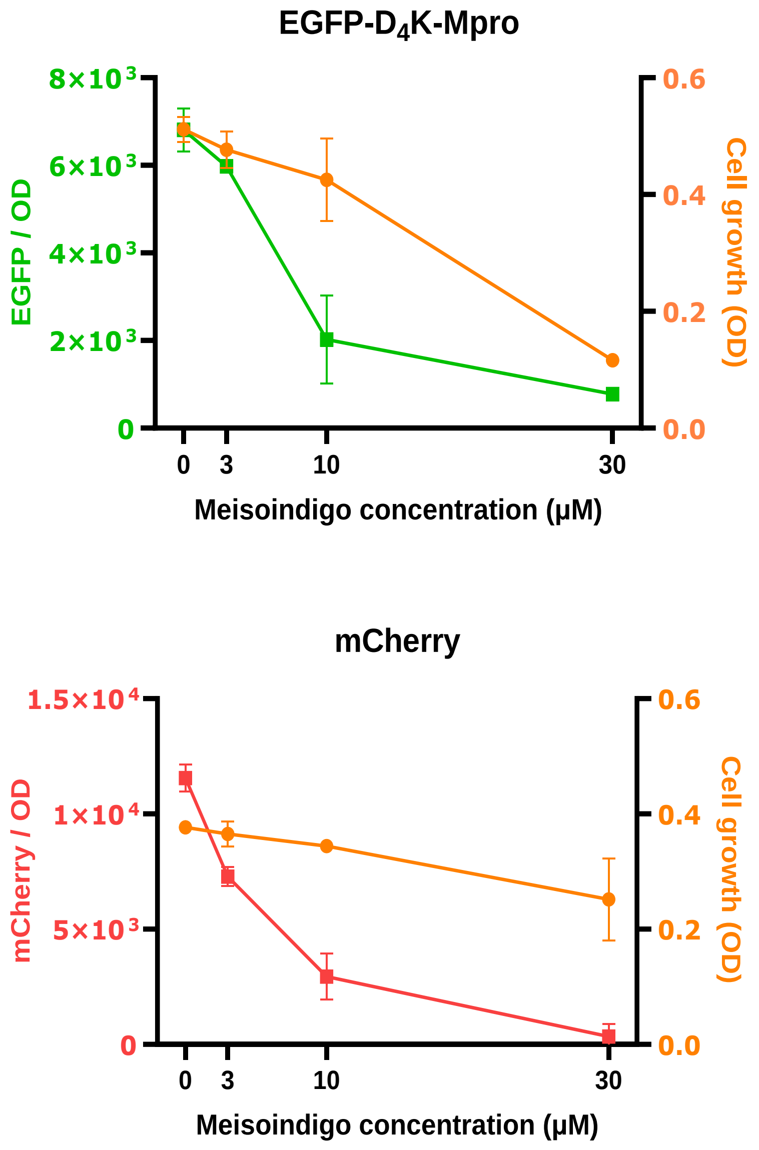


**Fig S5.** Meisoindigo concentration-dependent changes in growth (orange line) and EGFP fluorescence of the EGFP-D4K-Mpro strain (green line) and the mCherry strain (red line). Error bars indicate SD (n = 3).

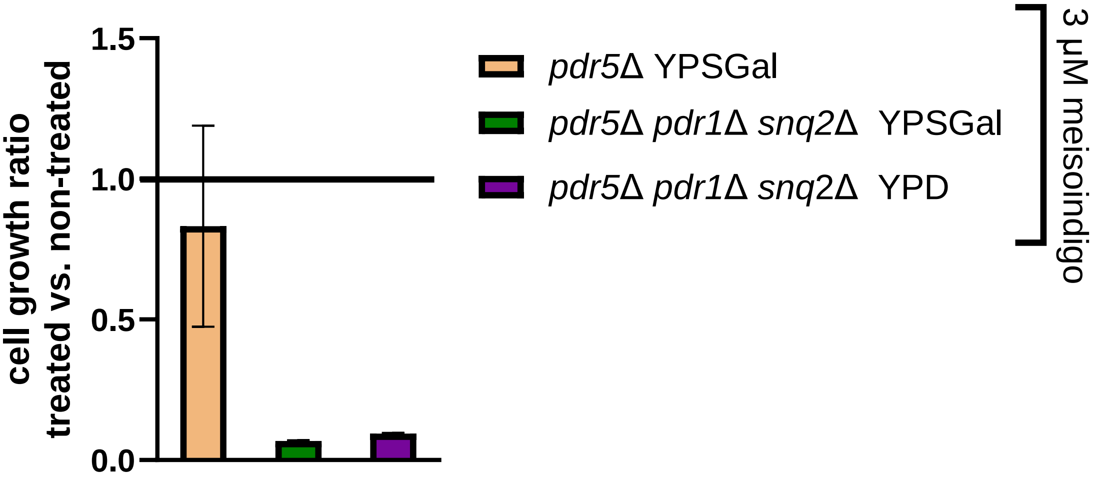


**Fig S6.** Effects of meisoindigo on yeast growth depending on the type of multidrug resistance gene deletion. The growth parameter was expressed as the ratio of OD values of cells treated with 3 μM meisoindigo to control cells to which DMSO was added as a vehicle. Error bars indicate SD (n = 3).

**Table S1.** List of oligonucleotides used in this work.

| **Name** | **Sequence 5’ -> 3’** | **Use and specification** |
|---|---|---|
| GAL1A | ACGAATCAAATTAACAACCATAGGA | amplification of repair DNA, sequencing |
| GAL1D | ATGTCAAGAATAGGTATCCAAAACG | |
| GAL1_gRNa_1 | GTTTTAGAGCTAGAAATAGCAAGTTAAAATAAGGC | insertion of guide sgRNA sequence into pML104 vector |
| GAL1_gRNa_2 | AACCAACTATAATTTAAGAGGATCATTTATCTTTCACTGC | |
| GAL1ko1 | ATATACCTCTATACTTTAACGTCAAGGAGAAAAAACTATAGTATACTTCTTTTTTTTACTTTGTTCAGAACAACTTCTCA | oligonucleotides for hybridization, DNA repair for gene deletion |
| GAL1ko2 | TGAGAAGTTGTTCTGAACAAAGTAAAAAAAAGAAGTATACTATAGTTTTTTCTCCTTGACGTTAAAGTATAGAGGTATAT | |
| Mprofd | TCTGGTTTCAGAAAGATGGCT | sequencing |
| Mprorv | TTATTGGAAAGTAACACCAGAACA | |
| MproD4Kfd | AGACGACGACGACAAGTCTGGTTTCAGAAAGATGGCTTTCC | insertion of the sequence encoding the D4K fragment |
| MproD4Krv | TTGTCGTCGTCGTCTTTGTACAATTCATCCATACCATGG | |
| PDR1_gRNa_1 | ACGTCGCTCCACGTTTTAGAGCTAGAAATAGCAAGTTAAAATAAGGCTAG | insertion of guide sgRNA sequence into pML104 vector |
| PDR1_gRNa_2 | AACGTGGAGCGACGTTTATCCAGGATCATTTATCTTTCACTGCGGAG | |
| PDR1_A | ACCCTAAATGGAGTTTCTTTTCTTG | amplification of repair DNA, sequencing, diagnostic |
| PDR1_D | CCCATAAGAAATACACCTCATGGTA | |
| PDR1_B | CACTTCCGCTATTATCACCACTACT | diagnostic |
| PDR5_A | TTGAACGTAATCTGAGCAATACAAA | amplification of repair DNA, sequencing, diagnostic |
| PDR5_D | TCACACTAAATGCTGATGCCTATAA | |
| PDR5_B | TACCTAAAACGACTAGCAATTCACC | diagnostic |

| | | |
|---|---|---|
| SNQ2_A | CACCACTTTTTATGCTTGTATATGCT | amplification of repair DNA, sequencing, diagnostic |
| SNQ2_D | GTTTAGCTTGTACTTGCAACACCTT | |
| SNQ2_B | GTCCTGTCAGCATTTTTGTTAGTTT | diagnostic |
| pBSK-GAL1fd | GTATACTTCTTTTTTTTACTTTGT | vector linearization by PCR |
| pBSK-GAL1rv | TATAGTTTTTTCTCCTTGACGT | |
| mCherryfd | GGAGAAAAAACTATAATGGTGAGCAAGGGCGAGG | amplification of the mCherry gene insert for cloning |
| mCherryrv | AAAAAAGAAGTATACTTACTTGTACAGCTCGTCCATGCC | |

**Table S2.** List of *Saccharomyces cerevisiae* strains used in this work.

| **Strain** | **Genotype** | **Source** |
|---|---|---|
| BY4741 | *MATa his3*Δ1 *leu2*Δ0 *met15*Δ0 *ura3*Δ0 | Euroscarf |
| Δgal1 | *MATa his3*Δ1 *leu2*Δ0 *met15*Δ0 *ura3*Δ0 *gal1*Δ0 | This work |
| Δpdr5 | *MATa his3*Δ1 *leu2*Δ0 *met15*Δ0 *ura3*Δ0 *pdr5*Δ0::KanMX6 | This work |
| Δpdr5, Δpdr1, Δsnq2 | *MATa his3*Δ1 *leu2*Δ0 *met15*Δ0 *ura3*Δ0 *pdr5*Δ0::KanMX6 *pdr1*Δ0::KanMX6 *Δsnq2*Δ0::hphMX6 | This work |
| EGFP-SAVLQ-Mpro | *MATa his3*Δ1 *leu2*Δ0 *met15*Δ0 *ura3*Δ0 *pdr5*Δ0::KanMX6 *gal1*Δ0::yEGFP$_{(SAVLQ)}$yMpro | This work |
| EGFP-SAVLQ-Mpro Δpdr5, Δsnq2 | *MATa his3*Δ1 *leu2*Δ0 *met15*Δ0 *ura3*Δ0 *pdr5*Δ0::KanMX6 *Δsnq2*Δ0::hphMX6 *gal1*Δ0::yEGFP$_{(SAVLQ)}$yMpro | This work |
| EGFP-SAVLQ-Mpro Δpdr5, Δsnq2, Δpdr1 | *MATa his3*Δ1 *leu2*Δ0 *met15*Δ0 *ura3*Δ0 *pdr5*Δ0::KanMX6 *pdr1*Δ0::KanMX6 *Δsnq2*Δ0::hphMX6 *gal1*Δ0::yEGFP$_{(SAVLQ)}$yMpro | This work |
| EGFP-D$_4$K-Mpro | *MATa his3*Δ1 *leu2*Δ0 *met15*Δ0 *ura3*Δ0 *pdr5*Δ0::KanMX6 *gal1*Δ0::yEGFP$_{(DDDDK)}$yMpro | This work |
| mCherry | *MATa his3*Δ1 *leu2*Δ0 *met15*Δ0 *ura3*Δ0 *pdr5*Δ0::KanMX6 *gal1*Δ0::mCherry | This work |

### <u>DNA sequences</u>

*PDR5* deletion cassette – *pdr5*Δ::*kanMX6*

TTGAACGTAATCTGAGCAATACAAACAAGGCCTCTCCTATACATATATAATTGTGATGTGCATAACCTTATGGCT
GTTCGCTTTTATTATCATACCTTAGAATGAAATCCAAAAGAAAAAAGTCACGCAAAGTTGCAAACATATAACAAC
TGTGTTAGTTATCACTCGACTTTGTTATTCTAATTATAAATAAATTGGCAACTAGGAACTTTCGAAAAAGAAATT
AAAGACCCTTTTAAGTTTTCGTATCCGCTCGTTCGAAAGACTTTAGACAAAAGTTTAGCTTGCCTCGTCCCCGCC
GGGTCACCCGGCCAGCGACATGGAGGCCCAGAATACCCTCCTTGACAGTCTTGACGTGCGCAGCTCAGGGGCATG
ATGTGACTGTCGCCCGTACATTTAGCCCATACATCCCCATGTATAATCATTTGCATCCATACATTTTGATGGCCG
CACGGCGCGAAGCAAAAATTACGGCTCCTCGCTGCAGACCTGCGAGCAGGGAAACGCTCCCCTCACAGACGCGTT
GAATTGTCCCCACGCCGCGCCCCTGTAGAGAAATATAAAAGGTTAGGATTTGCCACTGAGGTTCTTCTTTCATAT
ACTTCCTTTTAAAATCTTGCTAGGATACAGTTCTCACATCACATCCGAACATAAACAACCATGGGTAAGGAAAAG
ACTCACGTTTCGAGGCCGCGATTAAATTCCAACATGGATGCTGATTTATATGGGTATAAATGGGCTCGCGATAAT
GTCGGGCAATCAGGTGCGACAATCTATCGATTGTATGGGAAGCCCGATGCGCCAGAGTTGTTTCTGAAACATGGC
AAAGGTAGCGTTGCCAATGATGTTACAGATGAGATGGTCAGACTAAACTGGCTGACGGAATTTATGCCTCTTCCG
ACCATCAAGCATTTTATCCGTACTCCTGATGATGCATGGTTACTCACCACTGCGATCCCCGGCAAAACAGCATTC
CAGGTATTAGAAGAATATCCTGATTCAGGTGAAAATATTGTTGATGCGCTGGCAGTGTTCCTGCGCCGGTTGCAT
TCGATTCCTGTTTGTAATTGTCCTTTTAACAGCGATCGCGTATTTCGTCTCGCTCAGGCGCAATCACGAATGAAT
AACGGTTTGGTTGATGCGAGTGATTTTGATGACGAGCGTAATGGCTGGCCTGTTGAACAAGTCTGGAAAGAAATG
CATAAGCTTTTGCCATTCTCACCGGATTCAGTCGTCACTCATGGTGATTTCTCACTTGATAACCTTATTTTTGAC
GAGGGGAAATTAATAGGTTGTATTGATGTTGGACGAGTCGGAATCGCAGACCGATACCAGGATCTTGCCATCCTA
TGGAACTGCCTCGGTGAGTTTTCTCCTTCATTACAGAAACGGCTTTTTCAAAAATATGGTATTGATAATCCTGAT
ATGAATAAATTGCAGTTTCATTTGATGCTCGATGAGTTTTTCTAATCAGTACTGACAATAAAAAGATTCTTGTTT
TCAAGAACTTGTCATTTGTATAGTTTTTTTATATTGTAGTTGTTCTATTTTAATCAAATGTTAGCGTGATTTATA
TTTTTTTTCGCCTCGACATCATCTGCCCAGATGCGAAGTTAAGTGCGCAGAAAGTAATATCATGCGTCAATCGTA
TGTGAATGCTGGTCGCTATACTGCTGTCGATTCGATACTAACGCCGCCATCCAGTTTTAGAATTTTGAATTTGGT
TAAGAAAAGAAACTTACCAAGATGGACTTTTTTAATAAACATACATAATCACTACATATAGGTGCGTAATAATAA
GTTTTTATTTTTTTTTTCTTAATTCAGCGAGCTTTCTACATTTCAATTTTCTTGAATTTACCAGCTCTGATAAAT
CAAAGTTCAATGTCCGAAAGAATTCCGAGAATTTTATTCCTAGGTTATCTCATCGGTACTTTTTTTTTATAGGCA
TCAGCATTTAGTGTGA

nucleotides homologous to the flanking region of *PDR5* gene

kanMX6 cassette (G418 resistance)

a primer binding site

PDR3 deletion cassette/repair DNA - *pdr3Δ::kanMX6* (G418 resistance)

TTCACTTTTCATTTTCTTCCAGAACAACAGGCGCTGCCCTTGTTTCCGCGGAGCTGCCTCCTCTGCCGCTCGGAC
TTTCCGCGGAATAATAAATGAACTATCACAGTGAGTCTTAGAATGCAAATATGTAGATACTGTTATAGAATCTCA
TTAATGTATTTATGTATTTCCTGTGTTTGTGTCTTCAGGAGCCTTCCAAAGCAACCATAGGTCTTAGGCGACAAC
TGCATCAGCAGTTTTATTAATTTTTTCTTATTGCGTGACCGCAGTTTAGCTTGCCTCGTCCCCGCCGGGTCACCC
GGCCAGCGACATGGAGGCCCAGAATACCCTCCTTGACAGTCTTGACGTGCGCAGCTCAGGGGCATGATGTGACTG
TCGCCCGTACATTTAGCCCATACATCCCCATGTATAATCATTTGCATCCATACATTTTGATGGCCGCACGGCGCG
AAGCAAAAATTACGGCTCCTCGCTGCAGACCTGCGAGCAGGGAAACGCTCCCCTCACAGACGCGTTGAATTGTCC
CCACGCCGCGCCCCTGTAGAGAAATATAAAAGGTTAGGATTTGCCACTGAGGTTCTTCTTTCATATACTTCCTTT
TAAAATCTTGCTAGGATACAGTTCTCACATCACATCCGAACATAAACAACCATGGGTAAGGAAAAGACTCACGTT
TCGAGGCCGCGATTAAATTCCAACATGGATGCTGATTTATATGGGTATAAATGGGCTCGCGATAATGTCGGGCAA
TCAGGTGCGACAATCTATCGATTGTATGGGAAGCCCGATGCGCCAGAGTTGTTTCTGAAACATGGCAAAGGTAGC
GTTGCCAATGATGTTACAGATGAGATGGTCAGACTAAACTGGCTGACGGAATTTATGCCTCTTCCGACCATCAAG
CATTTTATCCGTACTCCTGATGATGCATGGTTACTCACCACTGCGATCCCCGGCAAAACAGCATTCCAGGTATTA
GAAGAATATCCTGATTCAGGTGAAAATATTGTTGATGCGCTGGCAGTGTTCCTGCGCCGGTTGCATTCGATTCCT
GTTTGTAATTGTCCTTTTAACAGCGATCGCGTATTTCGTCTCGCTCAGGCGCAATCACGAATGAATAACGGTTTG
GTTGATGCGAGTGATTTTGATGACGAGCGTAATGGCTGGCCTGTTGAACAAGTCTGGAAAGAAATGCATAAGCTT
TTGCCATTCTCACCGGATTCAGTCGTCACTCATGGTGATTTCTCACTTGATAACCTTATTTTTGACGAGGGGAAA
TTAATAGGTTGTATTGATGTTGGACGAGTCGGAATCGCAGACCGATACCAGGATCTTGCCATCCTATGGAACTGC
CTCGGTGAGTTTTCTCCTTCATTACAGAAACGGCTTTTTCAAAAATATGGTATTGATAATCCTGATATGAATAAA
TTGCAGTTTCATTTGATGCTCGATGAGTTTTTCTAATCAGTACTGACAATAAAAAGATTCTTGTTTTCAAGAACT
TGTCATTTGTATAGTTTTTTTATATTGTAGTTGTTCTATTTTAATCAAATGTTAGCGTGATTTATATTTTTTTTC
GCCTCGACATCATCTGCCCAGATGCGAAGTTAAGTGCGCAGAAAGTAATATCATGCGTCAATCGTATGTGAATGC
TGGTCGCTATACTGCTGTCGATTCGATACTAACGCCGCCATCCAGTTTAAACGCAAAGAAAATAGGGAAGCAGAG
CATAACCATAGTAAATGGGACACTTATAACTCAAGAATTTAATTGACCCCCTACTCATACTGAGGAATGTGGCGT
CCTCGTCATTATATGTTCGTAAGCTAATTGGGCGAAACTTTAGTCTGTCCTTTCTGTTCGGTCTTCACATAAGAT
CAACGATTTCTTCATTAATTTTGGGCGCGCTGTTCAAAAAGGTCCTGGAATGCAGCTGTTCAGTAGATATT

nucleotides homologous to the flanking region of *PDR3* gene

kanMX6 cassette (G418 resistance)

a primer binding site

*SNQ2* deletion cassette - *snq2*Δ::*hphMX6* (hygromycin B resistance)

CACCACTTTTTATGCTTGTATATGCTTGTAGATAAATAAATACCTAACGAAAACAATATATACGAACCTAGTGTA
TTTGTATCTTTTGTTTTTCAAGTTGAAGTGTTGCGAGGTCAAAAAAAAAAAAGCTCACTGTAGGATATAGGAGTT
ACTATATCACCAGTGCATTACATTCTCAGTGCATCCATCGTCTTCAACATTGATTATTTTCTCCTTCCATTGATT
AGAGTTCAAGCTCCCTGAGAAAACGAAGGTATAGCGGACGTACCCGCAGAGACATAAAAAAAGAAAAACTATATC
GAAGACCGAAAGCAGTAAAAAAGTGGATAGAATAACACAGCTACCAAAATACGTAAAGAGAATTCAGACATGGAG
GCCCAGAATACCCTCCTTGACAGTCTTGACGTGCGCAGCTCAGGGGCATGATGTGACTGTCGCCCGTACATTTAG
CCCATACATCCCCATGTATAATCATTTGCATCCATACATTTTGATGGCCGCACGGCGCGAAGCAAAAATTACGGC
TCCTCGCTGCAGACCTGCGAGCAGGGAAACGCTCCCCTCACAGACGCGTTGAATTGTCCCCACGCCGCGCCCCTG
TAGAGAAATATAAAAGGTTAGGATTTGCCACTGAGGTTCTTCTTTCATATACTTCCTTTTAAAATCTTGCTAGGA
TACAGTTCTCACATCACATCCGAACATAAACAACCATGGGTAAAAAGCCTGAACTCACCGCGACGTCTGTCGAGA
AGTTTCTGATCGAAAAGTTCGACAGCGTCTCCGACCTGATGCAGCTCTCGGAGGGCGAAGAATCTCGTGCTTTCA
GCTTCGATGTAGGAGGGCGTGGATATGTCCTGCGGGTAAATAGCTGCGCCGATGGTTTCTACAAAGATCGTTATG
TTTATCGGCACTTTGCATCGGCCGCGCTCCCGATTCCGGAAGTGCTTGACATTGGGGAATTCAGCGAGAGCCTGA
CCTATTGCATCTCCCGCCGTGCACAGGGTGTCACGTTGCAAGACCTGCCTGAAACCGAACTGCCCGCTGTTCTGC
AGCCGGTCGCGGAGGCCATGGATGCGATCGCTGCGGCCGATCTTAGCCAGACGAGCGGGTTCGGCCCATTCGGAC
CGCAAGGAATCGGTCAATACACTACATGGCGTGATTTCATATGCGCGATTGCTGATCCCCATGTGTATCACTGGC
AAACTGTGATGGACGACACCGTCAGTGCGTCCGTCGCGCAGGCTCTCGATGAGCTGATGCTTTGGGCCGAGGACT
GCCCCGAAGTCCGGCACCTCGTGCACGCGGATTTCGGCTCCAACAATGTCCTGACGGACAATGGCCGCATAACAG
CGGTCATTGACTGGAGCGAGGCGATGTTCGGGGATTCCCAATACGAGGTCGCCAACATCTTCTTCTGGAGGCCGT
GGTTGGCTTGTATGGAGCAGCAGACGCGCTACTTCGAGCGGAGGCATCCGGAGCTTGCAGGATCGCCGCGGCTCC
GGGCGTATATGCTCCGCATTGGTCTTGACCAACTCTATCAGAGCTTGGTTGACGGCAATTTCGATGATGCAGCTT
GGGCGCAGGGTCGATGCGACGCAATCGTCCGATCCGGAGCCGGGACTGTCGGGCGTACACAAATCGCCCGCAGAA
GCGCGGCCGTCTGGACCGATGGCTGTGTAGAAGTACTCGCCGATAGTGGAAACCGACGCCCCAGCACTCGTCCGA
GGGCAAAGGAATAATCAGTACTGACAATAAAAAGATTCTTGTTTTCAAGAACTTGTCATTTGTATAGTTTTTTTA
TATTGTAGTTGTTCTATTTTAATCAAATGTTAGCGTGATTTATATTTTTTTTCGCCTCGACATCATCTGCCCAGA
TGCGAAGTTAAGTGCGCAGAAAGTAATATCATGCGTCAATCGTATGTGAATGCTGGTCGCTATACTGTGTGGGCT
TCAGATAACTTAACATTTTGTGCATTCATCTGCCTTTAATTCTTAATTCTCATTTTACACTATAGAAGAAGACAC
GAGTACATCTCGTTTCACTTTTATACTGACATTCATATAATTAACAATAAGTATAGTTTATAATAGATACAAAAA
AAAAGTCTTGTATTACCATGTTAATAGTTACATTGTGTATAACTCTCGTAGTTTTCAAAAGTCAAGTCTTGTTGA
GTAAAAAAGTTTATTTTACCCTTGGGTTTCGACTCAAAATATTGGATTACGATCTAAGCAAGGTGTTGCAAGTAC
AAGCTAAAC

nucleotides homologous to the flanking region of *SNQ2* gene

hphMX6 cassette (hygromycin B resistance)

a primer binding site

Repair DNA - *gal1*Δ::EGFP-SAVLQ-Mpro

ACGAATCAAATTAACAACCATAGGATGATAATGCGATTAGTTTTTTAGCCTTATTTCTGGGGTAATTAATCAGCG
AAGCGATGATTTTTGATCTATTAACAGATATATAAATGGAAAAGCTGCATAACCACTTTAACTAATACTTTCAAC
ATTTTCAGTTTGTATTACTTCTTATTCAAATGTCATAAAAGTATCAACAAAAAATTGTTAATATACCTCTATACT
TTAACGTCAAGGAGAAAAAACTATAATGTCTAAAGGTGAAGAATTATTCACTGGTGTTGTCCCAATTTTGGTTGA
ATTAGATGGTGATGTTAATGGTCACAAATTTTCTGTCTCCGGTGAAGGTGAAGGTGATGCTACTTACGGTAAATT
GACCTTAAAATTTATTTGTACTACTGGTAAATTGCCAGTTCCATGGCCAACCTTAGTCACTACTTTAACTTATGG
TGTTCAATGTTTTTCTAGATACCCAGATCATATGAAACAACATGACTTTTTCAAGTCTGCCATGCCAGAAGGTTA
TGTTCAAGAAAGAACTATTTTTTTCAAAGATGACGGTAACTACAAGACCAGAGCTGAAGTCAAGTTTGAAGGTGA
TACCTTAGTTAATAGAATCGAATTAAAAGGTATTGATTTTAAAGAAGATGGTAACATTTTAGGTCACAAATTGGA
ATACAACTATAACTCTCACAATGTTTACATCATGGCTGACAAACAAAAGAATGGTATCAAAGTTAACTTCAAAAT
TAGACACAACATTGAAGATGGTTCTGTTCAATTAGCTGACCATTATCAACAAAATACTCCAATTGGTGATGGTCC
AGTCTTGTTACCAGACAACCATTACTTATCCACTCAATCTGCCTTATCCAAAGATCCAAACGAAAAGAGAGACCA
CATGGTCTTGTTAGAATTTGTTACTGCTGCTGGTATTACCCATGGTATGGATGAATTGTACAAATCTGCTGTTTT
GCAATCTGGTTTCAGAAAGATGGCTTTCCCATCTGGTAAGGTTGAAGGTTGTATGGTTCAAGTTACTTGTGGTAC
TACTACTTTGAACGGTTTGTGGTTGGACGACGTTGTTTACTGTCCAAGACACGTTATCTGTACTTCTGAAGACAT
GTTGAACCCAAACTACGAAGACTTGTTGATCAGAAAGTCTAACCACAACTTCTTGGTTCAAGCTGGTAACGTTCA
ATTGAGAGTTATCGGTCACTCTATGCAAAACTGTGTTTTGAAGTTGAAGGTTGACACTGCTAACCCAAAGACTCC
AAAGTACAAGTTCGTTAGAATCCAACCAGGTCAAACTTTCTCTGTTTTGGCTTGTTACAACGGTTCTCCATCTGG
TGTTTACCAATGTGCTATGAGACCAAACTTCACTATCAAGGGTTCTTTCTTGAACGGTTCTTGTGGTTCTGTTGG
TTTCAACATCGACTACGACTGTGTTTCTTTCTGTTACATGCACCACATGGAATTGCCAACTGGTGTTCACGCTGG
TACTGACTTGGAAGGTAACTTCTACGGTCCATTCGTTGACAGACAAACTGCTCAAGCTGCTGGTACTGACACTAC
TATCACTGTTAACGTTTTGGCTTGGTTGTACGCTGCTGTTATCAACGGTGACAGATGGTTCTTGAACAGATTCAC
TACTACTTTGAACGACTTCAACTTGGTTGCTATGAAGTACAACTACGAACCATTGACTCAAGACCACGTTGACAT
CTTGGGTCCATTGTCTGCTCAAACTGGTATCGCTGTTTTGGACATGTGTGCTTCTTTGAAGGAATTGTTGCAAAA
CGGTATGAACGGTAGAACTATCTTGGGTTCTGCTTTGTTGGAAGACGAATTCACTCCATTCGACGTTGTTAGACA
ATGTTCTGGTGTTACTTTCCAATAAGTATACTTCTTTTTTTTACTTTGTTCAGAACAACTTCTCATTTTTTTCTA
CTCATAACTTTAGCATCACAAAATACGCAATAATAACGAGTAGTAACACTTTTATAGTTCATACATGCTTCAACT
ACTTAATAAATGATTGTATGATAATGTTTTCAATGTAAGAGATTTCGATTATCCACAAACTTTAAAACACAGGGA
CAAAATTCTTGATATGCTTTCAACCGCTGCGTTTTGGATACCTATTCTTGACAT

nucleotides homologous to the flanking region of *GAL1* gene

EGFP codon optimized coding sequences

$M^{pro}$ codon optimized coding sequences

The sequence encoding the linking region (SAVLQ) recognized and cleaved by $M^{pro}$

a primer binding site

Repair DNA - *gal1*Δ::EGFP-$D_4$K-Mpro

ACGAATCAAATTAACAACCATAGGATGATAATGCGATTAGTTTTTTAGCCTTATTTCTGGGGTAATTAATCAGCG
AAGCGATGATTTTTGATCTATTAACAGATATATAAATGGAAAAGCTGCATAACCACTTTAACTAATACTTTCAAC
ATTTTCAGTTTGTATTACTTCTTATTCAAATGTCATAAAAGTATCAACAAAAAATTGTTAATATACCTCTATACT
TTAACGTCAAGGAGAAAAAACTATAATGTCTAAAGGTGAAGAATTATTCACTGGTGTTGTCCCAATTTTGGTTGA
ATTAGATGGTGATGTTAATGGTCACAAATTTTCTGTCTCCGGTGAAGGTGAAGGTGATGCTACTTACGGTAAATT
GACCTTAAAATTTATTTGTACTACTGGTAAATTGCCAGTTCCATGGCCAACCTTAGTCACTACTTTAACTTATGG
TGTTCAATGTTTTTCTAGATACCCAGATCATATGAAACAACATGACTTTTTCAAGTCTGCCATGCCAGAAGGTTA
TGTTCAAGAAAGAACTATTTTTTTCAAAGATGACGGTAACTACAAGACCAGAGCTGAAGTCAAGTTTGAAGGTGA
TACCTTAGTTAATAGAATCGAATTAAAAGGTATTGATTTTAAAGAAGATGGTAACATTTTAGGTCACAAATTGGA
ATACAACTATAACTCTCACAATGTTTACATCATGGCTGACAAACAAAAGAATGGTATCAAAGTTAACTTCAAAAT
TAGACACAACATTGAAGATGGTTCTGTTCAATTAGCTGACCATTATCAACAAAATACTCCAATTGGTGATGGTCC
AGTCTTGTTACCAGACAACCATTACTTATCCACTCAATCTGCCTTATCCAAAGATCCAAACGAAAAGAGAGACCA
CATGGTCTTGTTAGAATTTGTTACTGCTGCTGGTATTACCCATGGTATGGATGAATTGTACAAAGACGACGACGA
CAAGTCTGGTTTCAGAAAGATGGCTTTCCCATCTGGTAAGGTTGAAGGTTGTATGGTTCAAGTTACTTGTGGTAC
TACTACTTTGAACGGTTTGTGGTTGGACGACGTTGTTTACTGTCCAAGACACGTTATCTGTACTTCTGAAGACAT
GTTGAACCCAAACTACGAAGACTTGTTGATCAGAAAGTCTAACCACAACTTCTTGGTTCAAGCTGGTAACGTTCA
ATTGAGAGTTATCGGTCACTCTATGCAAAACTGTGTTTTGAAGTTGAAGGTTGACACTGCTAACCCAAAGACTCC
AAAGTACAAGTTCGTTAGAATCCAACCAGGTCAAACTTTCTCTGTTTTGGCTTGTTACAACGGTTCTCCATCTGG
TGTTTACCAATGTGCTATGAGACCAAACTTCACTATCAAGGGTTCTTTCTTGAACGGTTCTTGTGGTTCTGTTGG
TTTCAACATCGACTACGACTGTGTTTCTTTCTGTTACATGCACCACATGGAATTGCCAACTGGTGTTCACGCTGG
TACTGACTTGGAAGGTAACTTCTACGGTCCATTCGTTGACAGACAAACTGCTCAAGCTGCTGGTACTGACACTAC
TATCACTGTTAACGTTTTGGCTTGGTTGTACGCTGCTGTTATCAACGGTGACAGATGGTTCTTGAACAGATTCAC
TACTACTTTGAACGACTTCAACTTGGTTGCTATGAAGTACAACTACGAACCATTGACTCAAGACCACGTTGACAT
CTTGGGTCCATTGTCTGCTCAAACTGGTATCGCTGTTTTGGACATGTGTGCTTCTTTGAAGGAATTGTTGCAAAA
CGGTATGAACGGTAGAACTATCTTGGGTTCTGCTTTGTTGGAAGACGAATTCACTCCATTCGACGTTGTTAGACA
ATGTTCTGGTGTTACTTTCCAATAAGTATACTTCTTTTTTTTACTTTGTTCAGAACAACTTCTCATTTTTTTCTA
CTCATAACTTTAGCATCACAAAATACGCAATAATAACGAGTAGTAACACTTTTATAGTTCATACATGCTTCAACT
ACTTAATAAATGATTGTATGATAATGTTTTCAATGTAAGAGATTTCGATTATCCACAAACTTTAAAACACAGGGA
CAAAATTCTTGATATGCTTTCAACCGCTGCGTTTTGGATACCTATTCTTGACAT

nucleotides homologous to the flanking region of *GAL1* gene

EGFP codon optimized coding sequences

$M^{pro}$ codon optimized coding sequences

The sequence encoding the linking region (DDDDK) recognized and cleaved by enterokinase

a primer binding site

Repair DNA - *gal1*Δ::mCherry

ACGAATCAAATTAACAACCATAGGATGATAATGCGATTAGTTTTTTAGCCTTATTTCTGGGGTAATTAATCAGCG
AAGCGATGATTTTTGATCTATTAACAGATATATAAATGGAAAAGCTGCATAACCACTTTAACTAATACTTTCAAC
ATTTTCAGTTTGTATTACTTCTTATTCAAATGTCATAAAAGTATCAACAAAAAATTGTTAATATACCTCTATACT
TTAACGTCAAGGAGAAAAAACTATAATGGTGAGCAAGGGCGAGGAGGATAACATGGCCATCATCAAGGAGTTCAT
GCGCTTCAAGGTGCACATGGAGGGCTCCGTGGACGGCCACGAGTTCGAGATCGAGGGCGAGGGCGAGGGCCGCCC
CTACGAGGGCACCCAGACCGCCAAGCTGAAGGTGACCAAGGGTGGCCCCCTGCCCTTCGCCTGGGACATCCTGTC
CCCTCAGTTCATGTACGGCTCCAAGGCCTACGTGAAGCACCCCGCCGACATCCCCGACTACTTGAAGCTGTCCTT
CCCCGAGGGCTTCAAGTGGGAGCGCGTGATGAACTTCGAGGACGGCGGCGTGGTGACCGTGACCCAGGACTCCTC
CCTGCAGGACGGCGAGTTCATCTACAAGGTGAAGCTGCGCGGCACCAACTTCCCCTCCGACGGCCCCGTAATGCA
GAAGAAGACTATGGGCTGGGAGGCCTCCTCCGAGCGGATGTACCCCGAGGACGGCGCCCTGAAGGGCGAGATCAA
GCAGAGGCTGAAGCTGAAGGACGGCGGCCACTACGACGCTGAGGTCAAGACCACCTACAAGGCCAAGAAGCCCGT
GCAGCTGCCCGGCGCCTACAACGTCAACATCAAGTTGGACATCACCTCCCACAACGAGGACTACACCATCGTGGA
ACAGTACGAACGCGCCGAGGGCCGCCACTCCACCGGCGGCATGGACGAGCTGTACAAGTAAGTATACTTCTTTTT
TTTACTTTGTTCAGAACAACTTCTCATTTTTTTCTACTCATAACTTTAGCATCACAAAATACGCAATAATAACGA
GTAGTAACACTTTTATAGTTCATACATGCTTCAACTACTTAATAAATGATTGTATGATAATGTTTTCAATGTAAG
AGATTTCGATTATCCACAAACTTTAAAACACAGGGACAAAATTCTTGATATGCTTTCAACCGCTGCGTTTTGGAT
ACCTATTCTTGACAT

nucleotides homologous to the flanking region of *GAL1* gene

mCherry coding sequences

a primer binding site
