## Supplementary File 2 for "Meisoindigo: An Effective Inhibitor of SARS-CoV-2 Main Protease Revealed by Yeast System"

**Title**: Meisoindigo: An Effective Inhibitor of SARS-CoV-2 Main Protease Revealed by Yeast System

**Authors**: Wojciech Grabiński, Anna Kicińska, Ewa Kosicka, Martyna Baranek-Grabińska, Ewelina Hejenkowska, Joanna Budzik, Paulina Śliska, Weronika Śliwińska, Andonis Karachitos*

**Description**: The file contains images of 31 scanned yeast culture plates after screening with a library of drugs. Each drug on a single filter has an assigned number on each plate. Each plate was scanned after 24 and 48 hours of culture from the start of the screening. A table listing the drugs by number is shown below each plate image. Drugs marked in red in the table were selected for subsequent stages of the study.

PLATE 1

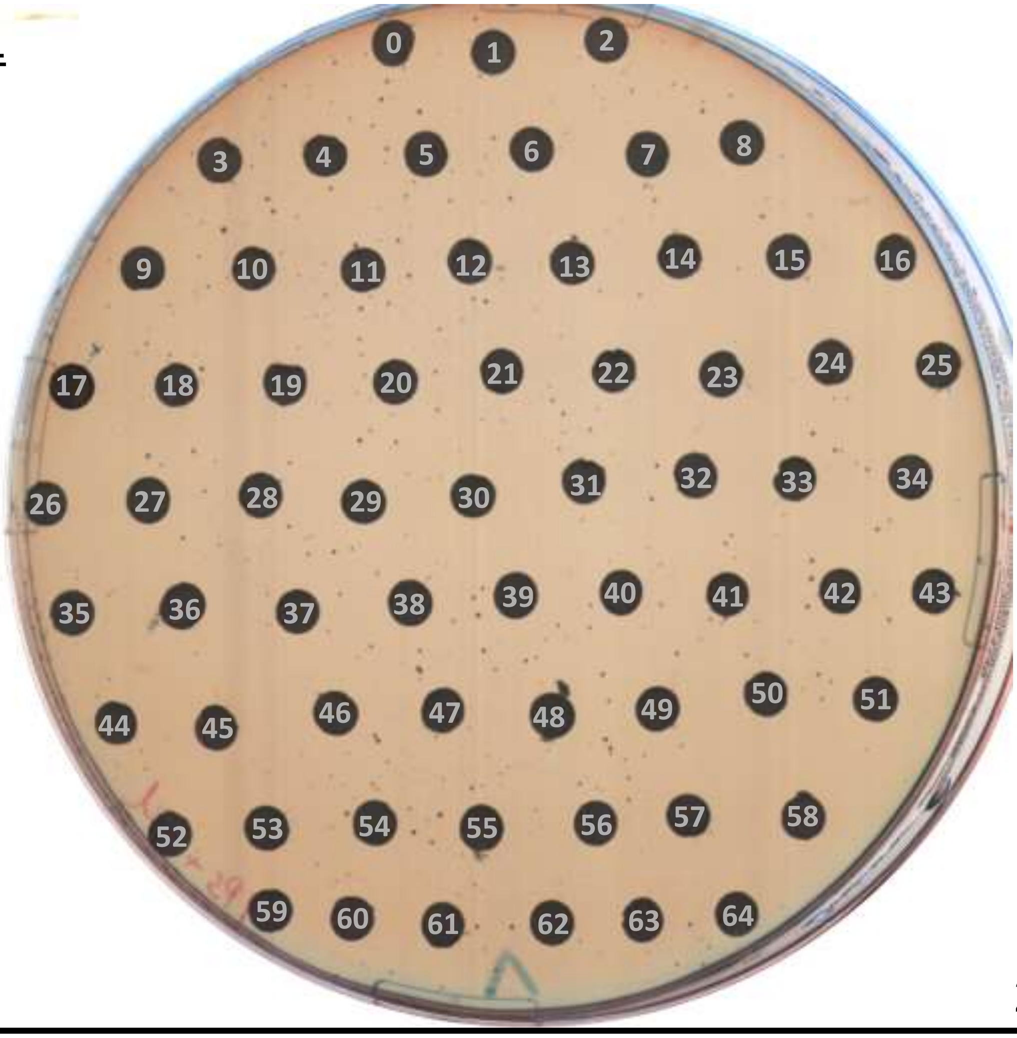

24 h

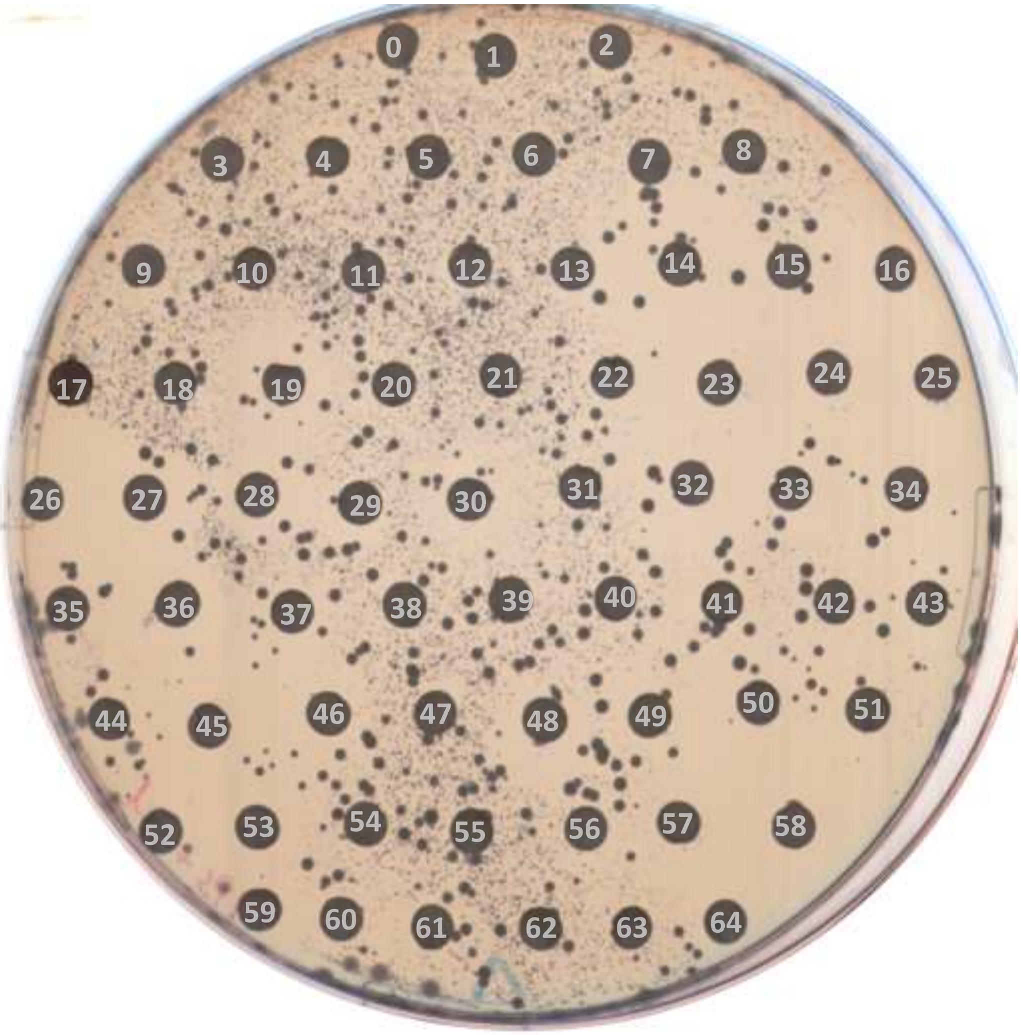

48 h

### PLATE 1

| no. | compound | no. | compound | no. | compound |
|---|---|---|---|---|---|
| 0 | DMSO | 22 | Faropenem sodium | 44 | Levetiracetam |
| 1 | Hydrocortisone buteprate | 23 | Nefazodone (hydrochloride) | 45 | Ornidazole (Levo-) |
| 2 | Fusidic acid (sodium salt) | 24 | Puromycin (dihydrochloride) | 46 | Artemether |
| 3 | Dithranol | 25 | Prasugrel (hydrochloride) | 47 | Troxipide |
| 4 | Oxiconazole nitrate | 26 | Tinoridine hydrochloride | 48 | Lesinurad (sodium) |
| 5 | Resveratrol | 27 | Tolazamide | 49 | Fexofenadine (hydrochloride) |
| 6 | Ethoxzolamide | 28 | Morinidazole (R enantiomer) | 50 | Treprostinil (sodium) |
| 7 | Ilaprazole | 29 | Rolapitant | 51 | Nadifloxacin |
| 8 | Morinidazole | 30 | Revefenacin | 52 | Guacetisal |
| 9 | Oseltamivir acid | 31 | Benzamil (hydrochloride) | 53 | Benactyzine hydrochloride |
| 10 | Mozavaptan | 32 | 2-Ethoxybenzamide | 54 | Nitroxoline |
| 11 | Bromisoval | 33 | Ivermectin | 55 | Tripelennamine (hydrochloride) |
| 12 | Tetrahydrobiopterin | 34 | Cyclobenzaprine (hydrochloride) | 56 | Benzyl benzoate |
| 13 | Amfenac (Sodium Hydrate) | 35 | Emamectin (Benzoate) | 57 | Levodropropizine |
| 14 | Hexaminolevulinate (hydrochloride) | 36 | Ipriflavone | 58 | Piperacillin (sodium) |
| 15 | Cefozopran (hydrochloride) | 37 | Terfenadine | 59 | Mafenide (Acetate) |
| 16 | Miglustat (hydrochloride) | 38 | Iopromide | 60 | Methylbenactyzium Bromide |
| 17 | Buspirone (hydrochloride) | 39 | Yohimbine (Hydrochloride) | 61 | Medroxyprogesterone acetate |
| 18 | Bleomycin (sulfate) | 40 | Capsaicin | 62 | Sorafenib |
| 19 | Terpin (hydrate) | 41 | Raltitrexed | 63 | Triflupromazine (hydrochloride) |
| 20 | Apronal | 42 | Dihydroergotamine (mesylate) | 64 | Cefamandole (sodium) |
| 21 | Alarelin (Acetate) | 43 | Vitamin K1 | - | - |

PLATE 2

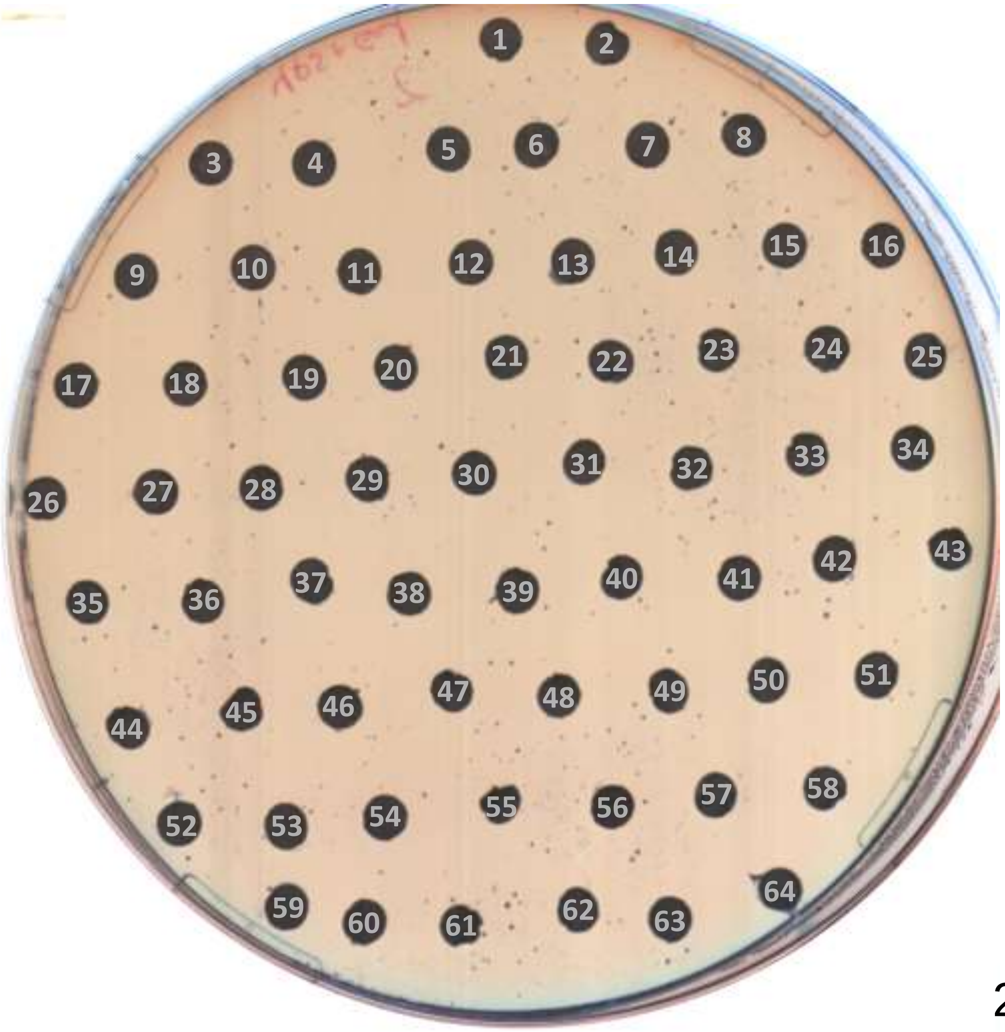

24 h

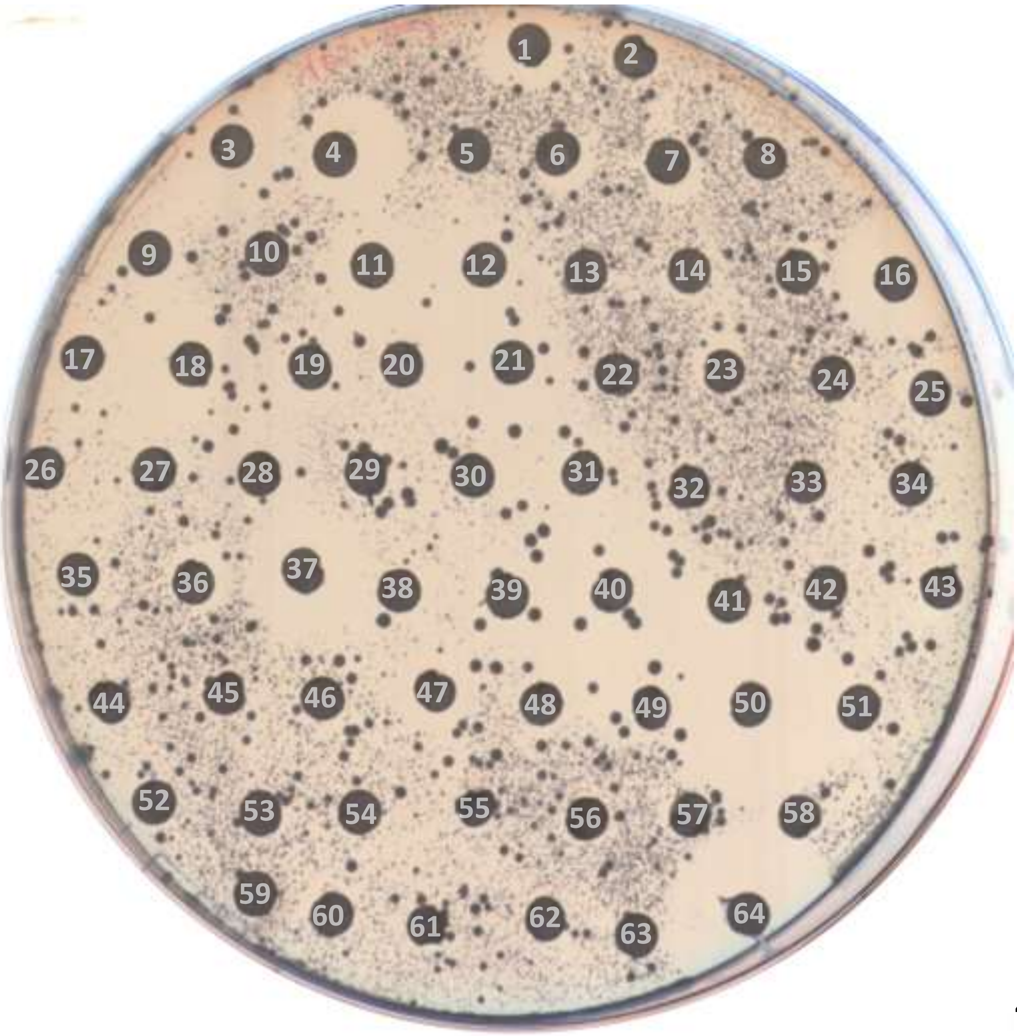

48 h

### PLATE 2

| no. | compound | no. | compound | no. | compound |
|---|---|---|---|---|---|
| - | - | 22 | Sildenafil (citrate) | 44 | Naphazoline (hydrochloride) |
| 1 | Riboflavin Tetrabutyrate | 23 | Crizotinib (hydrochloride) | 45 | Avanafil |
| 2 | Ibuprofen | 24 | Orotic acid | 46 | Nifuratel |
| 3 | Dinoprost (tromethamine salt) | 25 | Pravastatin (sodium) | 47 | Ebastine |
| 4 | Ketoprofen | 26 | Ethionamide | 48 | Fluticasone (propionate) |
| 5 | 6-Mercaptopurine | 27 | Grazoprevir potassium salt | 49 | Mosapride (citrate) |
| 6 | Trimethobenzamide hydrochloride | 28 | Chloroxine | 50 | Xylitol |
| 7 | Sodium gualenate | 29 | Parecoxib | 51 | Betaxolol (hydrochloride) |
| 8 | Dimenhydrinate | 30 | Didanosine | 52 | Nateglinide |
| 9 | Felbamate | 31 | Paroxetine (hydrochloride) | 53 | Edoxaban (tosylate monohydrate) |
| 10 | Benzbromarone | 32 | Nitisinone | 54 | Fenoldopam (mesylate) |
| 11 | 5-Azacytidine | 33 | Aspirin | 55 | Octocrylene |
| 12 | Etripamil | 34 | Allopurinol | 56 | L-Thyroxine |
| 13 | Esmolol (hydrochloride) | 35 | Irsogladine | 57 | Chlorpheniramine (maleate) |
| 14 | Brivudine | 36 | Palbociclib (isethionate) | 58 | Vildagliptin |
| 15 | Quinidine | 37 | Fluoxetine (hydrochloride) | 59 | Dapagliflozin |
| 16 | Anidulafungin | 38 | Diphylline | 60 | Alibendol |
| 17 | Atropine (sulfate monohydrate) | 39 | Pitavastatin (Calcium) | 61 | Daptomycin |
| 18 | Danazol | 40 | Ingenol Mebutate | 62 | Emtricitabine |
| 19 | Triclabendazole | 41 | Dibucaine (hydrochloride) | 63 | Sorafenib (Tosylate) |
| 20 | Mexiletine (hydrochloride) | 42 | Lansoprazole | 64 | Sertraline (hydrochloride) |
| 21 | Nicardipine (hydrochloride) | 43 | Dienogest | - | - |

PLATE 3

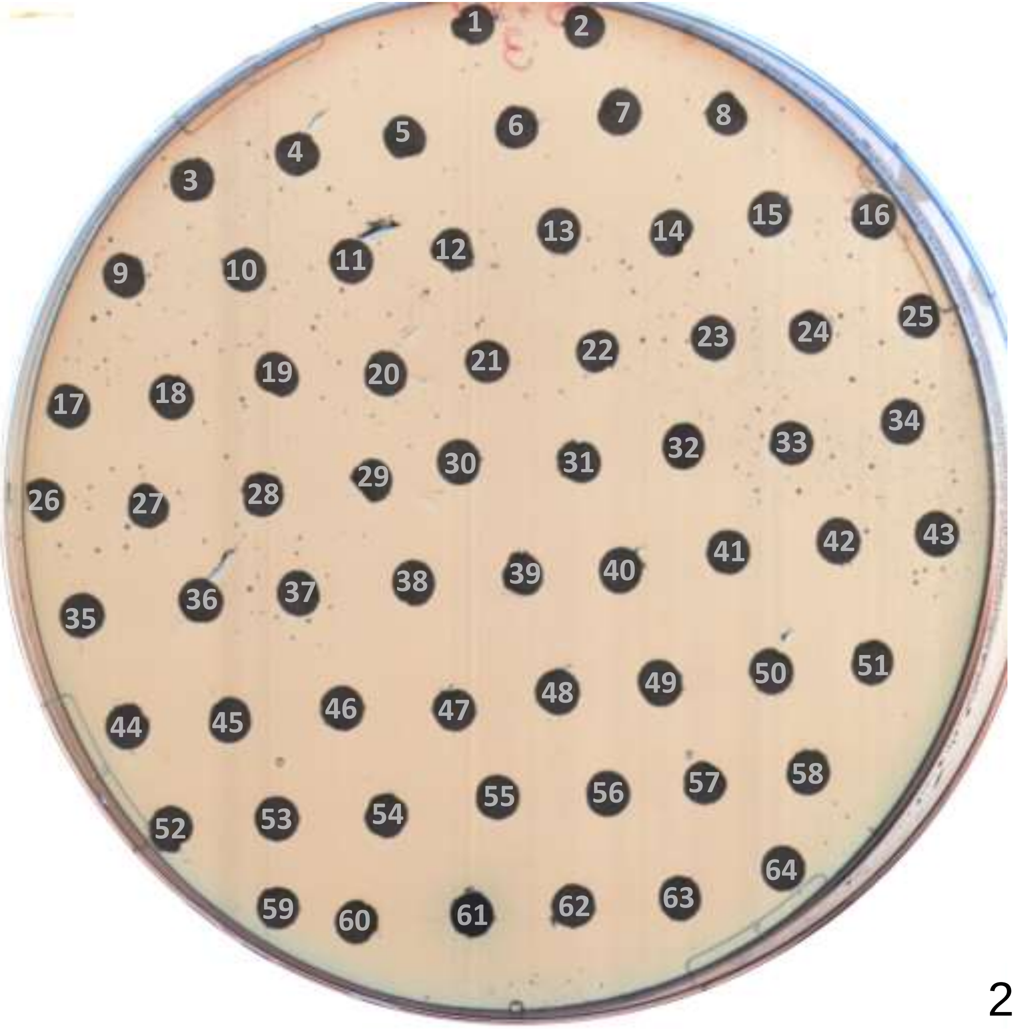

24 h

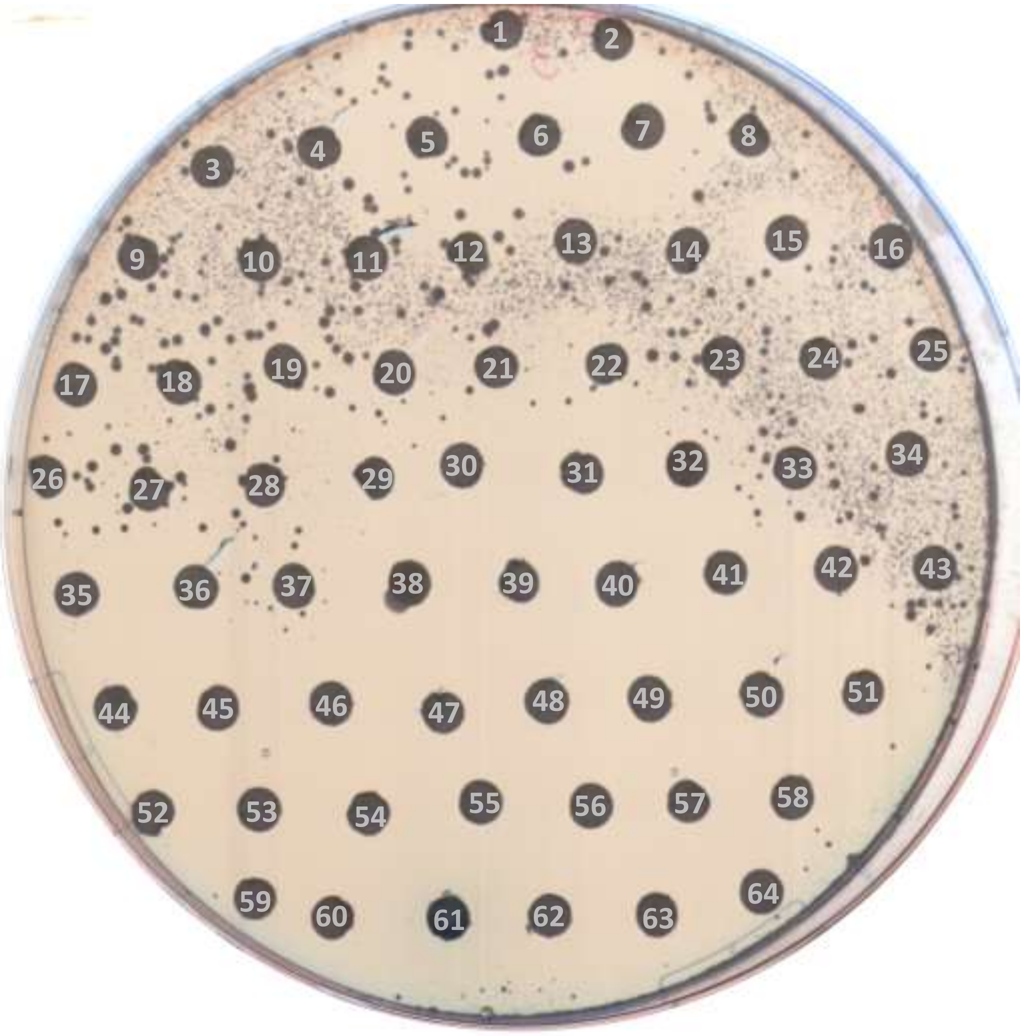

48 h

### PLATE 3

| no. | compound | no. | compound | no. | compound |
|---|---|---|---|---|---|
| - | - | 22 | Sodium nitroprusside | 44 | Tioconazole |
| 1 | Telaprevir | 23 | Isosorbide mononitrate | 45 | Carbamazepine |
| 2 | Nepafenac | 24 | Ponatinib | 46 | Clofibrate |
| 3 | Cidofovir | 25 | Phenformin (hydrochloride) | 47 | Vilanterol (trifenatate) |
| 4 | Alvimopan (dihydrate) | 26 | Umifenovir (hydrochloride) | 48 | Efinaconazole |
| 5 | DHEA | 27 | Pentostatin | 49 | Iloperidone |
| 6 | Mitiglinide (Calcium) | 28 | Canagliflozin (hemihydrate) | 50 | Simvastatin |
| 7 | Fenticonazole (Nitrate) | 29 | Pioglitazone (hydrochloride) | 51 | Irbesartan |
| 8 | Azathioprine | 30 | Cortisone acetate | 52 | Acitretin |
| 9 | Ramosetron (Hydrochloride) | 31 | Loperamide (hydrochloride) | 53 | Betaine (hydrochloride) |
| 10 | Zidovudine | 32 | Embelin | 54 | Migalastat (hydrochloride) |
| 11 | Etidronic acid | 33 | Febuxostat | 55 | Orlistat |
| 12 | Cefprozil (monohydrate) | 34 | Fenoprofen (Calcium hydrate) | 56 | Sodium diatrizoate |
| 13 | Cilostazol | 35 | Guanabenz (Acetate) | 57 | D-Mannitol |
| 14 | Azithromycin | 36 | Cefditoren (Pivoxil) | 58 | Hydroquinidine |
| 15 | Tamoxifen (Citrate) | 37 | Sumatriptan (succinate) | 59 | Levocarnitine propionate |
| 16 | Rimantadine (hydrochloride) | 38 | Glibenclamide | 60 | Chlorquinaldol |
| 17 | Mitotane | 39 | Tropicamide | 61 | Mitoxantrone |
| 18 | Loxapine | 40 | Carbidopa | 62 | Nortriptyline (hydrochloride) |
| 19 | Carvedilol | 41 | Nevirapine | 63 | Aripiprazole |
| 20 | Dexmedetomidine (hydrochloride) | 42 | Meropenem (trihydrate) | 64 | Belotecan (hydrochloride) |
| 21 | Procainamide (hydrochloride) | 43 | Mirabegron | - | - |

PLATE 4

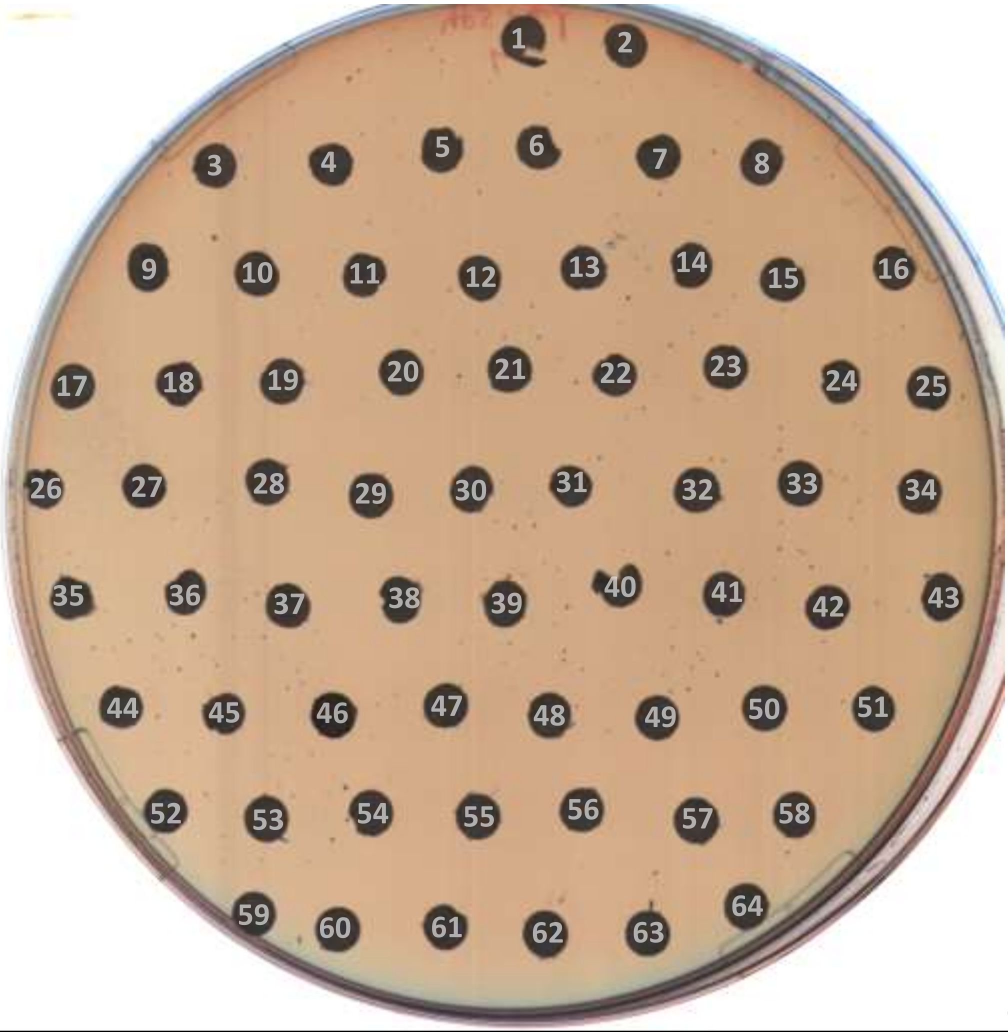

24 h

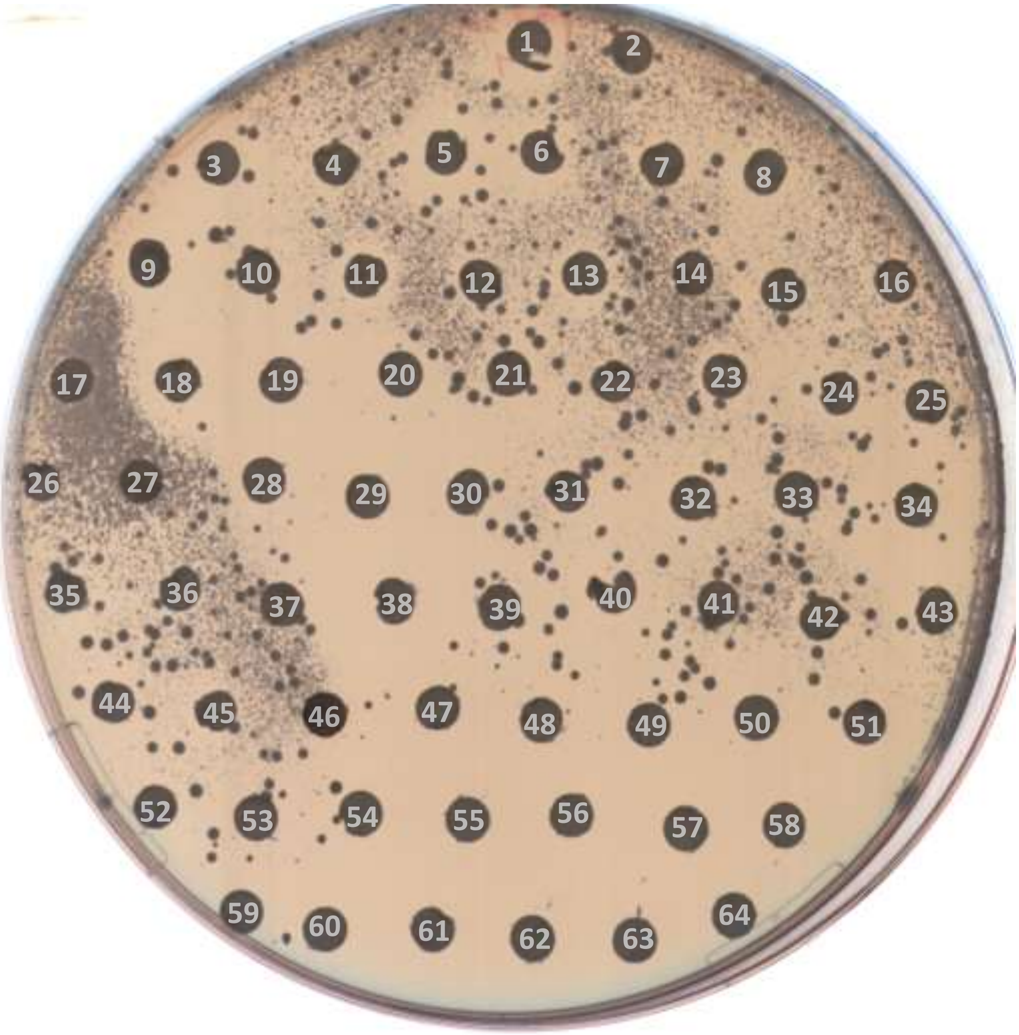

48 h

### PLATE 4

| no. | compound | no. | compound | no. | compound |
|---|---|---|---|---|---|
| - | - | 22 | Eslicarbazepine acetate | 44 | Alprenolol (hydrochloride) |
| 1 | Garenoxacin (Mesylate hydrate) | 23 | Niraparib (hydrochloride) | 45 | Estradiol |
| 2 | Sulfacarbamide | 24 | Vorapaxar | 46 | Temoporfin |
| 3 | Cinoxacin | 25 | Diflorasone | 47 | Acetazolamide |
| 4 | Glecaprevir | 26 | Sucrose | 48 | Gemifloxacin (mesylate) |
| 5 | Dydrogesterone | 27 | L-Thyroxine | 49 | Alpha-Estradiol |
| 6 | Ticarcillin (disodium) | 28 | Fludarabine | 50 | Nelarabine |
| 7 | Ferulic acid (sodium) | 29 | Auranofin | 51 | Rotigotine (Hydrochloride) |
| 8 | Alprenolol | 30 | Pexidartinib | 52 | Ropivacaine (hydrochloride) |
| 9 | Clioquinol | 31 | Ethosuximide | 53 | Ertugliflozin |
| 10 | Enasidenib | 32 | Riluzole hydrochloride | 54 | Aspartame |
| 11 | Taurodeoxycholic acid | 33 | Vorinostat | 55 | Lofexidine |
| 12 | Zanubrutinib | 34 | Benfluorex (hydrochloride) | 56 | Pralatrexate |
| 13 | Deoxycholic acid | 35 | Flucloxacillin sodium | 57 | Gadoteridol |
| 14 | Glycerol phenylbutyrate | 36 | Linezolid | 58 | Atomoxetine (hydrochloride) |
| 15 | Cortisone | 37 | Vinorelbine (ditartrate) | 59 | Lenalidomide |
| 16 | Ioversol | 38 | Cinacalcet (hydrochloride) | 60 | Irinotecan |
| 17 | 9-Aminoacridine | 39 | Piperonyl butoxide | 61 | Hydrocortisone cypionate |
| 18 | Asenapine (hydrochloride) | 40 | Eperisone (Hydrochloride) | 62 | Urapidil |
| 19 | Bepridil hydrochloride | 41 | Daphnetin | 63 | Tiotropium (Bromide) |
| 20 | Salmeterol | 42 | Estrone | 64 | L-Lactic acid |
| 21 | Mezlocillin (sodium) | 43 | Edoxaban | - | - |

PLATE 5

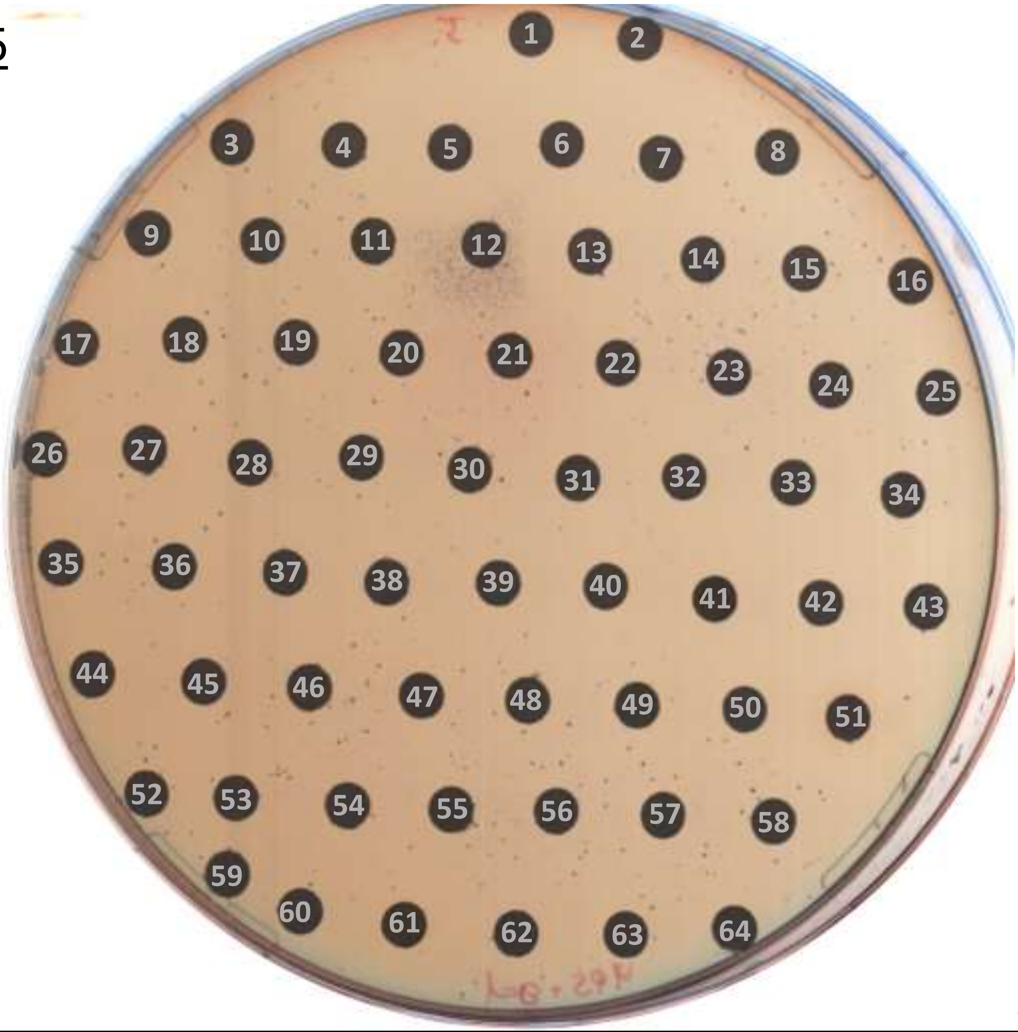

24 h

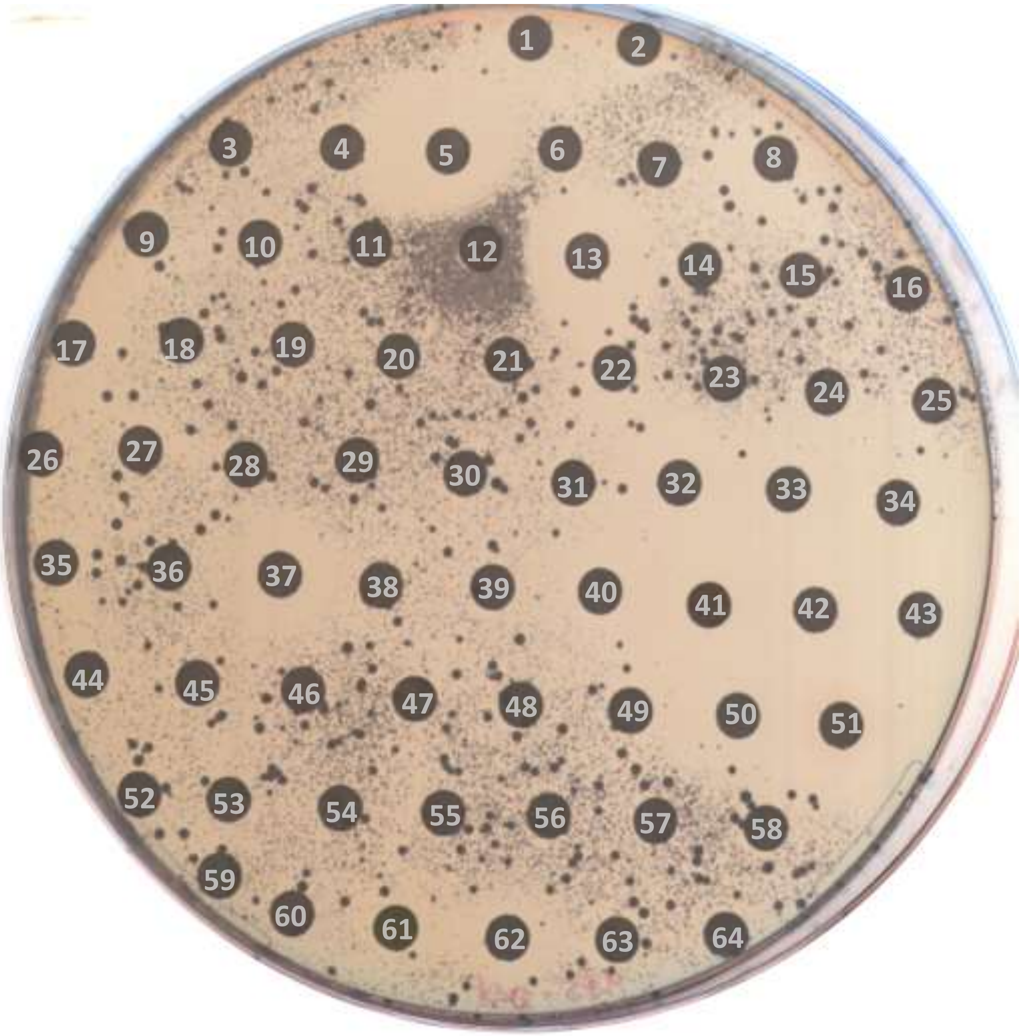

48 h

### PLATE 5

| no. | compound | no. | compound | no. | compound |
|---|---|---|---|---|---|
| - | - | 22 | Phenoxybenzamine | 44 | Binimetinib |
| 1 | 5-Fluorouracil | 23 | Lomitapide | 45 | Bicalutamide |
| 2 | Arterolane | 24 | Bimatoprost | 46 | Desonide |
| 3 | Dobutamine (hydrochloride) | 25 | Loteprednol Etabonate | 47 | Amantadine (hydrochloride) |
| 4 | Toloxatone | 26 | Bendamustine (hydrochloride) | 48 | Racecadotril |
| 5 | Pimavanserin tartrate | 27 | Epinastine | 49 | Tolvaptan |
| 6 | Hydroxyzine (dihydrochloride) | 28 | Tiopronin | 50 | Tetrahydrozoline (hydrochloride) |
| 7 | Helicid | 29 | Busulfan | 51 | Nifedipine |
| 8 | Oxybutynin | 30 | Ethambutol (dihydrochloride) | 52 | Sulfameter |
| 9 | Cevimeline (hydrochloride) | 31 | Menaquinone-4 | 53 | Deflazacort |
| 10 | Rabeprazole (sodium) | 32 | Butenafine (Hydrochloride) | 54 | Glycopyrrolate |
| 11 | Moxalactam (sodium salt) | 33 | Doxepin (Hydrochloride) | 55 | Hydroxyfasudil (hydrochloride) |
| 12 | Ixazomib | 34 | Sodium 4-phenylbutyrate | 56 | Sarpogrelate (hydrochloride) |
| 13 | Pinaverium bromide | 35 | Oxcarbazepine | 57 | Carteolol hydrochloride |
| 14 | Gallamine Triethiodide | 36 | Methylprednisolone | 58 | Vilazodone (Hydrochloride) |
| 15 | Guanidine (hydrochloride) | 37 | Clomipramine (hydrochloride) | 59 | Tranilast |
| 16 | L-(-)-α-Methyldopa (hydrate) | 38 | Mesna | 60 | Saquinavir |
| 17 | Reboxetine (mesylate) | 39 | Lacidipine | 61 | Nintedanib esylate |
| 18 | Xylometazoline (hydrochloride) | 40 | Riluzole | 62 | Amitriptyline (hydrochloride) |
| 19 | Bepotastine (Beslilate) | 41 | Oxytetracycline | 63 | Ropivacaine |
| 20 | Altretamine | 42 | Clotrimazole | 64 | Acetohexamide |
| 21 | Methylcobalamin | 43 | Edrophonium (chloride) | - | - |

PLATE 6

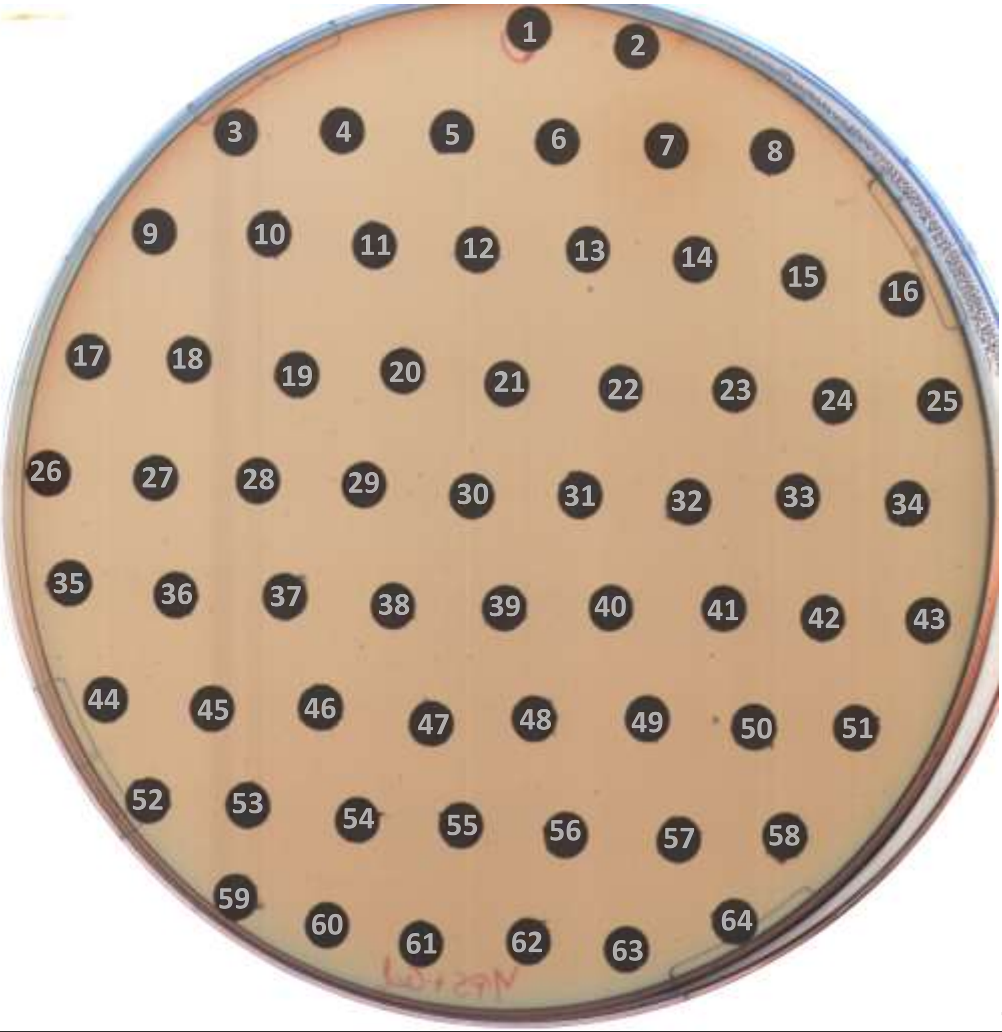

24 h

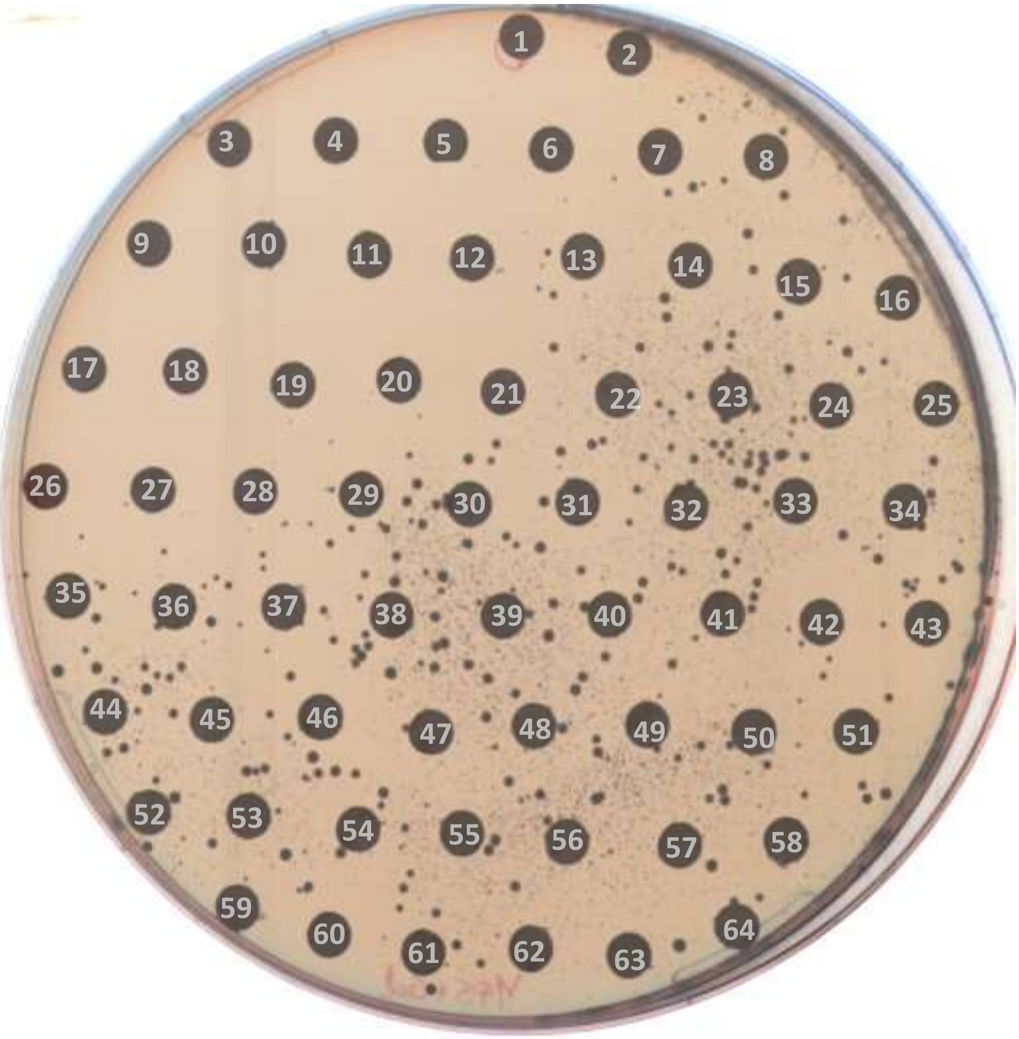

48 h

### PLATE 6

| no. | compound | no. | compound | no. | compound |
|---|---|---|---|---|---|
| - | - | 22 | Pyridostigmine (bromide) | 44 | Cisatracurium (besylate) |
| 1 | 6-Thioguanine | 23 | Lamivudine | 45 | Gabexate (mesylate) |
| 2 | Amifampridine | 24 | Aliskiren (hemifumarate) | 46 | Nilvadipine |
| 3 | Propafenone (hydrochloride) | 25 | Bambuterol hydrochloride | 47 | Duvelisib |
| 4 | Econazole (nitrate) | 26 | Clofazimine | 48 | Piracetam |
| 5 | Eflornithine | 27 | Diphenidol (hydrochloride) | 49 | Acetylcysteine |
| 6 | Elvitegravir | 28 | Clarithromycin | 50 | Prucalopride |
| 7 | Rifapentine | 29 | Ritonavir | 51 | Pramipexole (dihydrochloride) |
| 8 | Lodenafil | 30 | Cefoperazone | 52 | Clevudine |
| 9 | Luliconazole | 31 | Dabigatran etexilate | 53 | Nimorazole |
| 10 | Homatropine (methylbromide) | 32 | Tezacaftor | 54 | Agomelatine |
| 11 | Gadodiamide (hydrate) | 33 | Fludarabine (phosphate) | 55 | Salicylic acid |
| 12 | Prednisone acetate | 34 | Sonidegib | 56 | Pyridoxal phosphate |
| 13 | Fruquintinib | 35 | Silibinin | 57 | Sodium 4-aminosalicylate |
| 14 | Prednisone | 36 | Demeclocycline (hydrochloride) | 58 | Ramipril |
| 15 | Afloqualone | 37 | Oxybutynin (chloride) | 59 | Glimepiride |
| 16 | Alfuzosin | 38 | Zonisamide | 60 | Etomidate |
| 17 | Acetophenazine (dimaleate) | 39 | Olopatadine (hydrochloride) | 61 | Guaifenesin |
| 18 | Prucalopride (succinate) | 40 | Meprednisone | 62 | Salbutamol (hemisulfate) |
| 19 | (S)-Timolol (Maleate) | 41 | Brinzolamide | 63 | Tinidazole |
| 20 | Amiloride (hydrochloride) | 42 | Mupirocin | 64 | Clobetasol propionate |
| 21 | Bisacodyl | 43 | Naftopidil | - | - |

PLATE 7

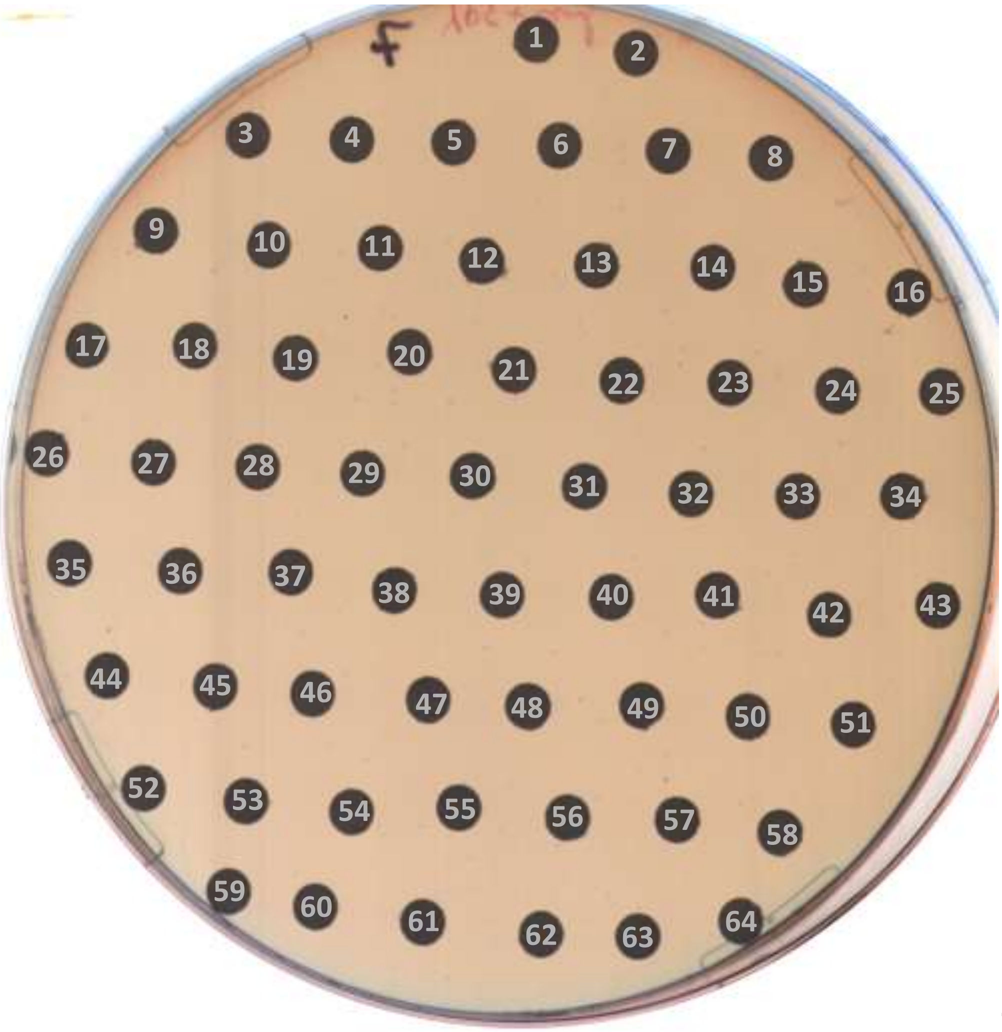

24 h

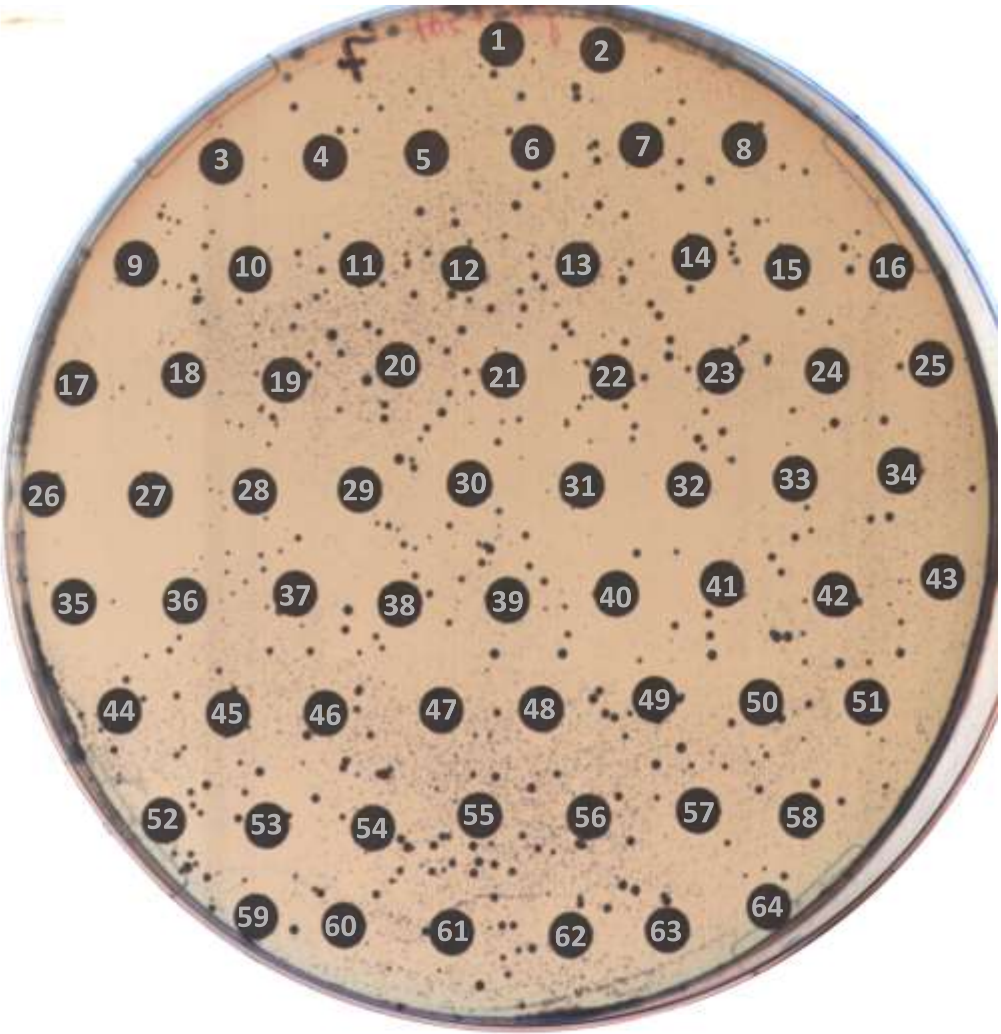

48 h

### PLATE 7

| no. | compound | no. | compound | no. | compound |
|---|---|---|---|---|---|
| - | - | 22 | Iproniazid (phosphate) | 44 | Osalmid |
| 1 | Adefovir dipivoxil | 23 | Primidone | 45 | Lodoxamide |
| 2 | Tenoxicam | 24 | Pheniramine (Maleate) | 46 | Ribociclib succinate hydrate |
| 3 | Dyclonine (hydrochloride) | 25 | Dropropizine | 47 | Goserelin (acetate) |
| 4 | Tirofiban | 26 | Nicergoline | 48 | Argatroban (monohydrate) |
| 5 | Betaxolol | 27 | Pramocaine (hydrochloride) | 49 | Dehydrocholic acid |
| 6 | Letrozole | 28 | Etofylline | 50 | Citalopram (hydrobromide) |
| 7 | Detomidine (hydrochloride) | 29 | Oxaprozin | 51 | Labetalol (hydrochloride) |
| 8 | Roxatidine | 30 | Piribedil | 52 | Tolperisone (hydrochloride) |
| 9 | Loratadine | 31 | Salicylanilide | 53 | Chlormadinone acetate |
| 10 | Adiphenine (hydrochloride) | 32 | Indinavir (sulfate) | 54 | Bemegride |
| 11 | Pemetrexed | 33 | Valbenazine | 55 | Pidotimod |
| 12 | Dipyridamole | 34 | Sulfachloropyridazine | 56 | Teniposide |
| 13 | L-5-Hydroxytryptophan | 35 | Nonivamide | 57 | Esomeprazole magnesium |
| 14 | Bendazol | 36 | Acefylline | 58 | Doravirine |
| 15 | Quinapril (hydrochloride) | 37 | Nikethamide | 59 | Flumethasone |
| 16 | Succinylsulfathiazole | 38 | Atorvastatin (hemicalcium salt) | 60 | Dirithromycin |
| 17 | Ambroxol | 39 | Dicoumarol | 61 | Acarbose |
| 18 | Sulfamethizole | 40 | Gastrodin | 62 | Varenicline |
| 19 | Secnidazole | 41 | Tenofovir alafenamide fumarate | 63 | Oxytocin (acetate) |
| 20 | Prednisolone acetate | 42 | Regorafenib | 64 | Naftidrofuryl (oxalate) |
| 21 | Ethynodiol diacetate | 43 | Pimecrolimus | - | - |

### PLATE 8

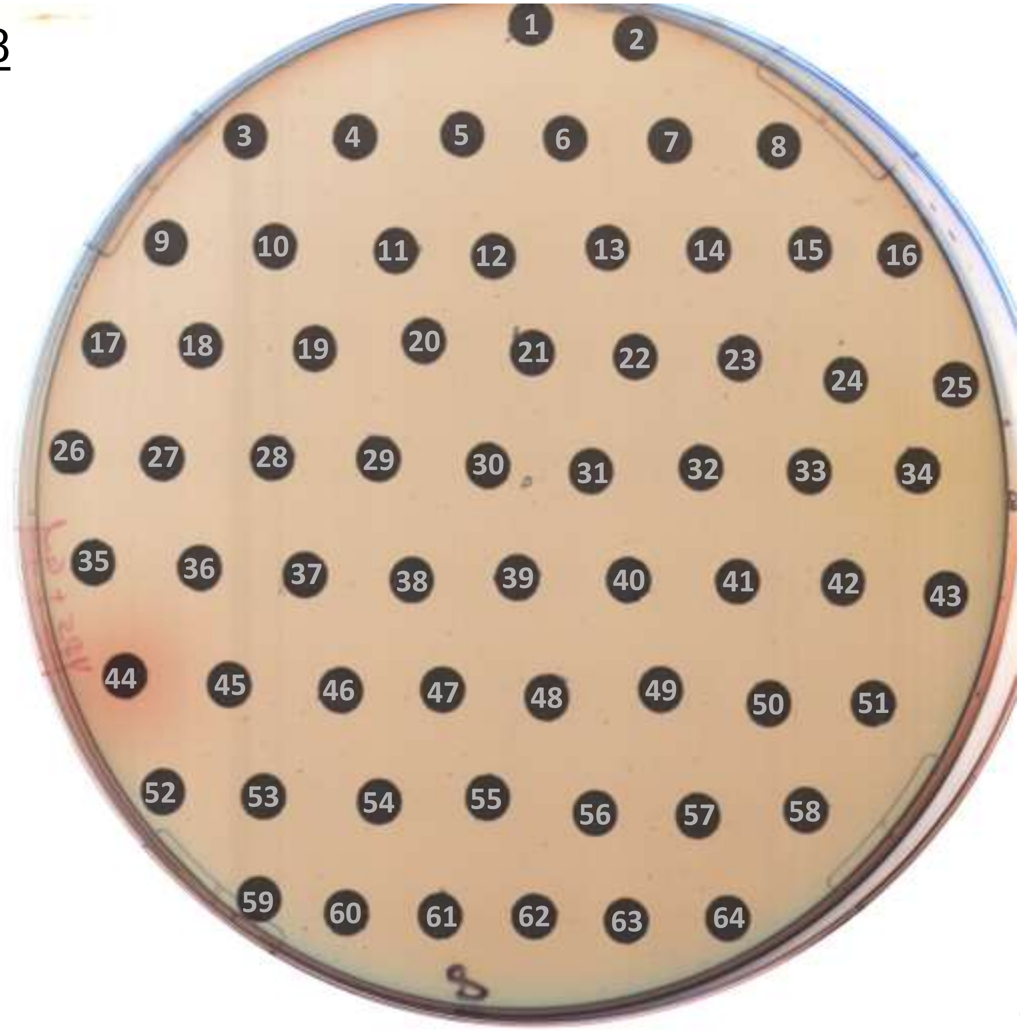

24 h

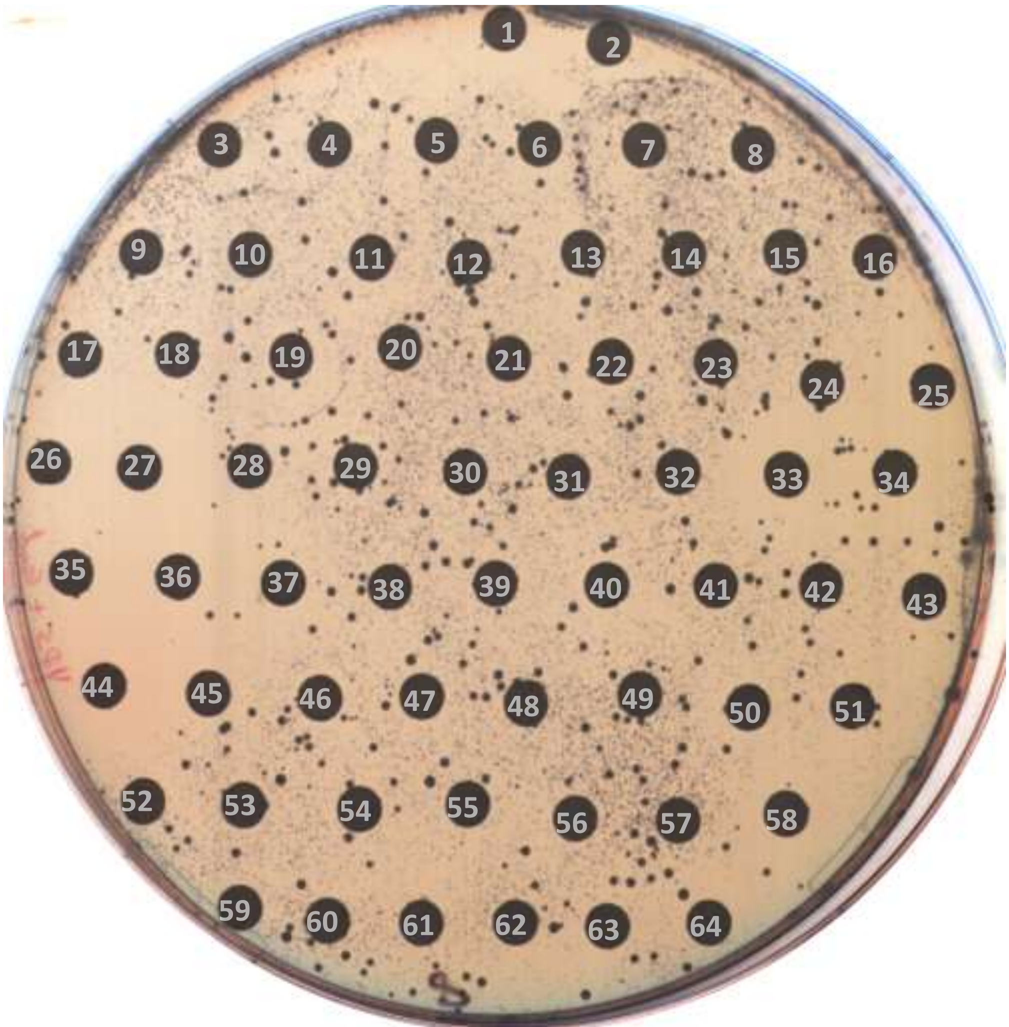

48 h

### PLATE 8

| no. | compound | no. | compound | no. | compound |
|---|---|---|---|---|---|
| - | - | 22 | 20(S)-Ginsenoside Rg3 | 44 | Daunorubicin (Hydrochloride) |
| 1 | Chlorhexidine (digluconate) | 23 | Naproxen | 45 | Diphenylpyraline (hydrochloride) |
| 2 | Midecamycin | 24 | Decloxizine (dihydrochloride) | 46 | Revaprazan (hydrochloride) |
| 3 | Aliskiren | 25 | Donepezil (Hydrochloride) | 47 | Pimozide |
| 4 | Icotinib | 26 | Metyrapone | 48 | DL-alpha-Tocopherol |
| 5 | Udenafil | 27 | Haloperidol | 49 | Ferulic acid |
| 6 | Afatinib | 28 | Bestatin | 50 | Phenytoin (sodium) |
| 7 | Cefuroxime (sodium) | 29 | Ospemifene | 51 | Lifitegrast |
| 8 | Tebipenem pivoxil | 30 | Prulifloxacin | 52 | Methylthiouracil |
| 9 | (-)-Sparteine | 31 | Itopride (hydrochloride) | 53 | Prothionamide |
| 10 | Diflunisal | 32 | Iopamidol | 54 | Cytidine |
| 11 | Varenicline (Hydrochloride) | 33 | Mefloquine (hydrochloride) | 55 | Norvancomycin (hydrochloride) |
| 12 | Amoxicillin (sodium) | 34 | Amsacrine | 56 | Acetylleucine |
| 13 | Relugolix | 35 | D-Pantothenic acid (sodium) | 57 | 10-Undecenoic acid |
| 14 | Diiodohydroxyquinoline | 36 | Niraparib (tosylate) | 58 | Siponimod |
| 15 | Carbinoxamine maleate salt | 37 | Broxyquinoline | 59 | Entrectinib |
| 16 | Oxiracetam | 38 | Abacavir (sulfate) | 60 | L-Tryptophan |
| 17 | Sulfanilamide | 39 | Pentagastrin | 61 | Eliglustat |
| 18 | Clonidine (hydrochloride) | 40 | Lactulose | 62 | Mephenesin |
| 19 | Oxantel (pamoate) | 41 | Cloperastine fendizoate | 63 | Podofilox |
| 20 | Lanatoside C | 42 | Levofloxacin (hydrate) | 64 | Betrixaban |
| 21 | Ajmaline | 43 | Methacholine (chloride) | - | - |

PLATE 9

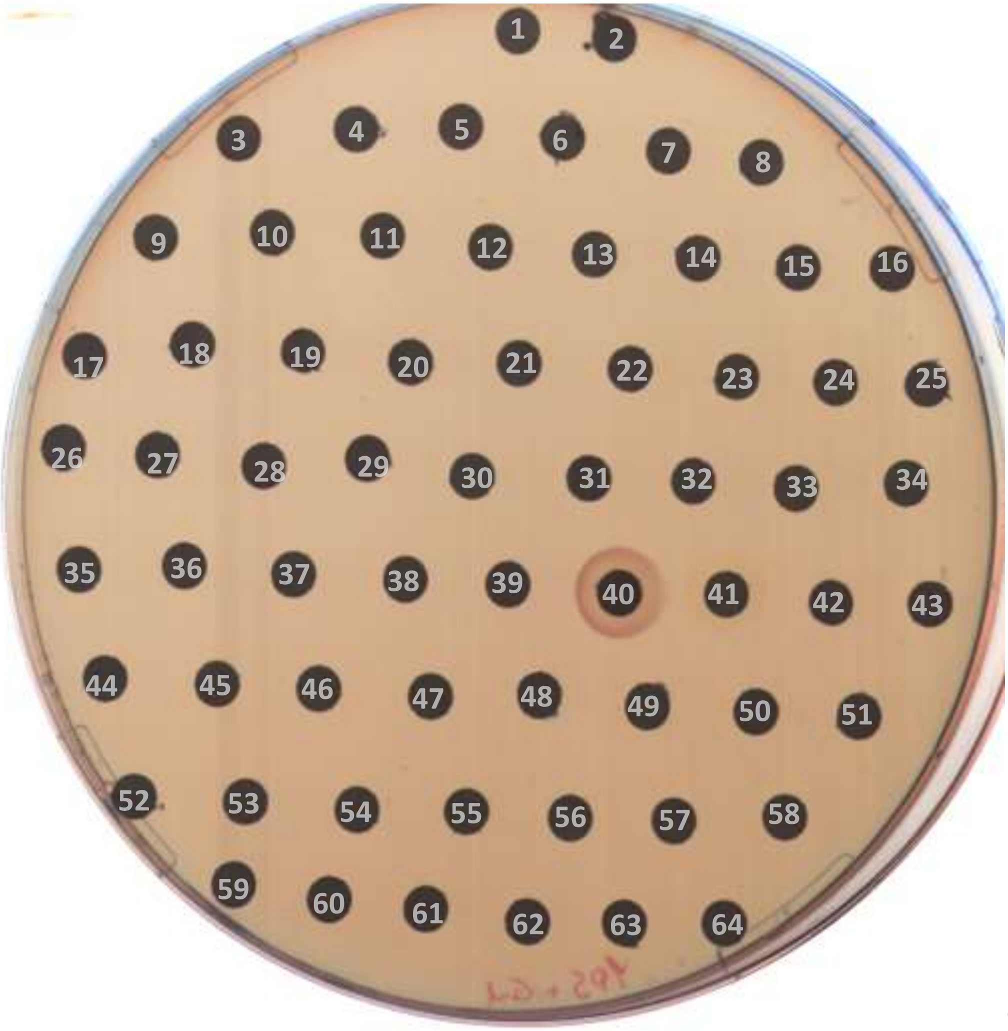

24 h

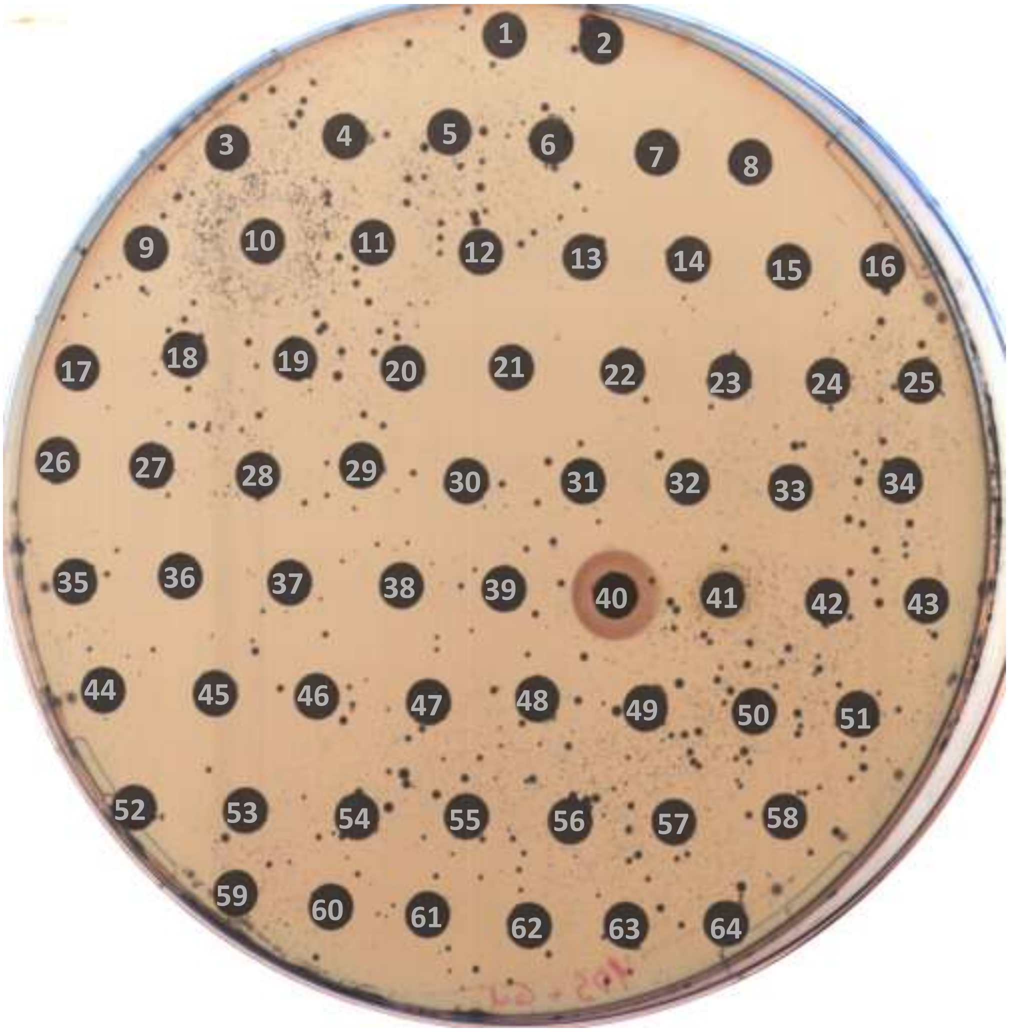

48 h

<u>PLATE 9</u>

| no. | compound | no. | compound | no. | compound |
|---|---|---|---|---|---|
| - | - | 22 | Tolmetin | 44 | Trospium (chloride) |
| 1 | Brompheniramine | 23 | Tafamidis meglumine | 45 | Alverine (citrate) |
| 2 | Isosorbide | 24 | Tricaprilin | 46 | Vinblastine (sulfate) |
| 3 | Baricitinib | 25 | Nifenazone | 47 | Nedaplatin |
| 4 | Neostigmine | 26 | Promazine | 48 | Diltiazem (hydrochloride) |
| 5 | Trifluridine/tipiracil hydrochloride | 27 | Carmofur | 49 | Nitazoxanide |
| 6 | Pamidronic acid | 28 | Carbasalate calcium | 50 | Ozagrel (sodium) |
| 7 | Minoxidil | 29 | Raltegravir (potassium salt) | 51 | Valganciclovir (hydrochloride) |
| 8 | (+)-Ketoconazole | 30 | Brimonidine (tartrate) | 52 | Felodipine |
| 9 | Cetylpyridinium | 31 | Conivaptan (hydrochloride) | 53 | Ambrisentan |
| 10 | Berberine | 32 | Fidaxomicin | 54 | Benfotiamine |
| 11 | Fadrozole | 33 | Danofloxacin (mesylate) | 55 | Ozagrel |
| 12 | Acemetacin | 34 | Pirenzepine (dihydrochloride) | 56 | Rivastigmine (tartrate) |
| 13 | Maltitol | 35 | Pyrithioxin (dihydrochloride) | 57 | Pirfenidone |
| 14 | Metformin | 36 | Trazodone (hydrochloride) | 58 | VAL-083 |
| 15 | Tetramisole | 37 | Parecoxib (Sodium) | 59 | Azelnidipine |
| 16 | Thiamine | 38 | Azaperone | 60 | Alosetron (Hydrochloride) |
| 17 | Flibanserin | 39 | D-Pantothenic acid | 61 | Pitolisant (hydrochloride) |
| 18 | Diclazuril | 40 | Tannic acid | 62 | Regorafenib (monohydrate) |
| 19 | Trimethadione | 41 | Ciclopirox | 63 | Trametinib (DMSO solvate) |
| 20 | Brimonidine | 42 | Diethylcarbamazine (citrate) | 64 | Tofacitinib |
| 21 | Cinacalcet | 43 | Darunavir (Ethanolate) | - | - |

PLATE 10

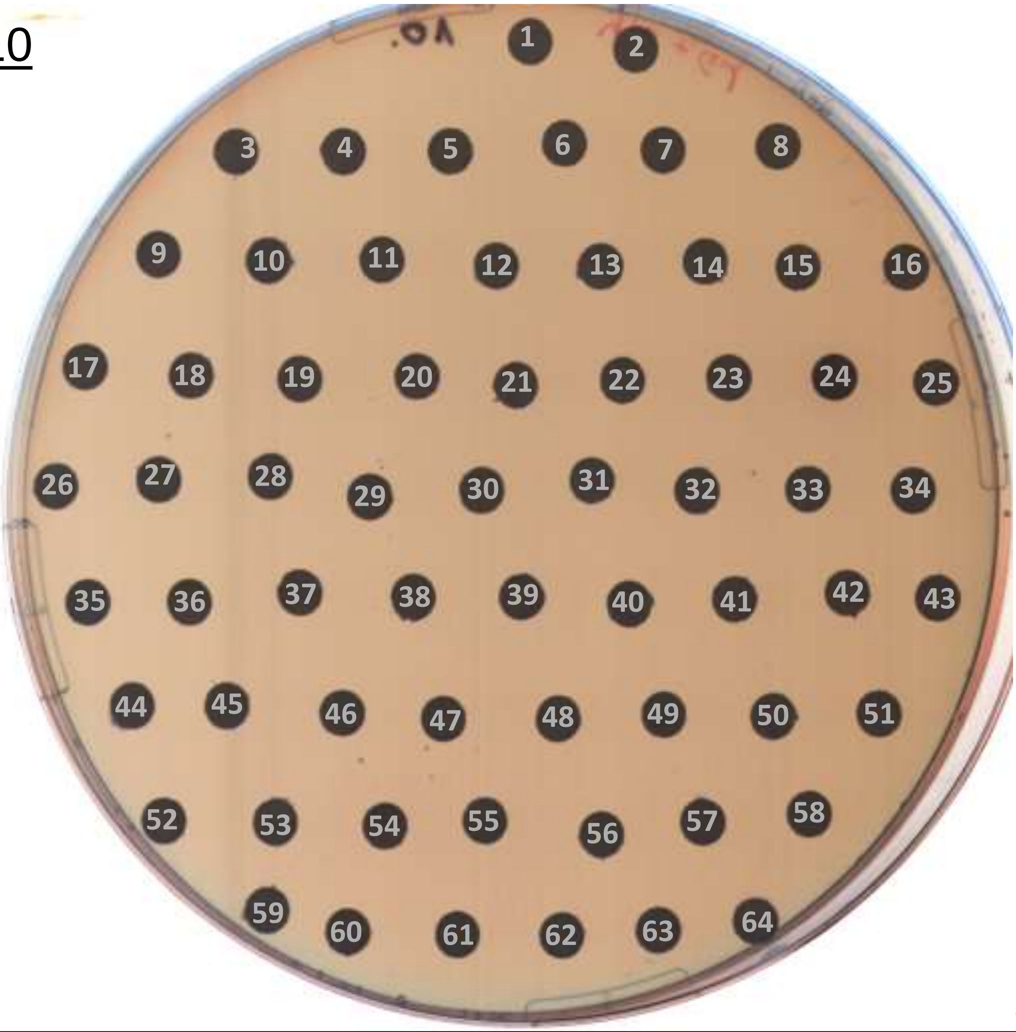

24 h

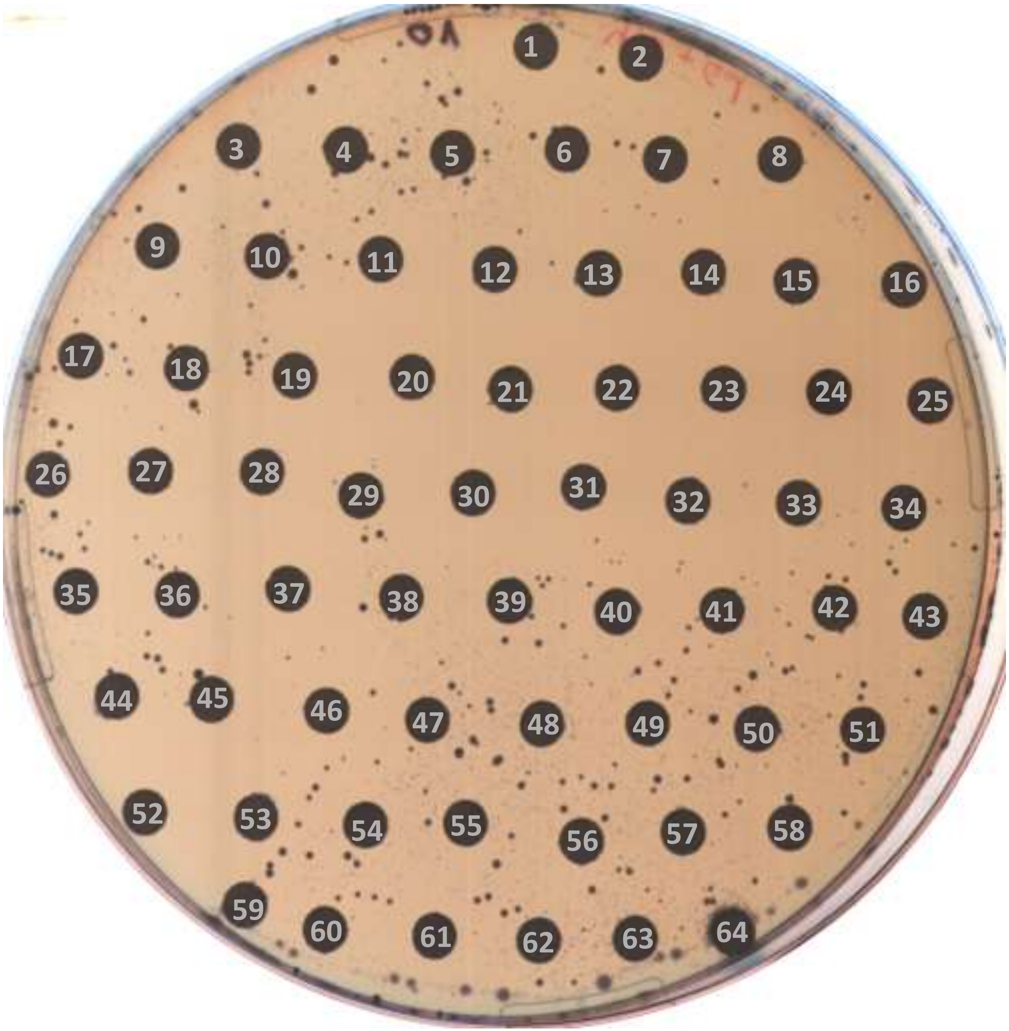

48 h

### PLATE 10

| no. | compound | no. | compound | no. | compound |
|---|---|---|---|---|---|
| - | - | 22 | Lumacaftor | 44 | Piroxicam |
| 1 | Umeclidinium (bromide) | 23 | Tavaborole | 45 | Rebamipide |
| 2 | Fenspiride (Hydrochloride) | 24 | Pixantrone (dimaleate) | 46 | Naftifine (hydrochloride) |
| 3 | Etoricoxib | 25 | Vortioxetine (hydrobromide) | 47 | Decitabine |
| 4 | Rivastigmine | 26 | Lubiprostone | 48 | Sivelestat (sodium tetrahydrate) |
| 5 | Atovaquone | 27 | Nebivolol (hydrochloride) | 49 | Aclidinium (Bromide) |
| 6 | Anastrozole | 28 | Clomiphene (citrate) | 50 | Vigabatrin |
| 7 | Encorafenib | 29 | Baricitinib (phosphate) | 51 | Losartan (potassium) |
| 8 | Nimodipine | 30 | Vortioxetine | 52 | Sertaconazole (nitrate) |
| 9 | Asenapine (maleate) | 31 | Artemisinin | 53 | Indomethacin |
| 10 | Apalutamide | 32 | Bucladesine (calcium) | 54 | Rifabutin |
| 11 | Clevidipine | 33 | Rimonabant (Hydrochloride) | 55 | Suramin (sodium salt) |
| 12 | Vonoprazan (Fumarate) | 34 | Isradipine | 56 | Ramatroban |
| 13 | Icotinib (Hydrochloride) | 35 | Isoniazid | 57 | Gadodiamide |
| 14 | Azilsartan | 36 | Benztropine (mesylate) | 58 | Cobimetinib (hemifumarate) |
| 15 | Tivozanib | 37 | Desogestrel | 59 | Saxagliptin |
| 16 | Trelagliptin (succinate) | 38 | Deracoxib | 60 | Levobetaxolol (hydrochloride) |
| 17 | Regorafenib (Hydrochloride) | 39 | Pranlukast | 61 | Quinagolide (hydrochloride) |
| 18 | Repaglinide | 40 | Doxapram (hydrochloride hydrate) | 62 | Fluocinonide |
| 19 | Maraviroc | 41 | Ibrutinib | 63 | Felbinac |
| 20 | Posaconazole | 42 | Mifepristone | 64 | Menadione |
| 21 | Pemetrexed (disodium) | 43 | Metoprolol (Succinate) | - | - |

PLATE 11

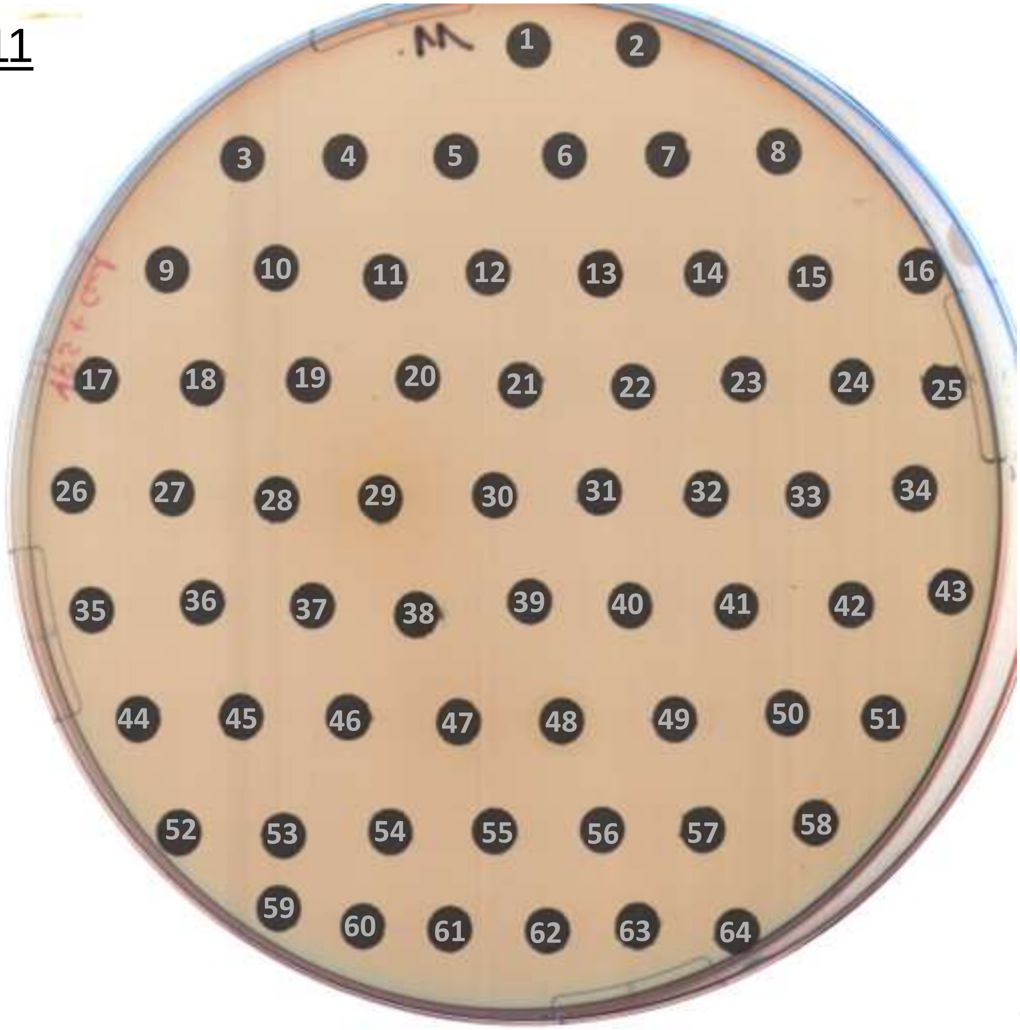

24 h

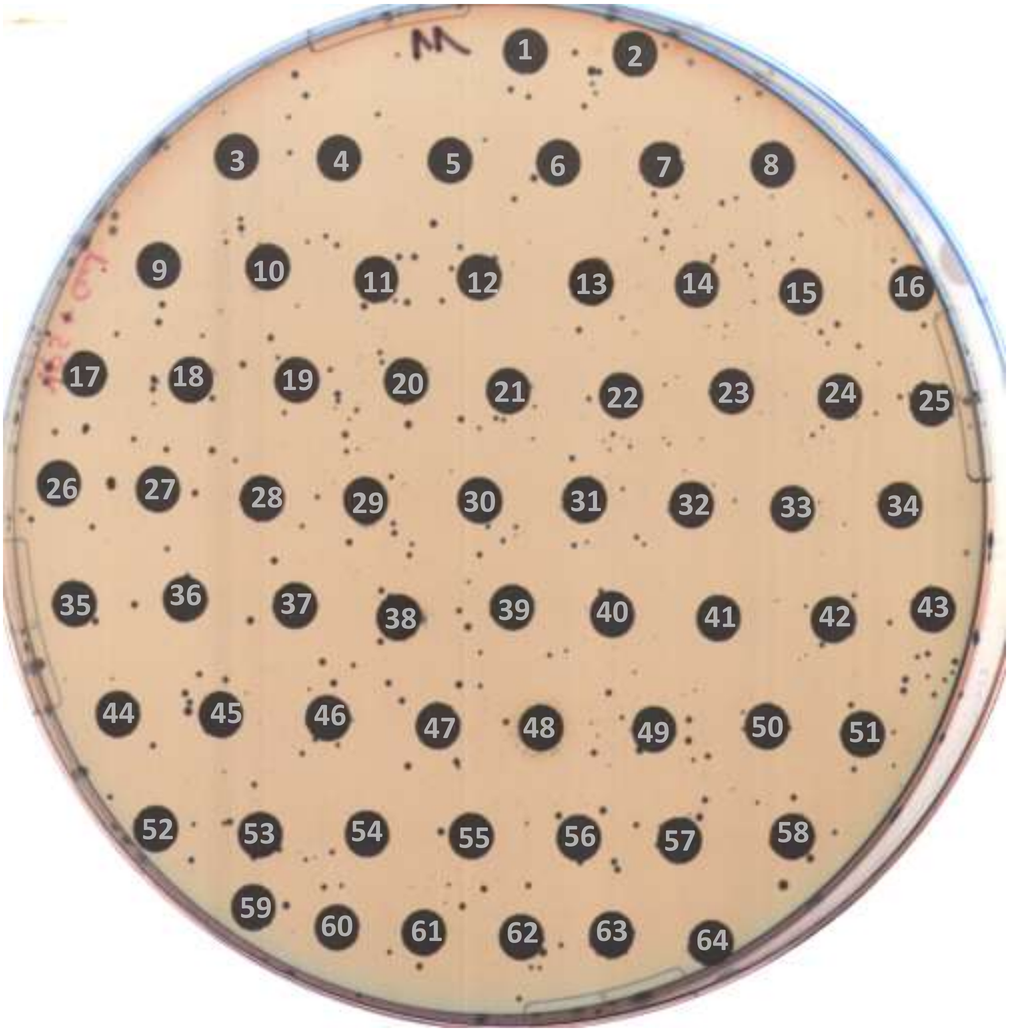

48 h

### PLATE 11

| no. | compound | no. | compound | no. | compound |
|---|---|---|---|---|---|
| - | - | 22 | Ertugliflozin L-pyroglutamic acid | 44 | Deoxycorticosterone acetate |
| 1 | Rucaparib (phosphate) | 23 | Flucytosine | 45 | Ramelteon |
| 2 | Sulfisoxazole | 24 | Hydroxyfasudil | 46 | Iopanoic acid |
| 3 | Esaxerenone | 25 | Trimetazidine (dihydrochloride) | 47 | Entacapone |
| 4 | Taurochenodeoxycholic acid | 26 | Carbetapentane (citrate) | 48 | Ciclopirox (olamine) |
| 5 | Erlotinib (Hydrochloride) | 27 | Perindopril (erbumine) | 49 | S-(+)-Ketoprofen |
| 6 | Eplerenone | 28 | Stiripentol | 50 | Calcium dobesilate |
| 7 | Cevimeline (hydrochloride hemihydrate) | 29 | Rifampicin | 51 | Imatinib (Mesylate) |
| 8 | Amlodipine | 30 | Gemfibrozil | 52 | Cinnarizine |
| 9 | Cefsulodin (sodium) | 31 | Butylphthalide | 53 | Eletriptan (hydrobromide) |
| 10 | Osimertinib | 32 | Naloxegol (oxalate) | 54 | Dofetilide |
| 11 | Idoxuridine | 33 | Doxazosin (mesylate) | 55 | Pirmenol (hydrochloride) |
| 12 | Anagliptin | 34 | Chlorhexidine | 56 | Fesoterodine (fumarate) |
| 13 | Flupirtine (Maleate) | 35 | Chlorzoxazone | 57 | Sulfasalazine |
| 14 | Dexamethasone acetate | 36 | Clofoctol | 58 | Tegafur |
| 15 | Trapidil | 37 | Methicillin (sodium salt) | 59 | Loxoprofen |
| 16 | Phenylephrine (hydrochloride) | 38 | Methacycline (hydrochloride) | 60 | Setiptiline (maleate) |
| 17 | Josamycin | 39 | Drofenine (hydrochloride) | 61 | Ibandronate (Sodium Monohydrate) |
| 18 | Alogliptin (Benzoate) | 40 | Pergolide (mesylate) | 62 | Ketotifen (fumarate) |
| 19 | Benidipine (hydrochloride) | 41 | Lenvatinib | 63 | Delavirdine (mesylate) |
| 20 | Meticrane | 42 | Demecarium Bromide | 64 | Cinepazide (Maleate) |
| 21 | Metolazone | 43 | Oxolamine (citrate) | - | - |

PLATE 12

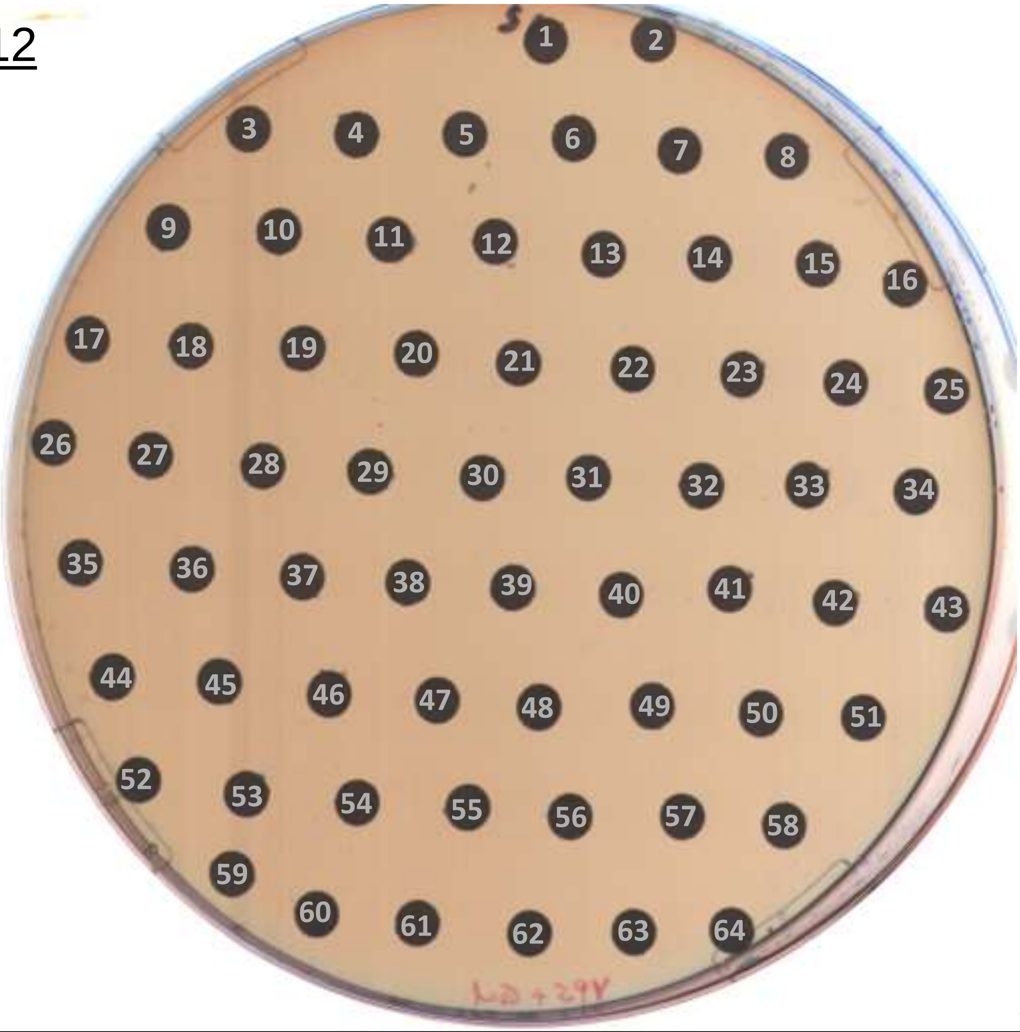

24 h

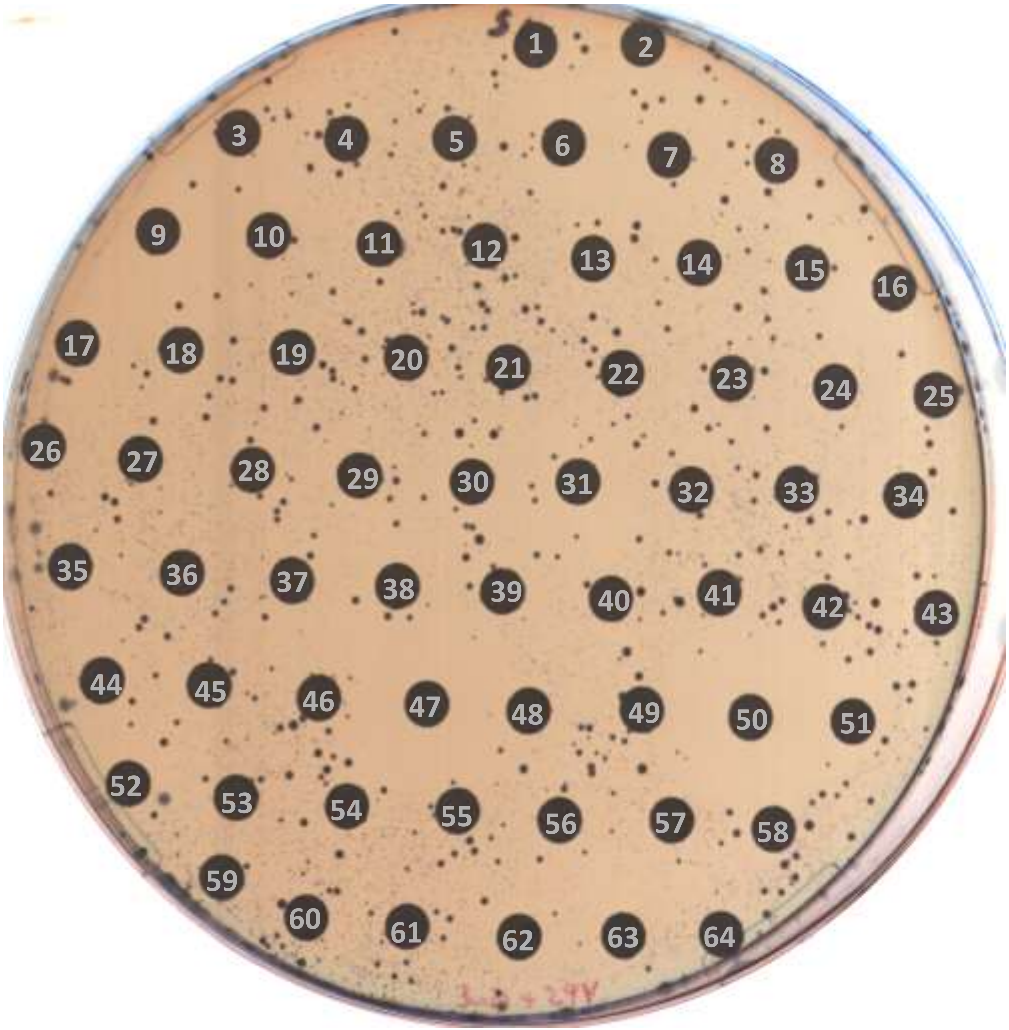

48 h

### PLATE 12

| no. | compound | no. | compound | no. | compound |
|---|---|---|---|---|---|
| - | - | 22 | Ledipasvir (acetone) | 44 | Talipexole dihydrochloride |
| 1 | Telbivudine | 23 | Retinoic acid | 45 | Bosentan |
| 2 | Etodolac | 24 | Urea | 46 | Indapamide |
| 3 | Lercanidipine (hydrochloride) | 25 | AZD7545 | 47 | Disulfiram |
| 4 | Cefixime | 26 | Temocapril (hydrochloride) | 48 | Rivaroxaban |
| 5 | Tamibarotene | 27 | Silodosin | 49 | Lidocaine (hydrochloride) |
| 6 | Telotristat etiprate | 28 | Dronedarone | 50 | Bifonazole |
| 7 | Safinamide | 29 | Bosutinib | 51 | Griseofulvin |
| 8 | Levamisole (hydrochloride) | 30 | Fomepizole | 52 | Amodiaquine (dihydrochloride dihydrate) |
| 9 | Afatinib (dimaleate) | 31 | Azelastine (hydrochloride) | 53 | Fertirelin |
| 10 | Ronidazole | 32 | Methotrexate | 54 | Cisapride |
| 11 | Prilocaine | 33 | Entecavir (monohydrate) | 55 | Danoprevir |
| 12 | Cysteamine hydrochloride | 34 | Suprofen | 56 | Amlexanox |
| 13 | Nifuroxazide | 35 | Antipyrine | 57 | Nicotinamide |
| 14 | Doxofylline | 36 | Etravirine | 58 | Diphenhydramine (hydrochloride) |
| 15 | Nizatidine | 37 | Tolterodine (tartrate) | 59 | Ganciclovir |
| 16 | Bromhexine (hydrochloride) | 38 | Budesonide | 60 | Sulfadoxine |
| 17 | Thiabendazole | 39 | Cefdinir | 61 | Vemurafenib |
| 18 | Setiptiline | 40 | Ranitidine (hydrochloride) | 62 | Anisindione |
| 19 | Pancuronium (dibromide) | 41 | Tiagabine (hydrochloride) | 63 | Levosulpiride |
| 20 | Aztreonam | 42 | Furazolidone | 64 | Lenalidomide (hemihydrate) |
| 21 | Aniracetam | 43 | Mefenamic acid | - | - |

PLATE13

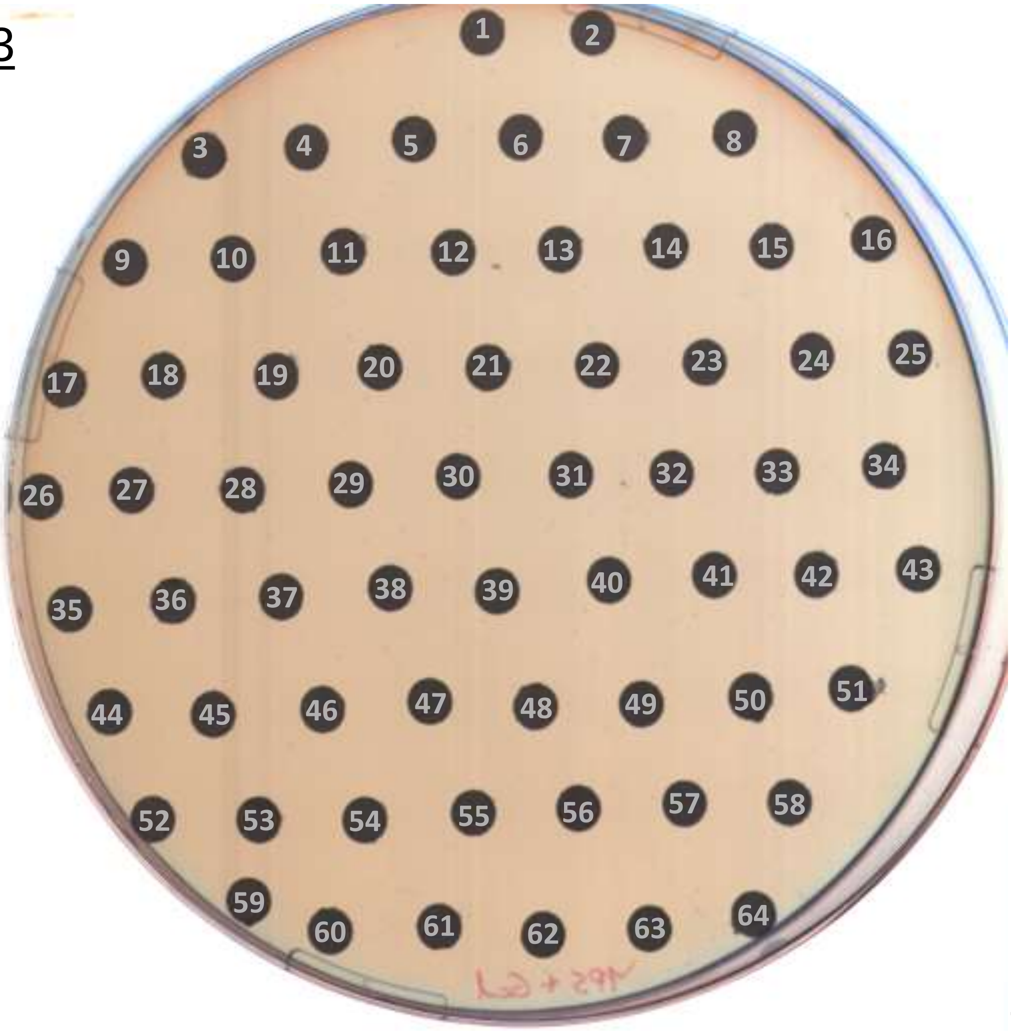

24 h

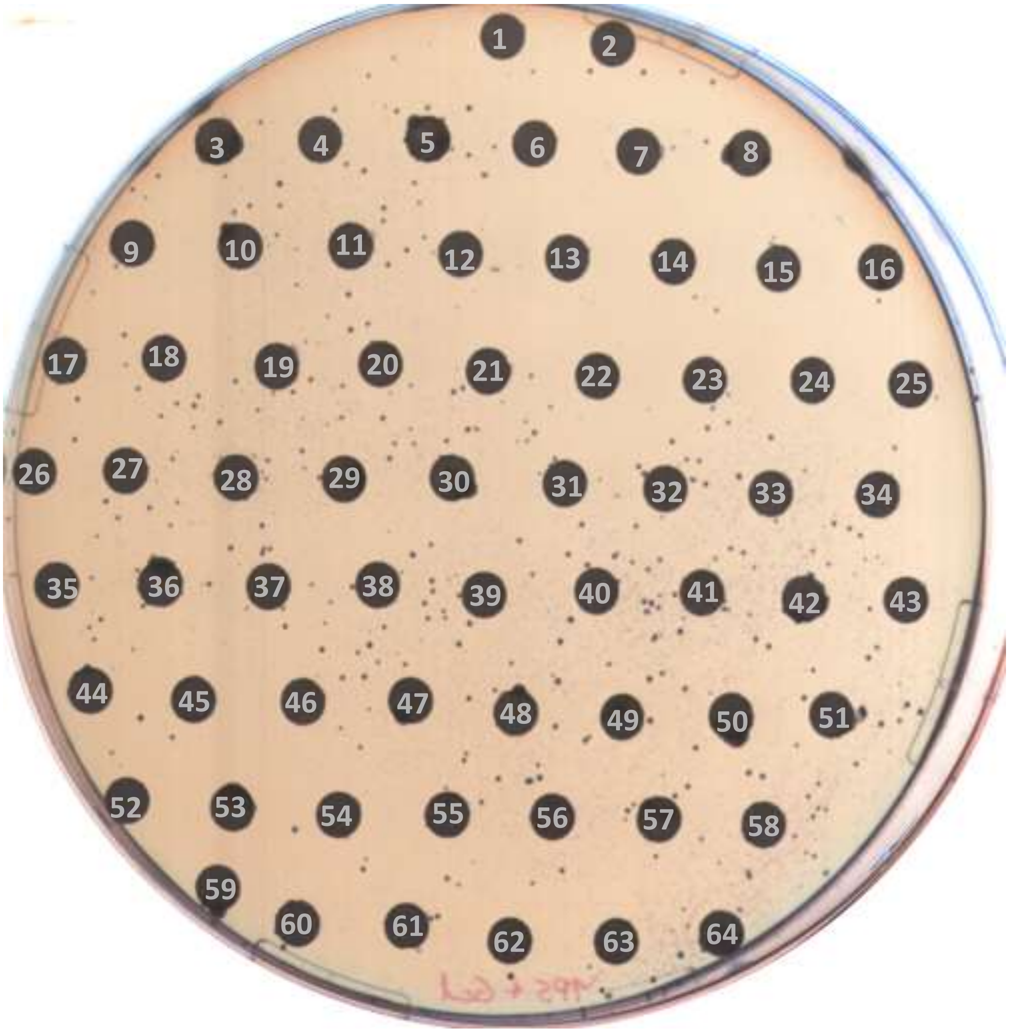

48 h

### PLATE13

| no. | compound | no. | compound | no. | compound |
|---|---|---|---|---|---|
| - | - | 22 | Magnolol | 44 | Loxapine (succinate) |
| 1 | Solifenacin (hydrochloride) | 23 | Avibactam (sodium hydrate) | 45 | Pizotifen |
| 2 | Erythromycin | 24 | Ingenol | 46 | Meloxicam |
| 3 | Meclofenoxate (hydrochloride) | 25 | Dutasteride | 47 | Olsalazine (Disodium) |
| 4 | Clindamycin (hydrochloride) | 26 | Saquinavir (Mesylate) | 48 | Dapoxetine (hydrochloride) |
| 5 | Albendazole | 27 | Aminohippurate (sodium) | 49 | Chlorthalidone |
| 6 | Fabomotizole (hydrochloride) | 28 | Aclacinomycin A hydrochloride | 50 | Ceftizoxime |
| 7 | Leflunomide | 29 | Triamcinolone (acetonide) | 51 | Ulipristal acetate |
| 8 | Nitroprusside (disodium dihydrate) | 30 | Teneligliptin (hydrobromide) | 52 | Ibudilast |
| 9 | Chlorpropamide | 31 | Benorilate | 53 | Nelfinavir |
| 10 | Exemestane | 32 | Nelfinavir (Mesylate) | 54 | Mepivacaine (hydrochloride) |
| 11 | Bephenium (hydroxynaphthoate) | 33 | Dabrafenib | 55 | Propylthiouracil |
| 12 | Voglibose | 34 | Amoxicillin (trihydrate) | 56 | Epirubicin (hydrochloride) |
| 13 | Voriconazole | 35 | Zileuton | 57 | Tiratricol |
| 14 | Topiroxostat | 36 | Nimesulide | 58 | Diclofenac |
| 15 | Lamotrigine | 37 | Nitrofurazone | 59 | (-)-Menthol |
| 16 | Pyrantel (pamoate) | 38 | Oxacillin (sodium monohydrate) | 60 | Prednisolone |
| 17 | Fosamprenavir | 39 | Pemirolast (potassium) | 61 | Capecitabine |
| 18 | Nitrofurantoin | 40 | Triamcinolone | 62 | Cabozantinib (S-malate) |
| 19 | Irinotecan | 41 | Tafamidis | 63 | Rilpivirine |
| 20 | Dronedarone (Hydrochloride) | 42 | Bufexamac | 64 | Decamethonium (Bromide) |
| 21 | Moxidectin | 43 | Estriol | - | - |

PLATE 14

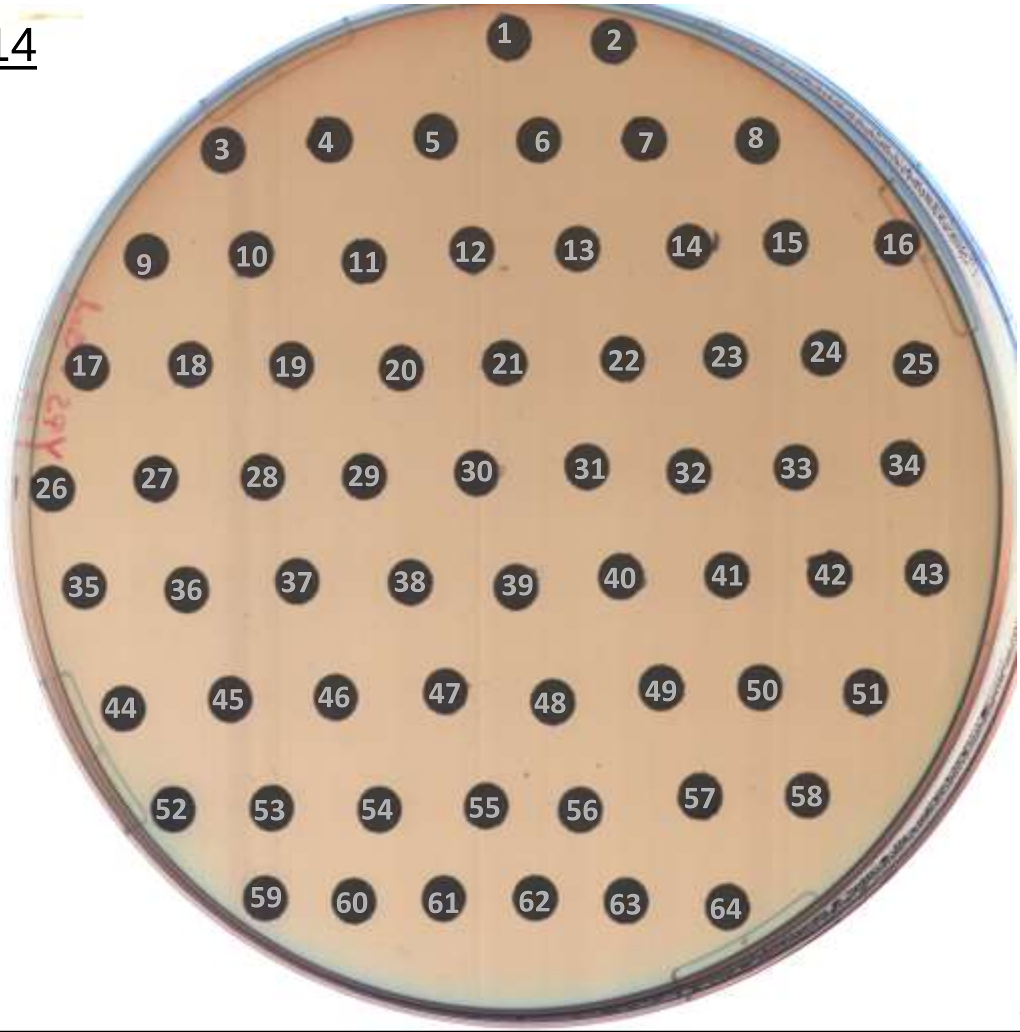

24 h

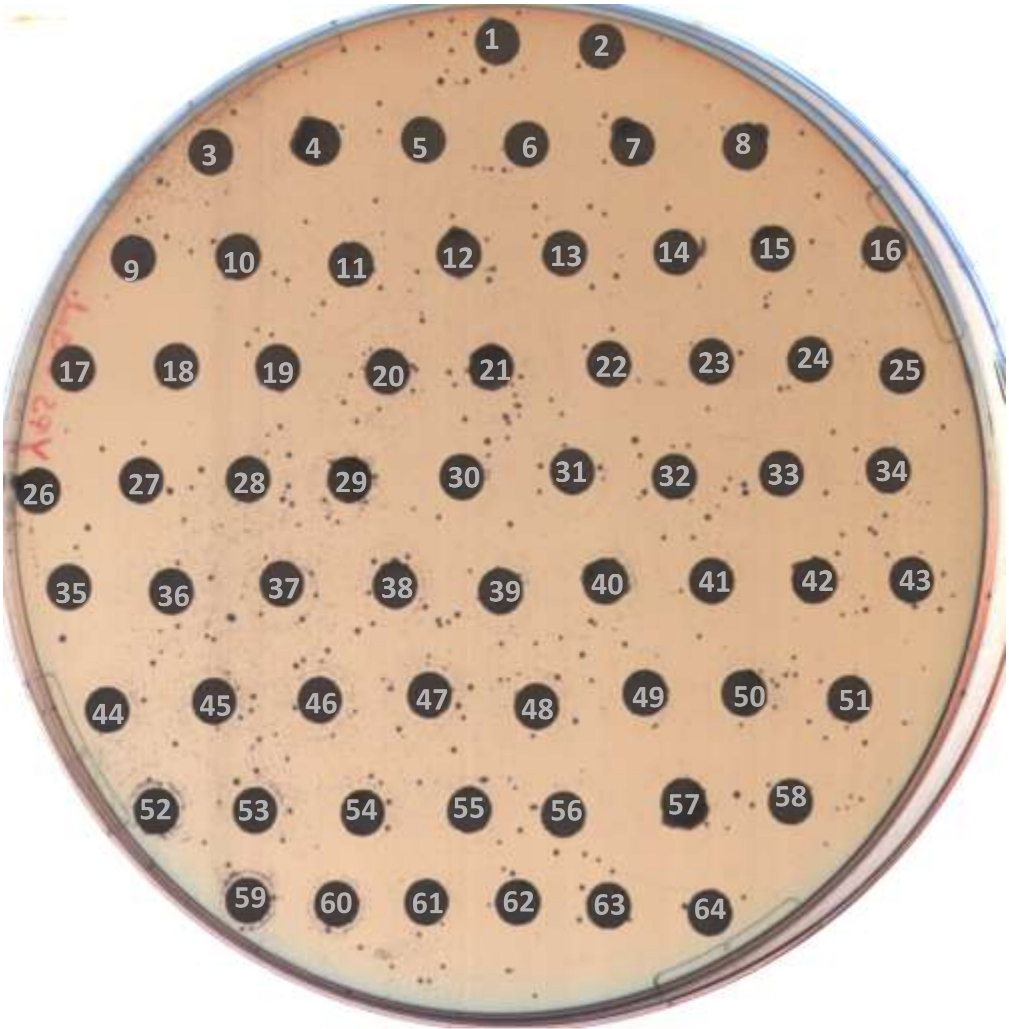

48 h

### PLATE 14

| no. | compound | no. | compound | no. | compound |
|---|---|---|---|---|---|
| - | - | 22 | Etretinate | 44 | Phenylbutazone |
| 1 | Tropisetron (Hydrochloride) | 23 | Triamterene | 45 | Ribavirin |
| 2 | Cefoperazone (sodium salt) | 24 | Proguanil | 46 | Lafutidine |
| 3 | Pamabrom | 25 | Flumazenil | 47 | Olmesartan medoxomil |
| 4 | Pazopanib | 26 | Ketanserin | 48 | Enoxacin (hydrate) |
| 5 | Ledipasvir | 27 | Trichlormethiazide | 49 | Deferasirox |
| 6 | Amprenavir | 28 | Gabapentin | 50 | Drospirenone |
| 7 | Cefoxitin (sodium) | 29 | Levobupivacaine (hydrochloride) | 51 | Macitentan |
| 8 | Etoposide | 30 | Enzalutamide | 52 | Mitoxantrone (dihydrochloride) |
| 9 | Ciclesonide | 31 | Topiramate | 53 | Domperidone |
| 10 | L-Hyoscyamine | 32 | Favipiravir | 54 | Amlodipine (maleate) |
| 11 | Antazoline (hydrochloride) | 33 | Methoxsalen | 55 | Cefmenoxime (hydrochloride) |
| 12 | Cefmetazole (sodium) | 34 | Fimasartan | 56 | Tipranavir |
| 13 | Zalcitabine | 35 | Buflomedil (hydrochloride) | 57 | Halobetasol (propionate) |
| 14 | Eicosapentaenoic Acid | 36 | Aprepitant | 58 | Ezetimibe |
| 15 | Menadione bisulfite (sodium) | 37 | Fluvoxamine (maleate) | 59 | Tropisetron |
| 16 | Theobromine | 38 | Metronidazole | 60 | Simeprevir |
| 17 | Efavirenz | 39 | Enalaprilat (dihydrate) | 61 | Enalapril (maleate) |
| 18 | Terconazole | 40 | Trilostane | 62 | Lornoxicam |
| 19 | Oxybenzone | 41 | Ivabradine (hydrochloride) | 63 | Aceclofenac |
| 20 | Memantine (hydrochloride) | 42 | Ziprasidone (hydrochloride monohydrate) | 64 | Talazoparib |
| 21 | Spironolactone | 43 | Sulfamethoxazole | - | - |

PLATE 15

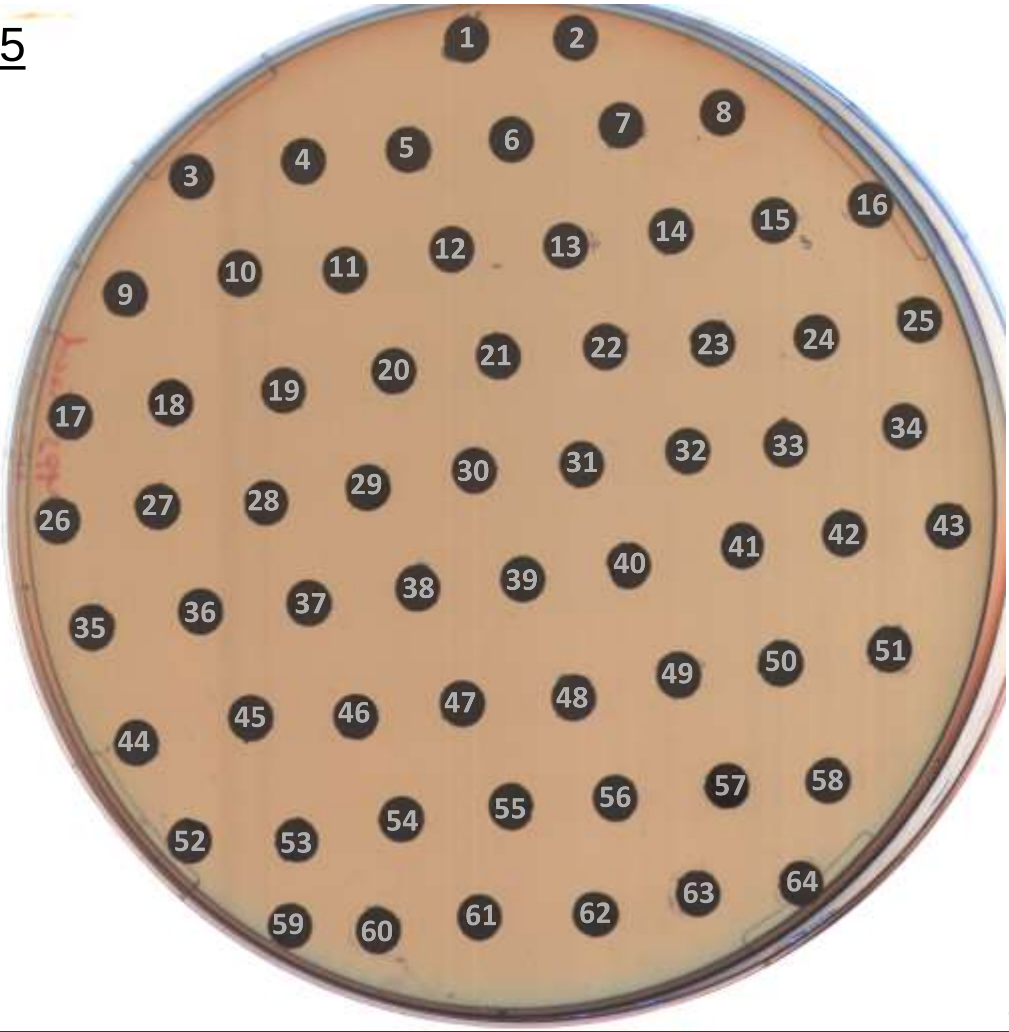

24 h

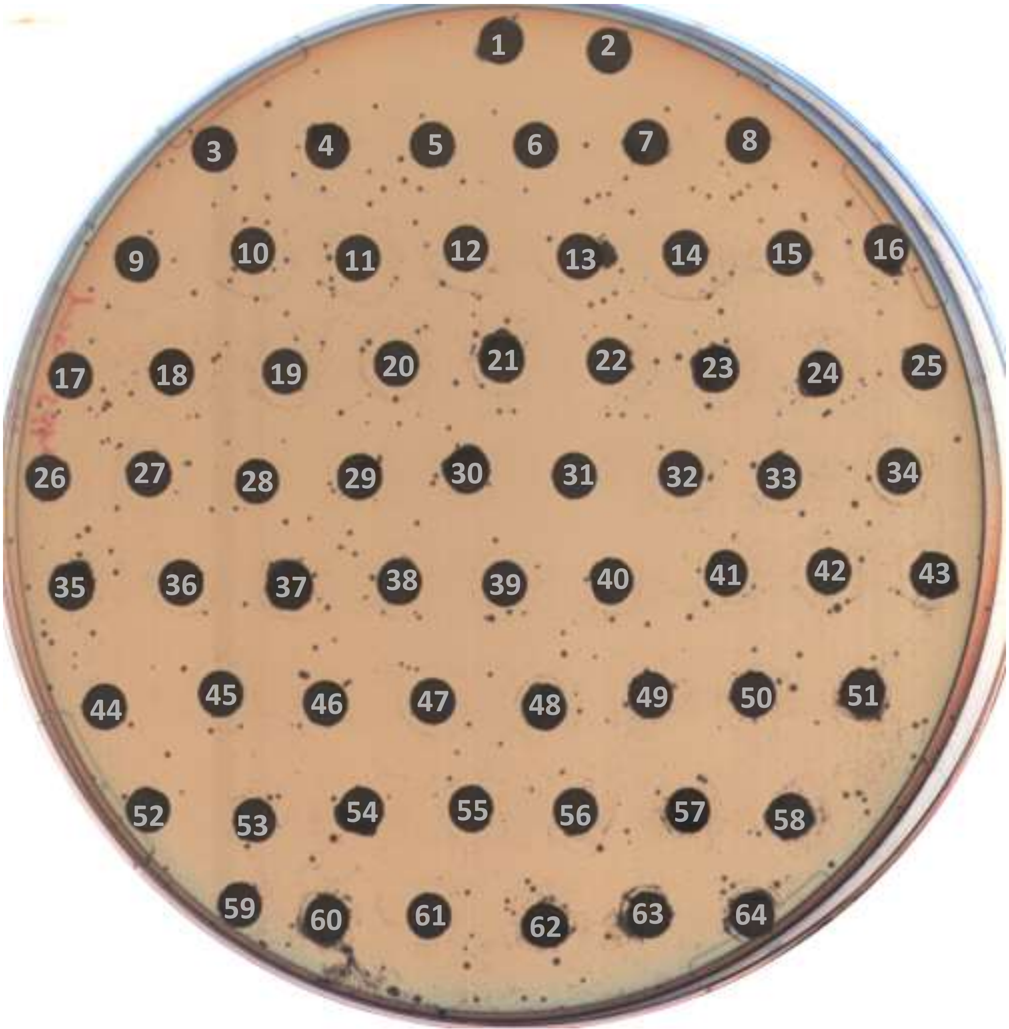

48 h

### PLATE 15

| no. | compound | no. | compound | no. | compound |
|---|---|---|---|---|---|
| - | - | 22 | Travoprost | 44 | Captopril |
| 1 | Apremilast | 23 | Dolutegravir | 45 | Rufinamide |
| 2 | Rosuvastatin (Calcium) | 24 | Biperiden (Hydrochloride) | 46 | Mycophenolate Mofetil |
| 3 | Manidipine (dihydrochloride) | 25 | Batilol | 47 | Amphotericin B |
| 4 | Cromolyn (sodium) | 26 | Halcinonide | 48 | Trelagliptin |
| 5 | Duloxetine (hydrochloride) | 27 | Deferasirox (Fe3+ chelate) | 49 | Temozolomide |
| 6 | Rotigotine | 28 | Pentamidine (isethionate) | 50 | Tofacitinib (citrate) |
| 7 | Cyproheptadine (hydrochloride) | 29 | Adenosine | 51 | Pazopanib (Hydrochloride) |
| 8 | Doxorubicin (hydrochloride) | 30 | Doxylamine (succinate) | 52 | Imidafenacin |
| 9 | Gimeracil | 31 | Domiphen (bromide) | 53 | N-Acetylprocainamide |
| 10 | Flurbiprofen | 32 | Permethrin | 54 | Vancomycin (hydrochloride) |
| 11 | Miglitol | 33 | γ-Oryzanol | 55 | Bergenin |
| 12 | Sotalol (hydrochloride) | 34 | Etamivan | 56 | Nicorandil |
| 13 | Mirodenafil (dihydrochloride) | 35 | Fosfluconazole | 57 | Tigecycline |
| 14 | Hydroxyurea | 36 | Olaparib | 58 | Phthalylsulfacetamide |
| 15 | Flecainide (acetate) | 37 | Rutin | 59 | Cyproheptadine (hydrochloride sesquihydrate) |
| 16 | Fluphenazine decanoate | 38 | Roflumilast | 60 | Trimipramine (maleate) |
| 17 | Trametinib | 39 | Cetirizine (dihydrochloride) | 61 | Tenofovir alafenamide hemifumarate |
| 18 | Norepinephrine | 40 | Omeprazole | 62 | Hydrocortisone 17-butyrate |
| 19 | Stavudine | 41 | Teriflunomide | 63 | Desvenlafaxine |
| 20 | Tedizolid (phosphate) | 42 | Canagliflozin | 64 | Reserpine (hydrochloride) |
| 21 | Triflusal | 43 | Medetomidine (hydrochloride) | - | - |

<u>PLATE 16</u>

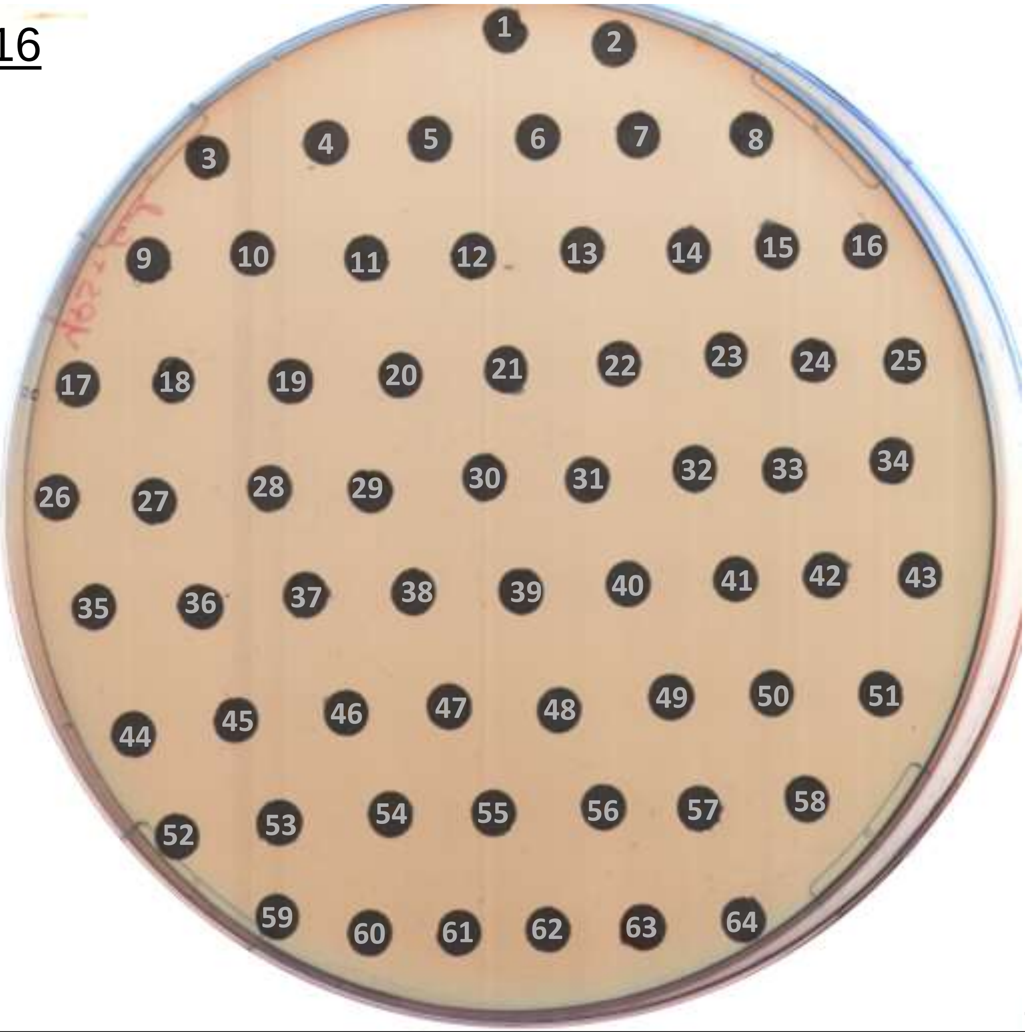

24 h

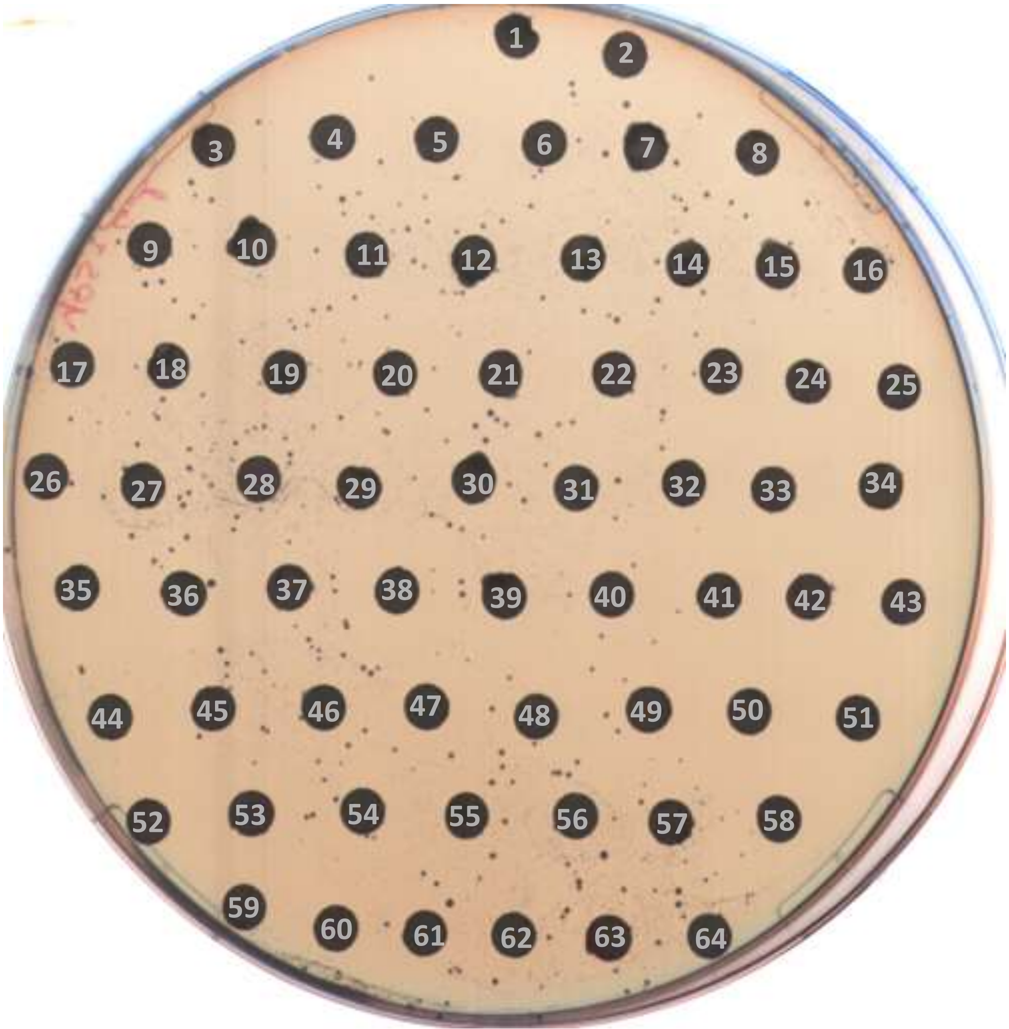

48 h

### PLATE 16

| no. | compound | no. | compound | no. | compound |
|---|---|---|---|---|---|
| - | - | 22 | Quinine | 44 | Carglumic Acid |
| 1 | Valproic acid | 23 | Nystatin | 45 | Troxerutin |
| 2 | Phenprocoumon | 24 | Gallic acid | 46 | Asunaprevir |
| 3 | Visomitin | 25 | Darunavir | 47 | (Z)-Capsaicin |
| 4 | Proxyphylline | 26 | Bexarotene | 48 | Pargyline (hydrochloride) |
| 5 | Fasudil (Hydrochloride) | 27 | Difluprednate | 49 | Betamethasone dipropionate |
| 6 | Latrepirdine (dihydrochloride) | 28 | Lovastatin | 50 | Allantoin |
| 7 | Valdecoxib | 29 | Sonidegib (diphosphate) | 51 | Temsirolimus |
| 8 | Lasofoxifene (Tartrate) | 30 | Levofloxacin | 52 | Solifenacin |
| 9 | Bortezomib | 31 | Fursultiamine | 53 | Tolterodine |
| 10 | Rofecoxib | 32 | Ertapenem sodium | 54 | Quinidine hydrochloride monohydrate |
| 11 | Carprofen | 33 | Pasiniazid | 55 | Sulfisomidin |
| 12 | Cabozantinib | 34 | Isoconazole (nitrate) | 56 | Sulfamerazine |
| 13 | (S)-10-Hydroxycamptothecin | 35 | Davercin | 57 | Propantheline (bromide) |
| 14 | Tadalafil | 36 | Ombitasvir | 58 | 4-Methylumbelliferone |
| 15 | Nimustine (hydrochloride) | 37 | Piperidolate (hydrochloride) | 59 | Desipramine hydrochloride |
| 16 | 6-Mercaptopurine hydrate | 38 | Deferoxamine (mesylate) | 60 | Guaiacol |
| 17 | Tenofovir (Disoproxil) | 39 | Velpatasvir | 61 | Nalidixic acid |
| 18 | Paeonol | 40 | Tulobuterol (hydrochloride) | 62 | Imipramine (hydrochloride) |
| 19 | Hydrocortisone | 41 | Lorlatinib | 63 | Valrubicin |
| 20 | Ibuprofen piconol | 42 | Sitagliptin (phosphate monohydrate) | 64 | Cephalothin (sodium) |
| 21 | Sapropterin (dihydrochloride) | 43 | Rotundine | - | - |

PLATE 17

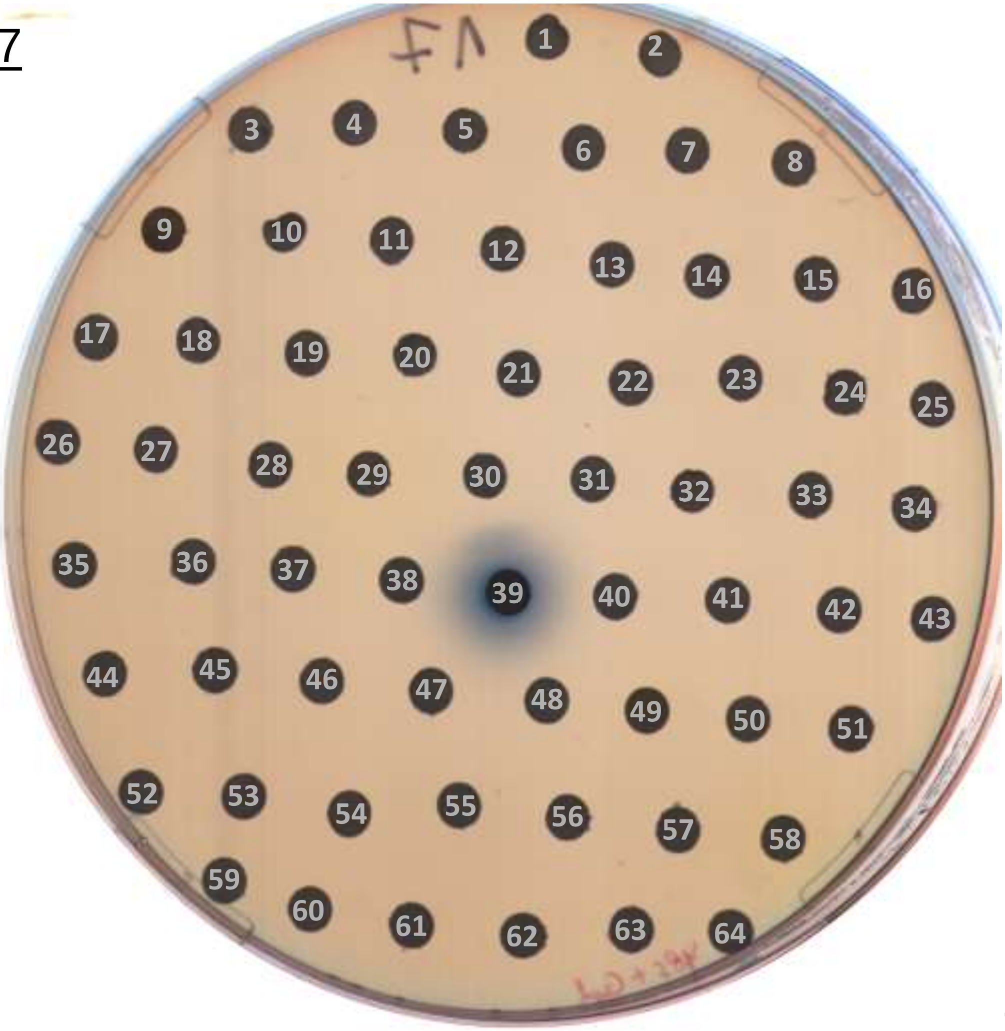

24 h

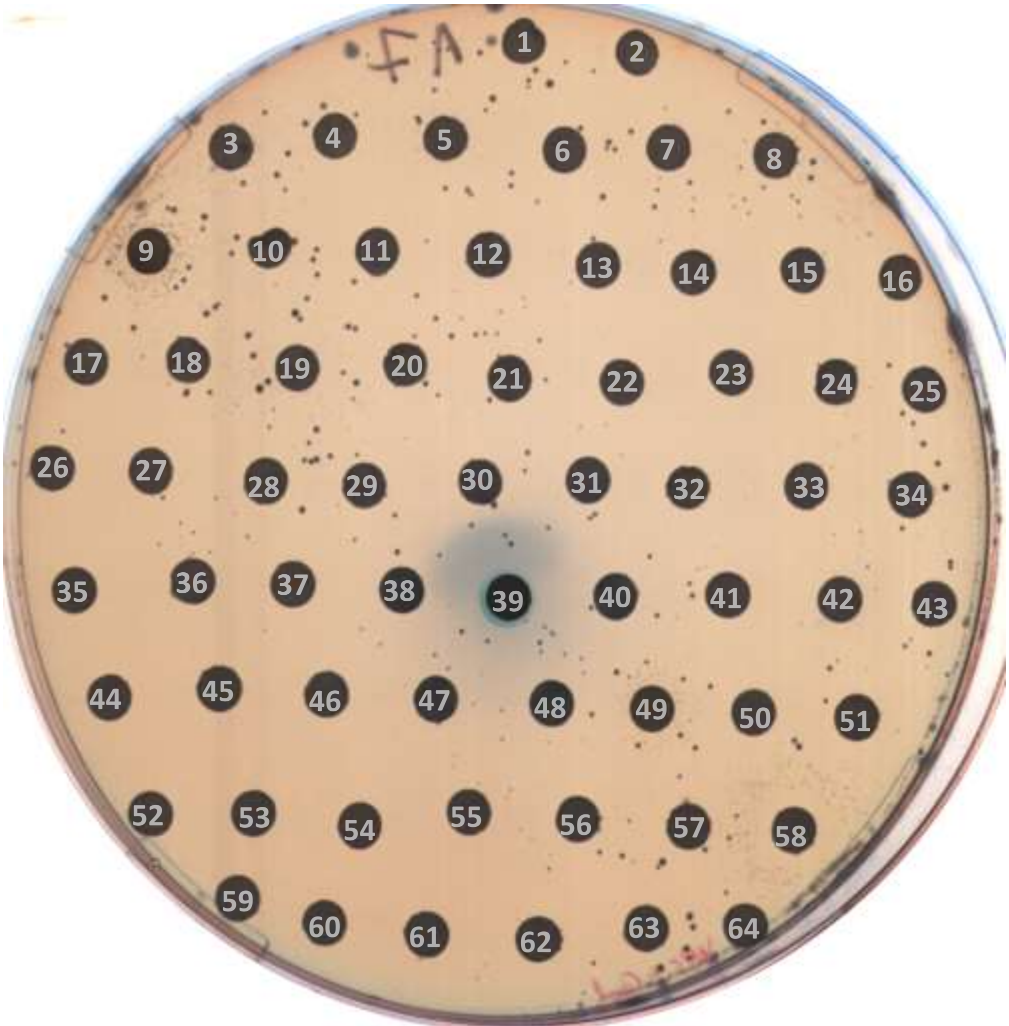

48 h

### PLATE 17

| no. | compound | no. | compound | no. | compound |
|---|---|---|---|---|---|
| - | - | 22 | Choline (chloride) | 44 | Bisoctrizole |
| 1 | Reserpine | 23 | Triclosan | 45 | Sodium tauroglycocholate |
| 2 | Amlodipine (besylate) | 24 | Chromocarb | 46 | Acetohydroxamic acid |
| 3 | Methylprednisolone succinate | 25 | Zaltoprofen | 47 | Fludrocortisone acetate |
| 4 | Andrographolide | 26 | Carbimazole | 48 | Vitamin D2 |
| 5 | Carmustine | 27 | Bivalirudin (TFA) | 49 | Ethacridine (lactate monohydrate) |
| 6 | Lodoxamide (tromethamine) | 28 | Ritodrine (hydrochloride) | 50 | Glasdegib |
| 7 | Nefopam (hydrochloride) | 29 | Carbazochrome (sodium sulfonate) | 51 | Cobicistat |
| 8 | Pazufloxacin (mesylate) | 30 | Ribociclib | 52 | Isavuconazole |
| 9 | Ethacridine (lactate) | 31 | Metadoxine | 53 | Fosinopril (sodium) |
| 10 | Talc | 32 | Lactitol (monohydrate) | 54 | Guanfacine (hydrochloride) |
| 11 | Lincomycin (hydrochloride monohydrate) | 33 | Spectinomycin (dihydrochloride pentahydrate) | 55 | Fenbufen |
| 12 | Homoharringtonine | 34 | Salsalate | 56 | Erythromycin Ethylsuccinate |
| 13 | Dimetridazole | 35 | Elbasvir | 57 | Caffeic acid |
| 14 | Efonidipine (hydrochloride monoethanolate) | 36 | Teprenone | 58 | Berberine (chloride hydrate) |
| 15 | Dapagliflozin | 37 | Chlorphenoxamine | 59 | Glycyrrhizic acid |
| 16 | Toremifene (citrate) | 38 | Valacyclovir (hydrochloride) | 60 | Everolimus |
| 17 | Alpelisib | 39 | Evans Blue | 61 | Fluralaner |
| 18 | Ranolazine (dihydrochloride) | 40 | Estramustine (phosphate sodium) | 62 | Avibactam (sodium) |
| 19 | Ceftibuten (dihydrate) | 41 | Racanisodamine | 63 | Ceritinib |
| 20 | Linaclotide | 42 | Crisaborole | 64 | Sucralfate |
| 21 | Spectinomycin (dihydrochloride) | 43 | Tamoxifen | - | - |

<u>PLATE 18</u>

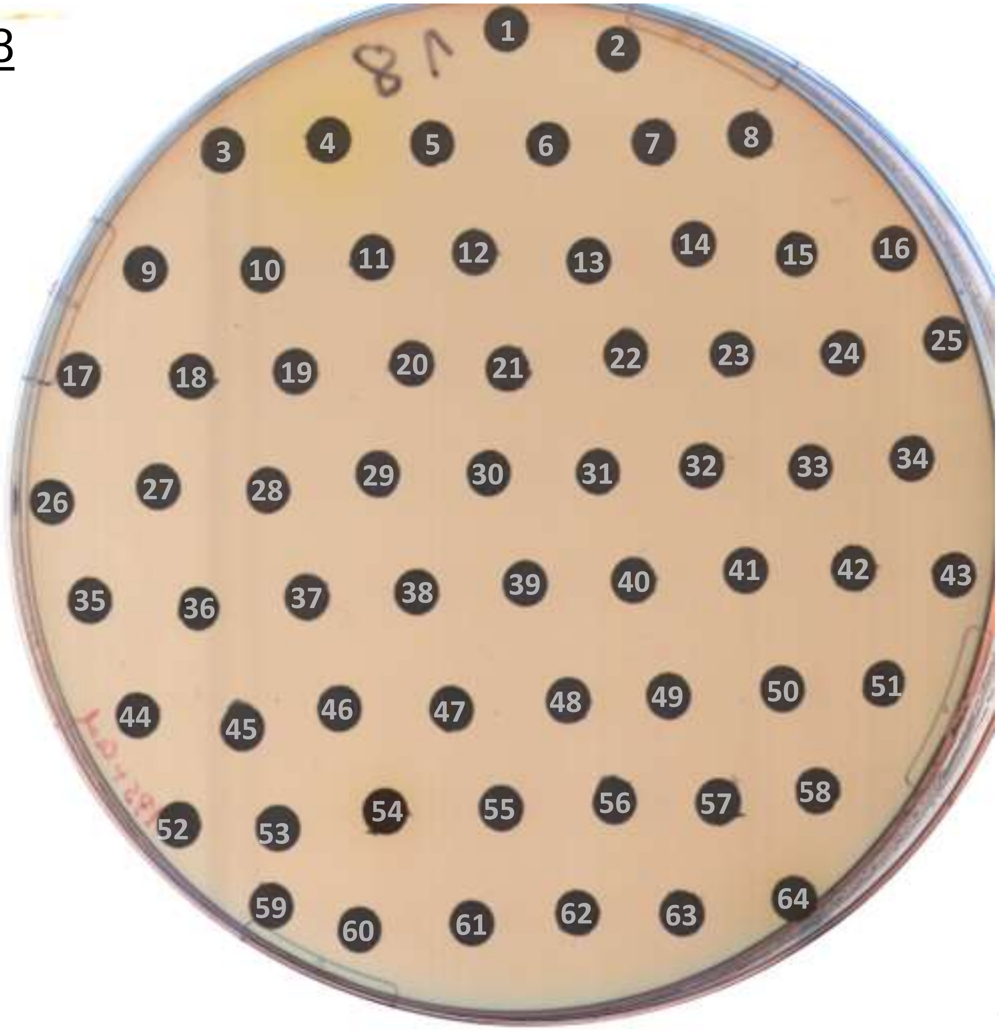

24 h

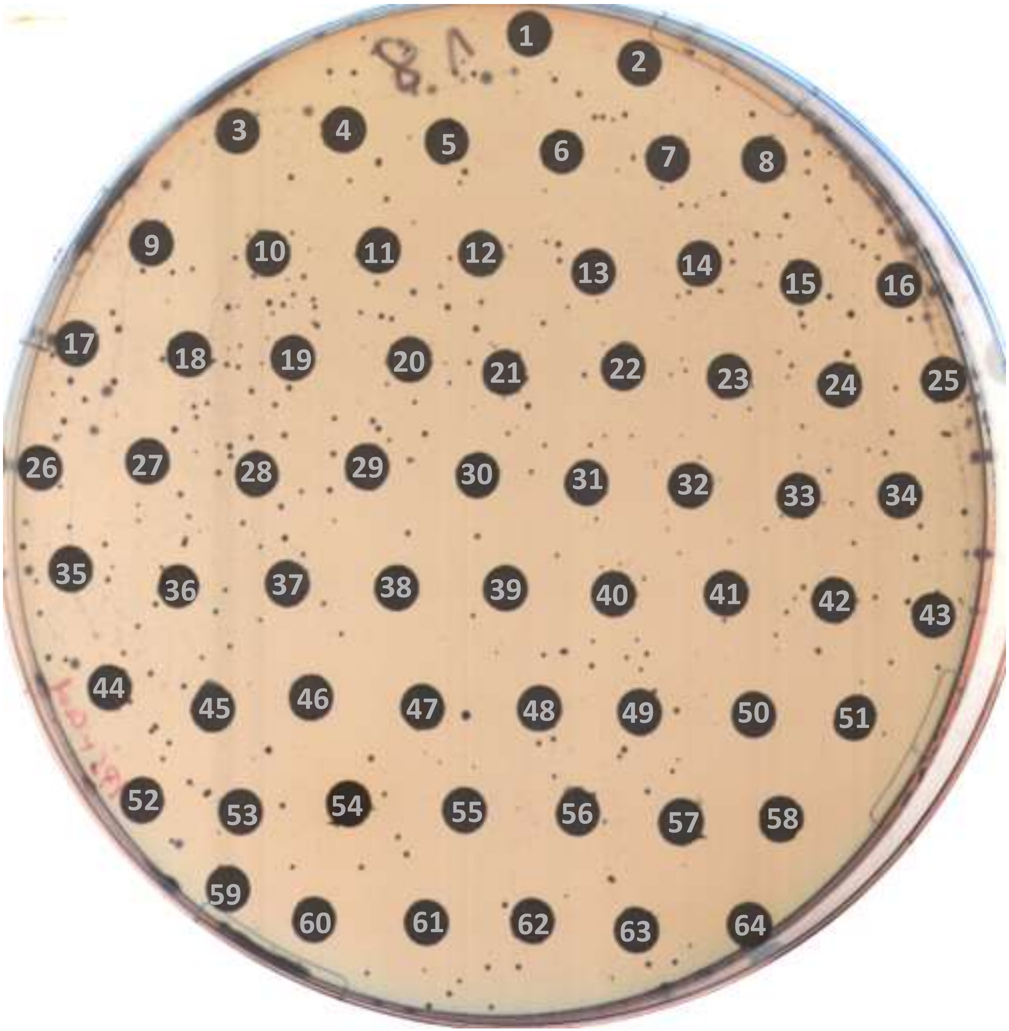

48 h

### PLATE 18

| no. | compound | no. | compound | no. | compound |
|---|---|---|---|---|---|
| - | - | 22 | Delafloxacin (meglumine) | 44 | Acalabrutinib |
| 1 | Omarigliptin | 23 | Dorzolamide (hydrochloride) | 45 | Erdosteine |
| 2 | Acetylspiramycin | 24 | Cefadroxil | 46 | Cholic acid |
| 3 | Hydrocortisone acetate | 25 | Megestrol acetate | 47 | Degarelix |
| 4 | Flavin Adenine Dinucleotide Disodium | 26 | Imrecoxib | 48 | Bosentan (hydrate) |
| 5 | Bremelanotide (Acetate) | 27 | Clebopride (malate) | 49 | Erlotinib |
| 6 | Zanamivir | 28 | Tazobactam | 50 | Iguratimod |
| 7 | Bucladesine (sodium) | 29 | Ursodiol | 51 | Clopidogrel (hydrogen sulfate) |
| 8 | Cefonicid (sodium) | 30 | Praziquantel | 52 | Otilonium (bromide) |
| 9 | Desoxycorticosterone pivalate | 31 | Theophylline | 53 | Galanthamine (hydrobromide) |
| 10 | L-Cystine | 32 | Retapamulin | 54 | Eltrombopag (Olamine) |
| 11 | Triamcinolone hexacetonide | 33 | Azatadine (dimaleate) | 55 | Cyproterone acetate |
| 12 | Pentoxifylline | 34 | Sulbutiamine | 56 | Teriparatide |
| 13 | Daclatasvir (dihydrochloride) | 35 | Adenine | 57 | Gemcitabine |
| 14 | Ceftazidime | 36 | Sofalcone | 58 | Butoconazole (nitrate) |
| 15 | Gefitinib | 37 | Sivelestat | 59 | Taurine |
| 16 | Deferiprone | 38 | Cefepime (Dihydrochloride Monohydrate) | 60 | Diphenmanil (methylsulfate) |
| 17 | Probucol | 39 | Olprinone (Hydrochloride) | 61 | Avobenzone |
| 18 | Daclatasvir | 40 | Desloratadine | 62 | Terlipressin |
| 19 | Escitalopram (oxalate) | 41 | Betahistine (dihydrochloride) | 63 | Heptaminol (hydrochloride) |
| 20 | Ceritinib dihydrochloride | 42 | Moexipril (hydrochloride) | 64 | Tigecycline (tetramesylate) |
| 21 | Penciclovir | 43 | Faropenem daloxate | - | - |

PLATE 19

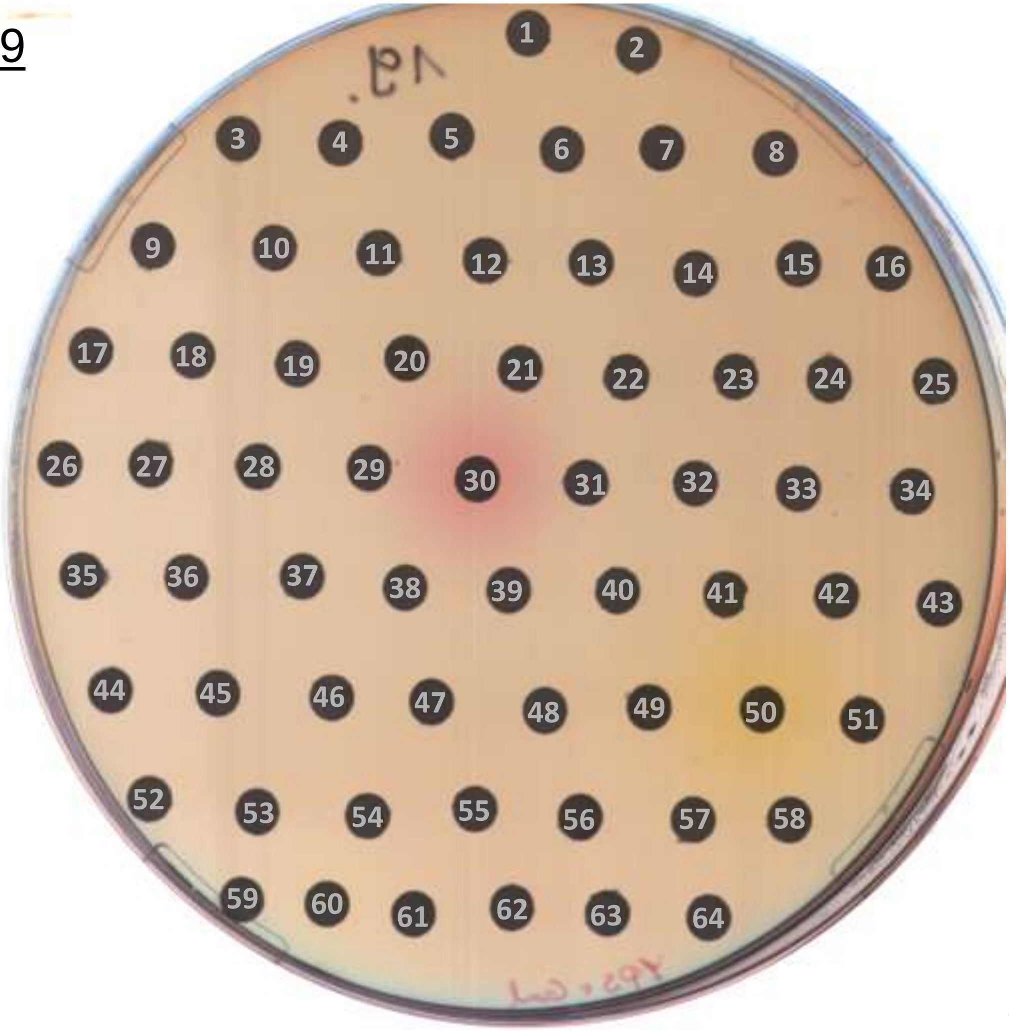

24 h

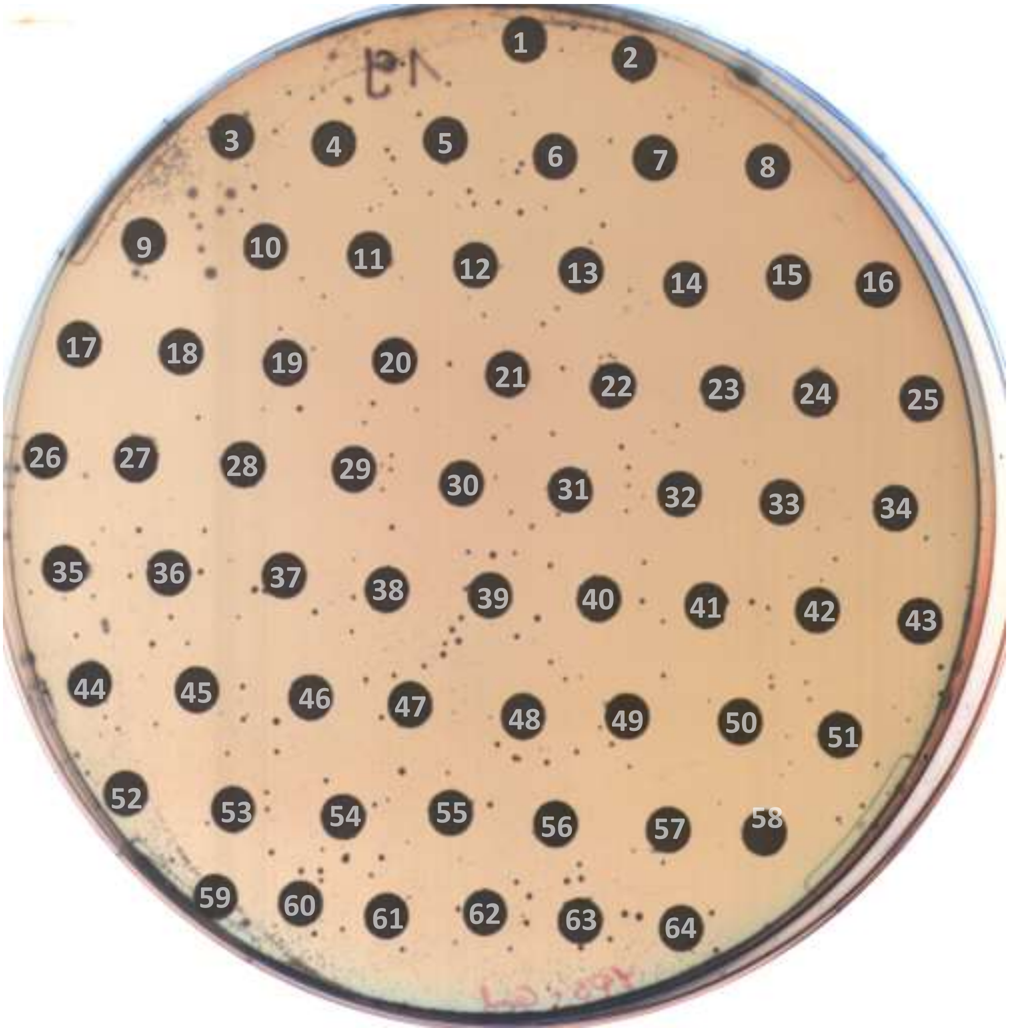

48 h

### PLATE 19

| no. | compound | no. | compound | no. | compound |
|---|---|---|---|---|---|
| - | - | 22 | Norethindrone acetate | 44 | Pioglitazone |
| 1 | Metaxalone | 23 | Solriamfetol | 45 | Abemaciclib (methanesulfonate) |
| 2 | Cladribine | 24 | Darifenacin (hydrobromide) | 46 | Ornipressin |
| 3 | Pyridoxine (hydrochloride) | 25 | Beclometasone dipropionate | 47 | Dabrafenib (Mesylate) |
| 4 | Carbetocin | 26 | Tizanidine (hydrochloride) | 48 | Verapamil (hydrochloride) |
| 5 | Abiraterone acetate | 27 | Sertindole | 49 | Succimer |
| 6 | Grazoprevir | 28 | Nalfurafine (hydrochloride) | 50 | Phenazopyridine (hydrochloride) |
| 7 | Candesartan Cilexetil | 29 | Valemetostat (tosylate) | 51 | Ozenoxacin |
| 8 | Ampiroxicam | 30 | Vitamin B12 | 52 | Losartan |
| 9 | Cobimetinib (racemate) | 31 | Progesterone | 53 | Azilsartan medoxomil |
| 10 | Clozapine | 32 | Desvenlafaxine (succinate hydrate) | 54 | Obeticholic acid |
| 11 | Clemizole (hydrochloride) | 33 | Diclofenac (diethylamine) | 55 | Pexidartinib (hydrochloride) |
| 12 | Montelukast (sodium) | 34 | Bumetanide | 56 | Dalbavancin (hydrochloride) |
| 13 | Histamine | 35 | Asenapine | 57 | Telotristat ethyl |
| 14 | Elagolix sodium | 36 | α-Vitamin E | 58 | Hexylresorcinol |
| 15 | Mycophenolic acid | 37 | Puerarin | 59 | Clofibric acid |
| 16 | Ketoconazole | 38 | i-Inositol | 60 | Balsalazide |
| 17 | Fluvastatin (sodium) | 39 | Sulfabenzamide | 61 | Pralidoxime (chloride) |
| 18 | Framycetin | 40 | Nitrendipine | 62 | Edaravone |
| 19 | Adapalene | 41 | Rucaparib (Camsylate) | 63 | Eprosartan (mesylate) |
| 20 | Furagin | 42 | Tenofovir alafenamide | 64 | Larotrectinib |
| 21 | Etomidate (hydrochloride) | 43 | Pindolol | - | - |

PLATE 20

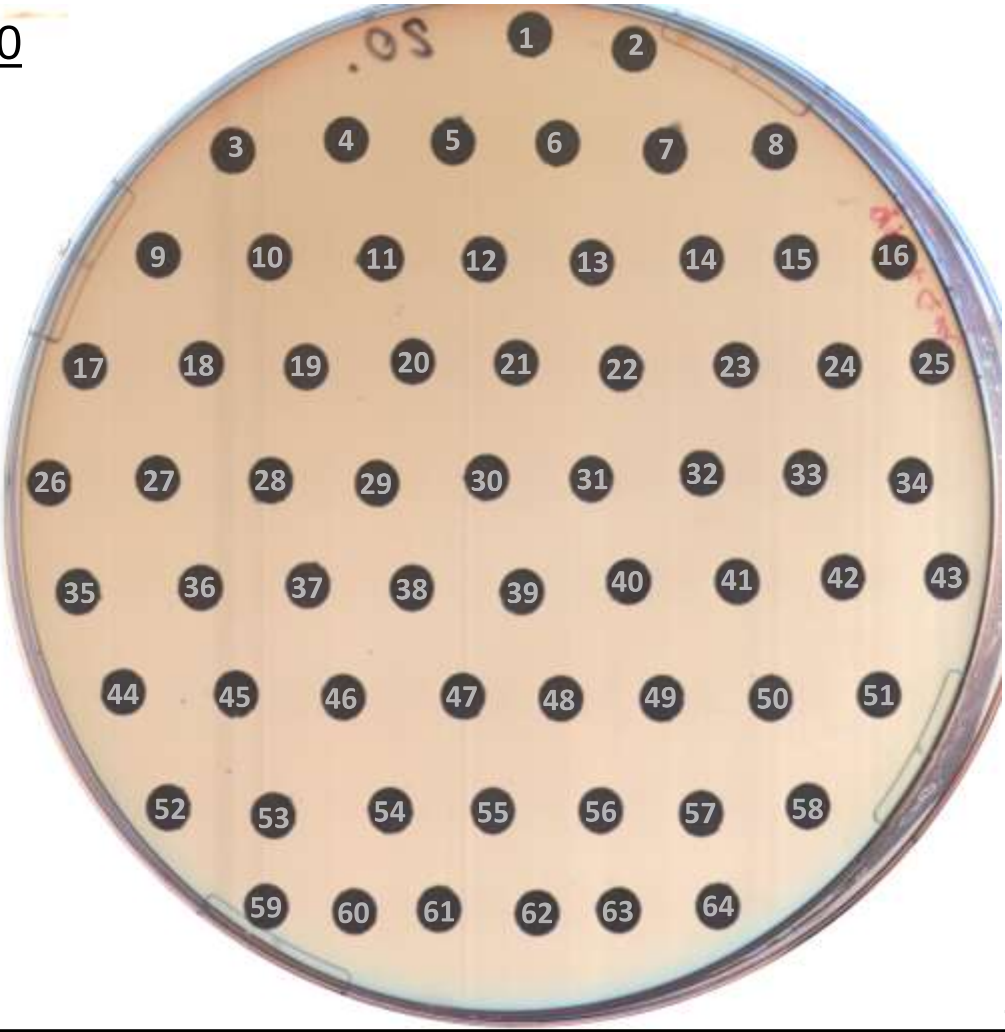

24 h

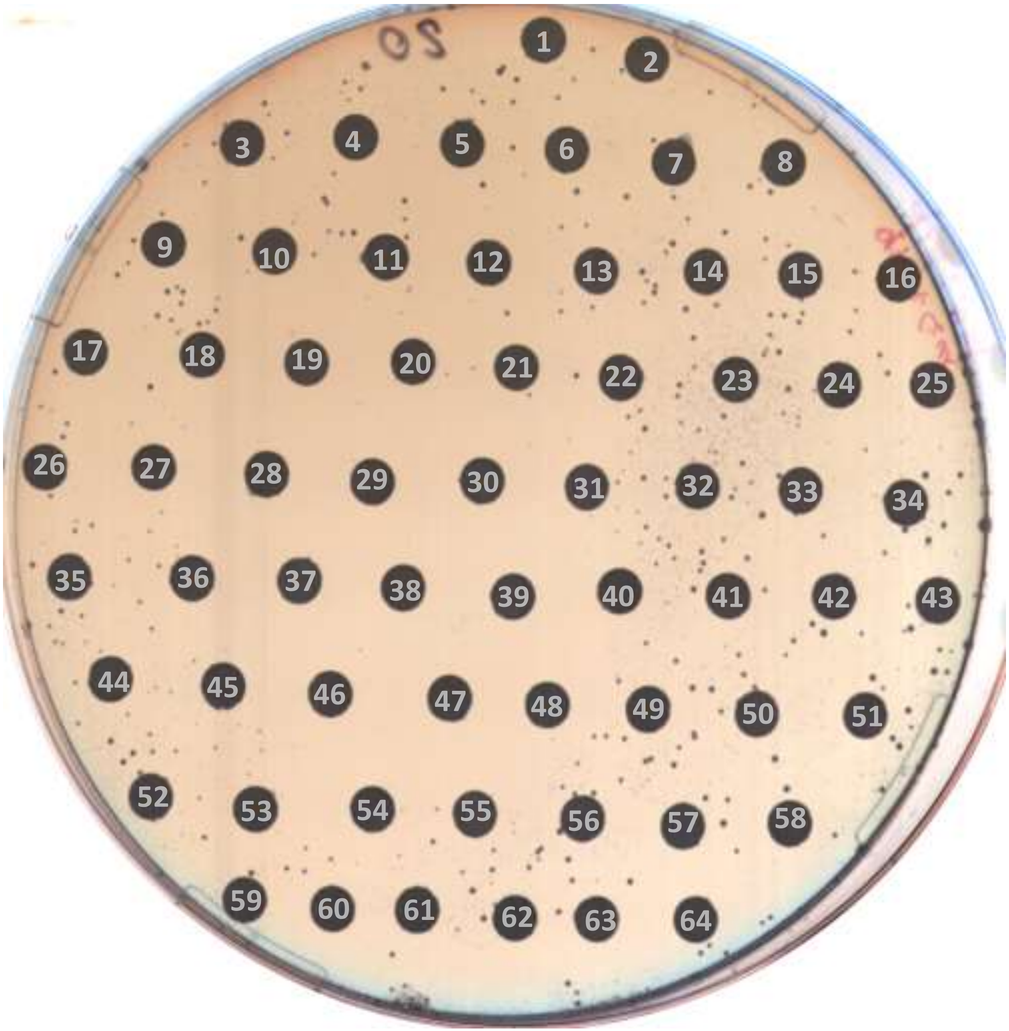

48 h

### PLATE 20

| no. | compound | no. | compound | no. | compound |
|---|---|---|---|---|---|
| - | - | 22 | Uridine | 44 | Dimemorfan (phosphate) |
| 1 | Probenecid | 23 | Fructose | 45 | Alcaftadine |
| 2 | Cilastatin | 24 | Benzoic acid | 46 | Sulbactam |
| 3 | Estradiol benzoate | 25 | Fudosteine | 47 | Sulfacetamide (Sodium) |
| 4 | Crotamiton | 26 | Tilorone (dihydrochloride) | 48 | Doripenem (monohydrate) |
| 5 | Raltegravir | 27 | Cyclandelate | 49 | Cefoselis (sulfate) |
| 6 | Epalrestat | 28 | Lidocaine | 50 | Chlorothiazide |
| 7 | Enasidenib (mesylate) | 29 | Atropine | 51 | Bethanechol (chloride) |
| 8 | Ruxolitinib (phosphate) | 30 | Axitinib | 52 | Atazanavir (sulfate) |
| 9 | Inosine pranobex | 31 | Docosahexaenoic Acid | 53 | Azlocillin (sodium salt) |
| 10 | Tetracycline (hydrochloride) | 32 | Cabazitaxel | 54 | Methazolamide |
| 11 | Diacerein | 33 | Idelalisib | 55 | Laropiprant |
| 12 | Amoxapine | 34 | Tucidinostat | 56 | Vilazodone |
| 13 | Iodipamide | 35 | Vesnarinone | 57 | Ripasudil |
| 14 | Citric acid | 36 | Nilutamide | 58 | Cloxacillin (sodium monohydrate) |
| 15 | Butamben | 37 | Estropipate | 59 | Clofarabine |
| 16 | Anethole (trithione) | 38 | Micafungin (sodium) | 60 | Olanzapine |
| 17 | Diatrizoic acid | 39 | LCZ696 | 61 | Ivacaftor |
| 18 | Trifluridine | 40 | Camphor | 62 | Diclofenac (Sodium) |
| 19 | Thioridazine (hydrochloride) | 41 | Clorprenaline hydrochloride | 63 | Dihydroergotoxine (mesylate) |
| 20 | Benzthiazide | 42 | D-Panthenol | 64 | Gestodene |
| 21 | 5-Aminolevulinic acid (hydrochloride) | 43 | Mestranol | - | - |

PLATE 21

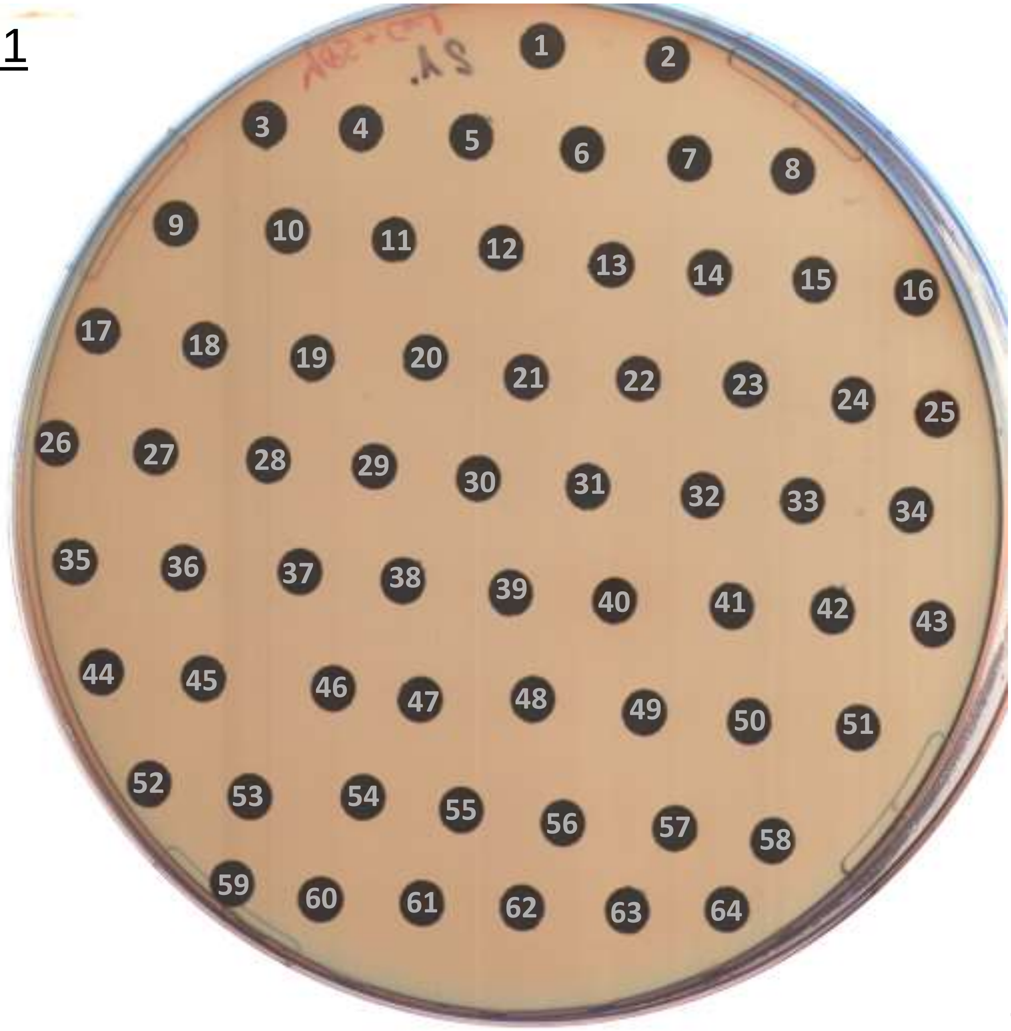

24 h

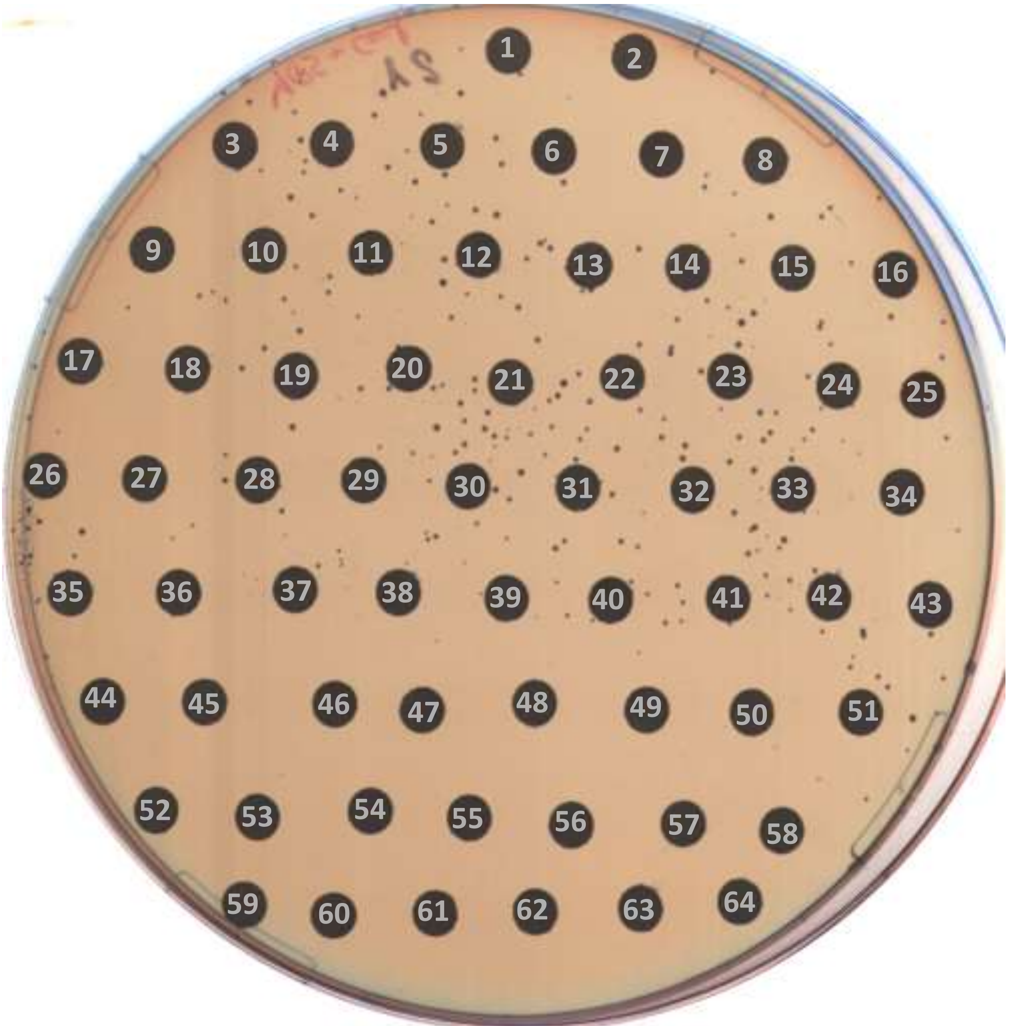

48 h

### PLATE 21

| no. | compound | no. | compound | no. | compound |
|---|---|---|---|---|---|
| - | - | 22 | Furosemide | 44 | Bedaquiline (fumarate) |
| 1 | Laquinimod | 23 | Tofogliflozin (hydrate) | 45 | Pranoprofen |
| 2 | Cilnidipine | 24 | Atrasentan (hydrochloride) | 46 | Acebutolol (hydrochloride) |
| 3 | Terazosin (hydrochloride dihydrate) | 25 | Opicapone | 47 | Tegaserod (maleate) |
| 4 | Rosiglitazone | 26 | Ciprofibrate | 48 | Prazosin (hydrochloride) |
| 5 | Proparacaine (Hydrochloride) | 27 | Famotidine | 49 | Salmeterol (xinafoate) |
| 6 | Dasabuvir | 28 | (±)-Bisoprolol (hemifumarate) | 50 | Seratrodast |
| 7 | Ledipasvir (D-tartrate) | 29 | Brexpiprazole | 51 | Atracurium (besylate) |
| 8 | Lopinavir | 30 | Milnacipran (hydrochloride) | 52 | Cytarabine |
| 9 | Rupatadine (Fumarate) | 31 | Benazepril (hydrochloride) | 53 | Nedocromil |
| 10 | Linagliptin | 32 | Sodium Picosulfate | 54 | Venlafaxine (hydrochloride) |
| 11 | Famciclovir | 33 | Abacavir | 55 | Tolfenamic Acid |
| 12 | Tipiracil (hydrochloride) | 34 | Terbinafine | 56 | Dasatinib |
| 13 | Selexipag | 35 | Betamethasone | 57 | Ixazomib citrate |
| 14 | Vecuronium (bromide) | 36 | Azelaic acid | 58 | Tetrabenazine |
| 15 | Doxifluridine | 37 | Vidarabine | 59 | Miconazole (nitrate) |
| 16 | Benserazide (hydrochloride) | 38 | Zafirlukast | 60 | Amorolfine (hydrochloride) |
| 17 | Vardenafil (hydrochloride) | 39 | Sacubitril | 61 | Solifenacin (Succinate) |
| 18 | Moclobemide | 40 | 5-Aminosalicylic Acid | 62 | L-Ascorbic acid |
| 19 | Procarbazine (Hydrochloride) | 41 | Methocarbamol | 63 | Pefloxacin (mesylate) |
| 20 | Floxuridine | 42 | Fenofibric acid | 64 | Lomefloxacin (hydrochloride) |
| 21 | Trimethoprim | 43 | Articaine (hydrochloride) | - | - |

PLATE 22

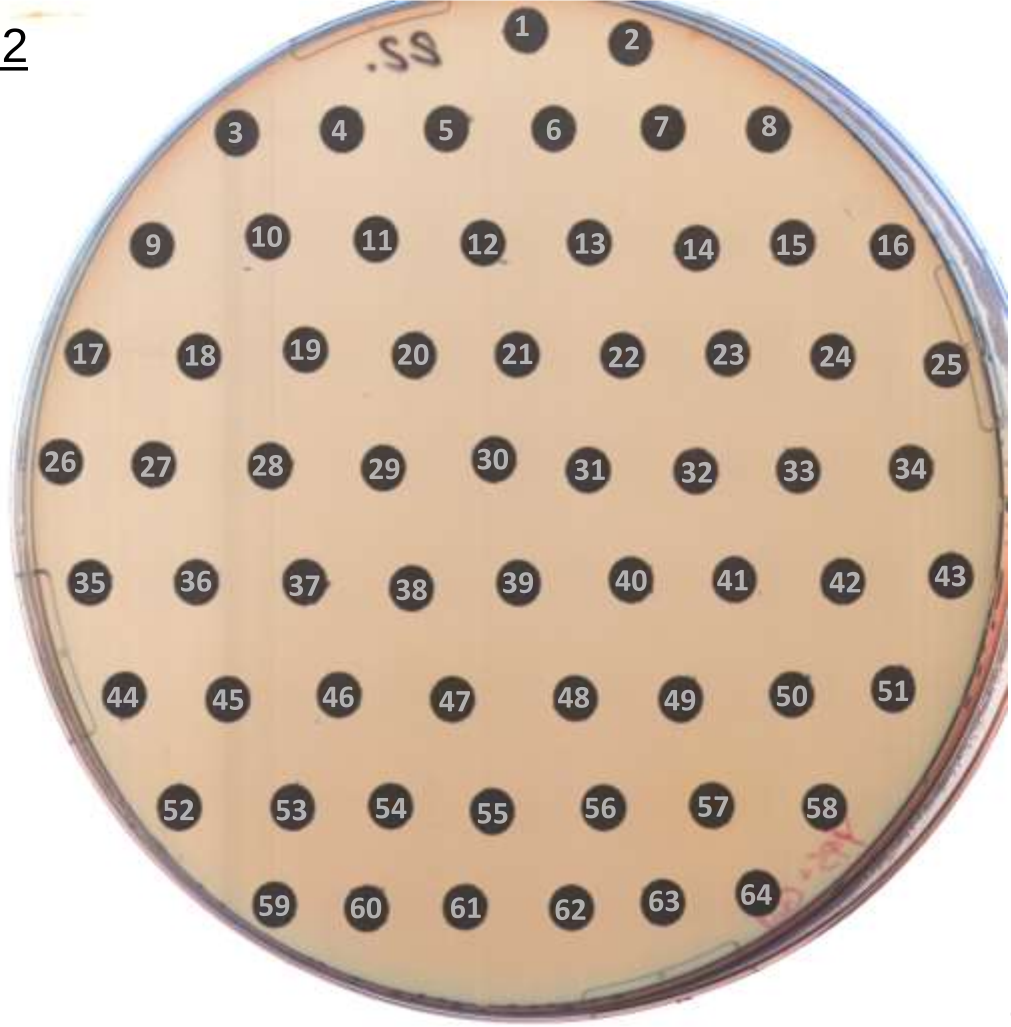

24 h

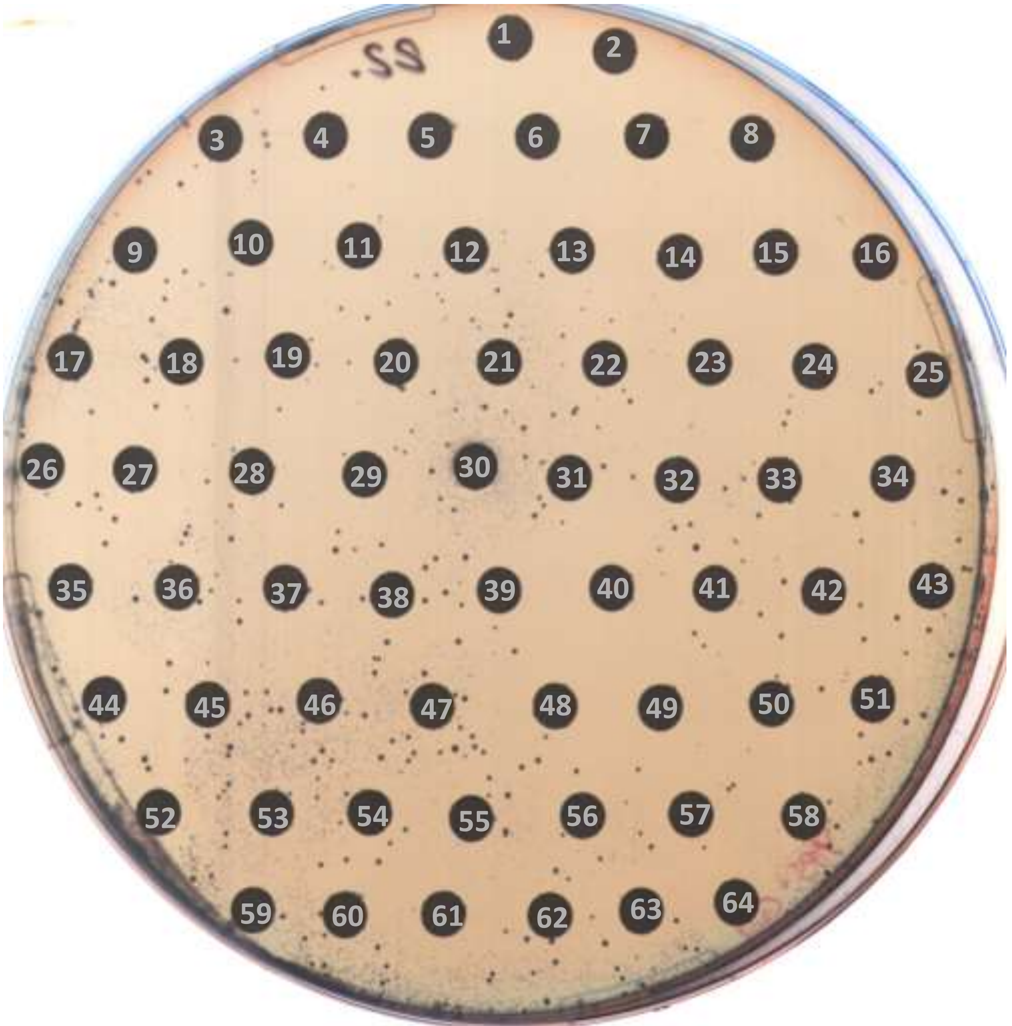

48 h

### PLATE 22

| no. | compound | no. | compound | no. | compound |
|---|---|---|---|---|---|
| - | - | 22 | Chlorprothixene | 44 | Acipimox |
| 1 | Anagrelide (hydrochloride) | 23 | Tobramycin | 45 | Cyclic somatostatin |
| 2 | Granisetron (Hydrochloride) | 24 | Cariprazine (hydrochloride) | 46 | Naproxen (sodium) |
| 3 | Cytisinicline | 25 | Sulfamonomethoxine | 47 | Chlortetracycline (hydrochloride) |
| 4 | Thiamine monochloride | 26 | Pantoprazole (sodium) | 48 | Vinpocetine |
| 5 | Mebhydrolin (napadisylate) | 27 | Ceftaroline fosamil | 49 | Clemastine (fumarate) |
| 6 | Norfloxacin | 28 | Niclosamide | 50 | Finasteride |
| 7 | Midostaurin | 29 | Thiamphenicol | 51 | Tolazoline (hydrochloride) |
| 8 | Fluconazole | 30 | Bedaquiline | 52 | Upadacitinib |
| 9 | Bilastine | 31 | Milrinone | 53 | Noscapine |
| 10 | Tasimelteon | 32 | Ouabain (Octahydrate) | 54 | Ranolazine |
| 11 | Dabigatran etexilate (mesylate) | 33 | Chlormethine (hydrochloride) | 55 | Dacarbazine |
| 12 | Diazoxide | 34 | Thalidomide | 56 | Penicillamine |
| 13 | Sofosbuvir | 35 | Atosiban | 57 | Erdafitinib |
| 14 | Troglitazone | 36 | D-Cycloserine | 58 | Pyrazinamide |
| 15 | Blonanserin | 37 | Larotrectinib sulfate | 59 | Flavoxate (hydrochloride) |
| 16 | Varenicline (Tartrate) | 38 | Paritaprevir | 60 | L-Epinephrine (Bitartrate) |
| 17 | Bromfenac (sodium hydrate) | 39 | Benznidazol | 61 | Hydroxyprogesterone caproate |
| 18 | Proglumide | 40 | Pimavanserin | 62 | Spiramycin |
| 19 | L-Carnitine (hydrochloride) | 41 | Pramiracetam | 63 | Glipizide |
| 20 | Estradiol (cypionate) | 42 | Crizotinib | 64 | Tafenoquine (Succinate) |
| 21 | Tedizolid | 43 | Sitafloxacin (hydrate) | - | - |

PLATE 23

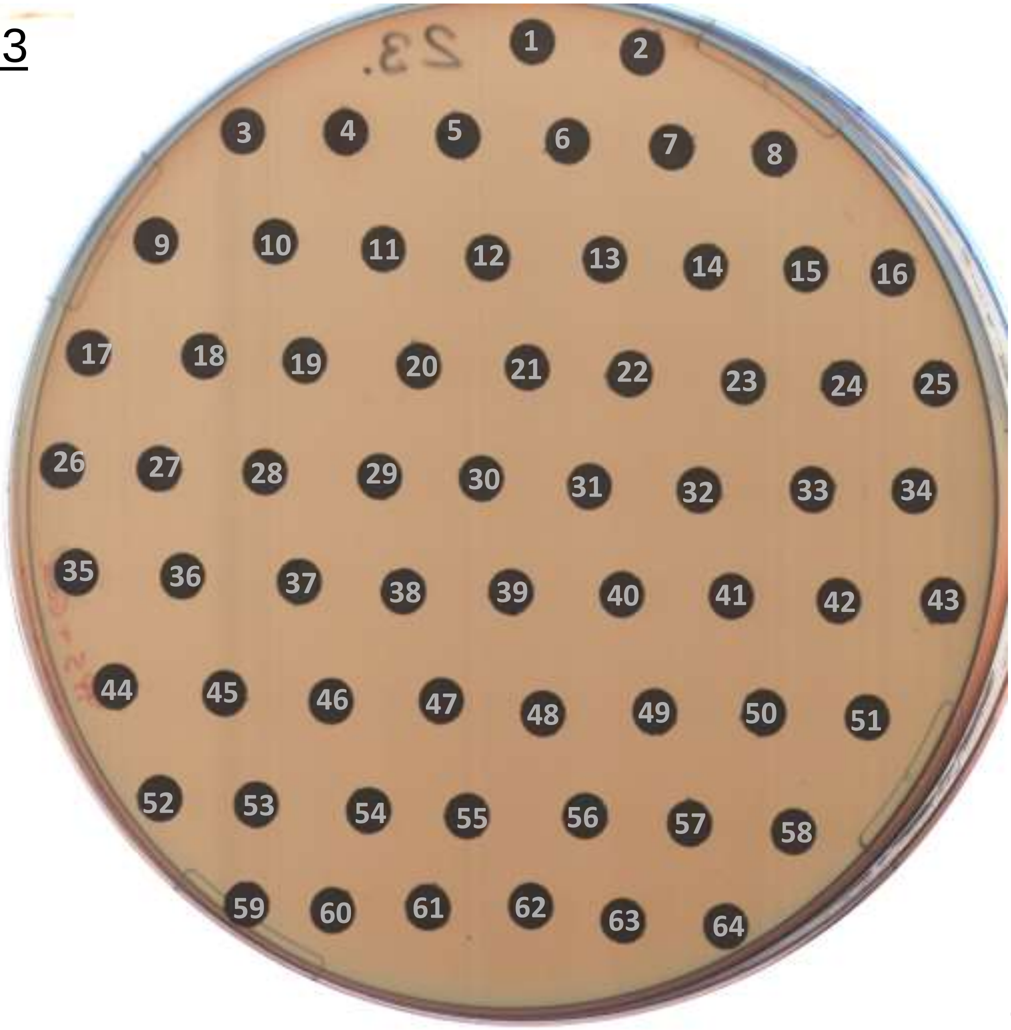

24 h

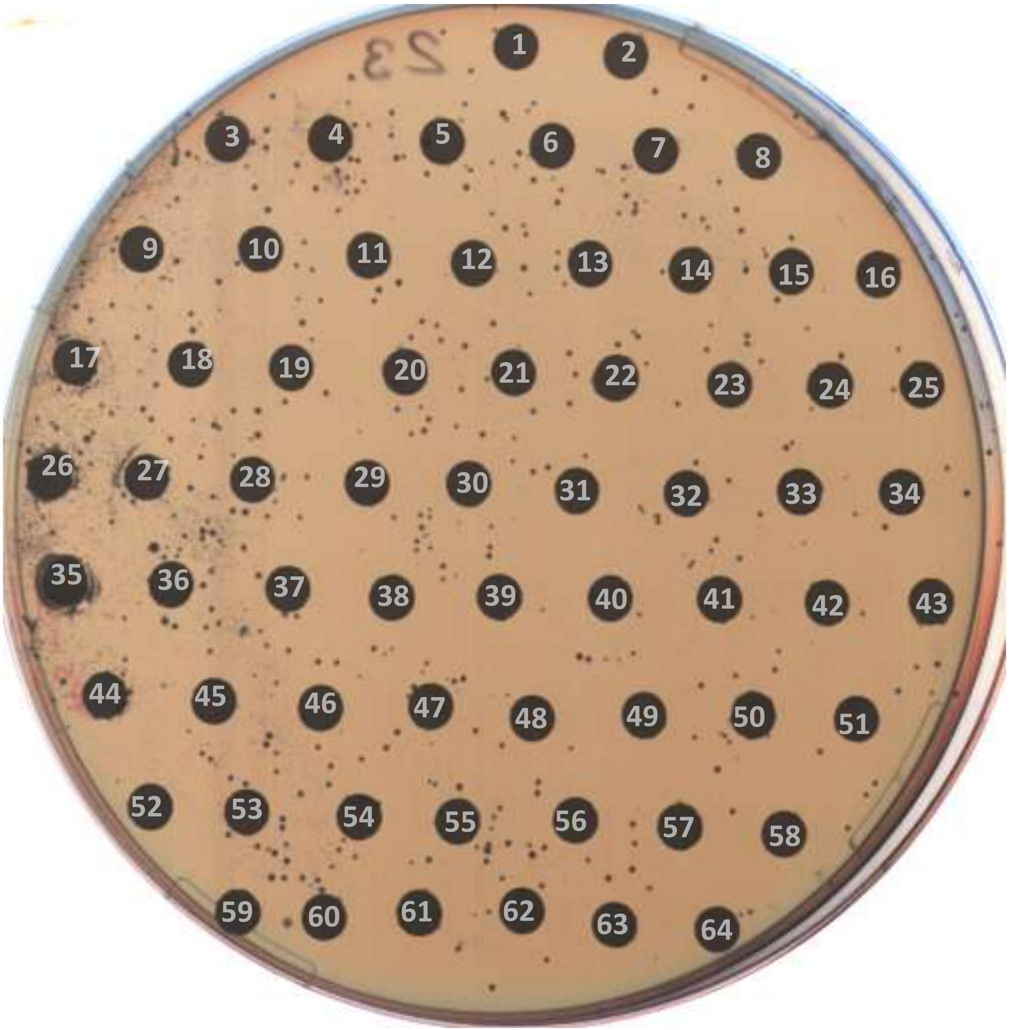

48 h

### PLATE 23

| no. | compound | no. | compound | no. | compound |
|---|---|---|---|---|---|
| - | - | 22 | Folic acid | 44 | Amezinium (methylsulfate) |
| 1 | Azasetron (hydrochloride) | 23 | Camostat (mesylate) | 45 | Vonoprazan |
| 2 | Aminophylline | 24 | Ornidazole | 46 | Sulpiride |
| 3 | Cefaclor | 25 | Naratriptan (hydrochloride) | 47 | Sulfaguanidine |
| 4 | Panobinostat | 26 | Flufenamic acid | 48 | Dibucaine |
| 5 | Mometasone furoate | 27 | Bronopol | 49 | Alimemazine hemitartrate |
| 6 | Mirtazapine | 28 | Milnacipran ((1S-cis) hydrochloride) | 50 | Propyphenazone |
| 7 | Torsemide | 29 | Ethynyl Estradiol | 51 | Nilotinib (monohydrochloride monohydrate) |
| 8 | Dimesna | 30 | Mianserin (hydrochloride) | 52 | Iohexol |
| 9 | Moxisylyte (hydrochloride) | 31 | Ipratropium (bromide) | 53 | Tosufloxacin (tosylate hydrate) |
| 10 | Ethamsylate | 32 | Phenindione | 54 | D-Sorbitol |
| 11 | Gabapentin (hydrochloride) | 33 | Chlormezanone | 55 | Phentolamine (mesylate) |
| 12 | Natamycin | 34 | Flubendazole | 56 | Benzydamine (hydrochloride) |
| 13 | Galanthamine | 35 | Sulfamethazine | 57 | Phenytoin |
| 14 | Niacin | 36 | Meglumine | 58 | Valproic acid (sodium salt) |
| 15 | Oseltamivir (phosphate) | 37 | Roxithromycin | 59 | DL-Menthol |
| 16 | Ivosidenib | 38 | Maprotiline (hydrochloride) | 60 | Atenolol |
| 17 | 6-Acetamidohexanoic acid | 39 | Homatropine (Bromide) | 61 | Diosmin |
| 18 | Lesinurad | 40 | Niflumic acid | 62 | Riociguat |
| 19 | Uridine triacetate | 41 | Danthron | 63 | Selumetinib |
| 20 | Bicyclol | 42 | Oxaceprol | 64 | Nafcillin (sodium monohydrate) |
| 21 | (-)-Huperzine A | 43 | Suplatast (Tosilate) | - | - |

PLATE 24

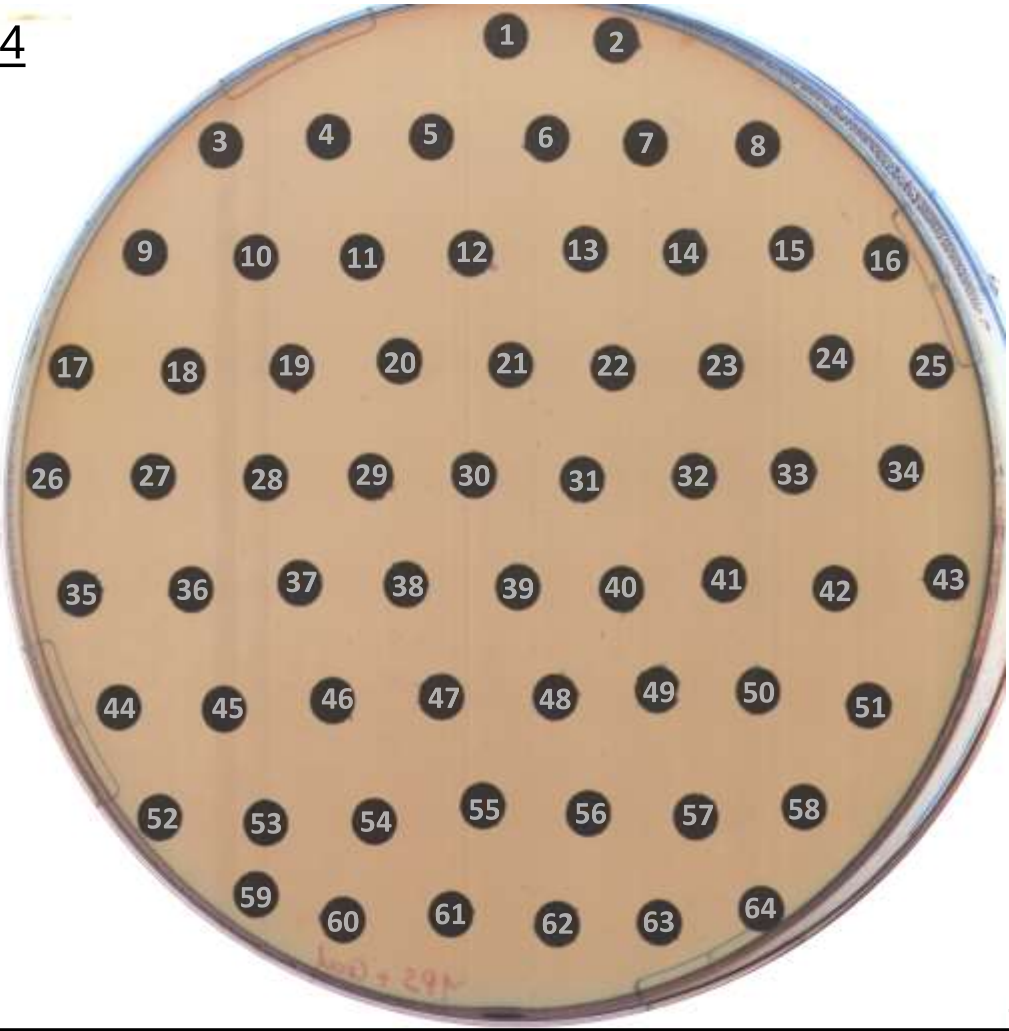

24 h

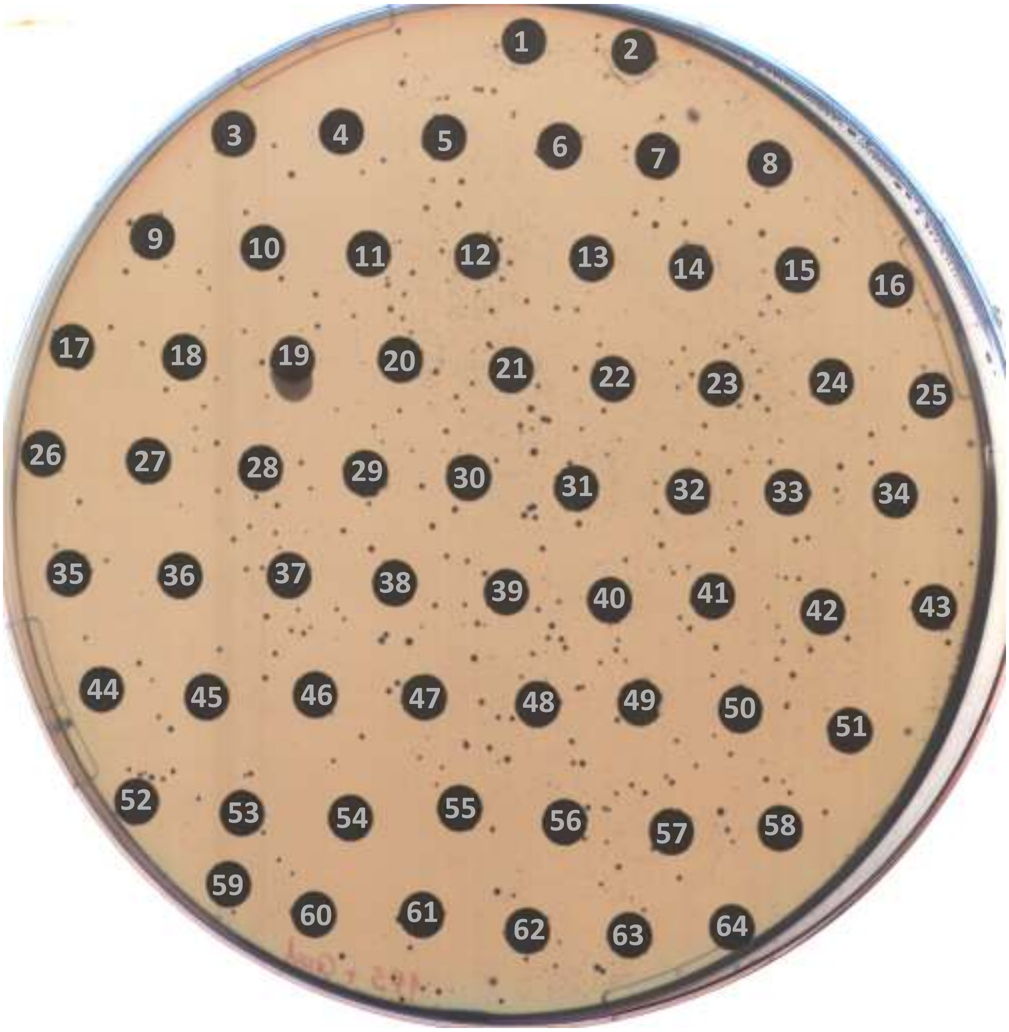

48 h

### PLATE 24

| no. | compound | no. | compound | no. | compound |
|---|---|---|---|---|---|
| - | - | 22 | Lusutrombopag | 44 | Fulvestrant |
| 1 | Dextrose | 23 | Propranolol (hydrochloride) | 45 | Telmisartan |
| 2 | Valpromide | 24 | Delamanid | 46 | Hydroxocobalamin (monohydrochloride) |
| 3 | Orphenadrine (citrate) | 25 | Pranlukast (hemihydrate) | 47 | Isoprenaline (hydrochloride) |
| 4 | Dapiprazole (hydrochloride) | 26 | Ketanserin (tartrate) | 48 | Cepharanthine |
| 5 | Moroxydine (hydrochloride) | 27 | Raloxifene (hydrochloride) | 49 | Novobiocin (Sodium) |
| 6 | Benzocaine | 28 | Nilotinib | 50 | Ifenprodil (tartrate) |
| 7 | Idebenone | 29 | Malotilate | 51 | HDAC-IN-7 |
| 8 | Sulfadimethoxine | 30 | Sulfadiazine | 52 | Nabumetone |
| 9 | Dexchlorpheniramine (maleate) | 31 | Ibutilide (fumarate) | 53 | Pimethixene maleate |
| 10 | Moxonidine | 32 | Pyrantel (tartrate) | 54 | Fingolimod (hydrochloride) |
| 11 | Quinine (hydrochloride dihydrate) | 33 | Diethylstilbestrol | 55 | Lomustine |
| 12 | Cyclosporin A | 34 | Methimazole | 56 | L-Proline |
| 13 | Olmesartan | 35 | Gliclazide | 57 | Baloxavir marboxil |
| 14 | Tolbutamide | 36 | Flunarizine (dihydrochloride) | 58 | Cinobufotalin |
| 15 | Dapsone | 37 | Ticlopidine (hydrochloride) | 59 | Coumarin |
| 16 | Escin | 38 | Gestrinone | 60 | Ecabet (sodium) |
| 17 | Sunitinib | 39 | Peretinoin | 61 | Alectinib (Hydrochloride) |
| 18 | Piroctone olamine | 40 | Thio-TEPA | 62 | Prostaglandin E2 |
| 19 | Thiamine nitrate | 41 | Megestrol | 63 | Sulfacetamide |
| 20 | Sulfogaiacol | 42 | Peficitinib | 64 | Cephapirin (sodium) |
| 21 | Salicylamide | 43 | Biotin | - | - |

### PLATE 25

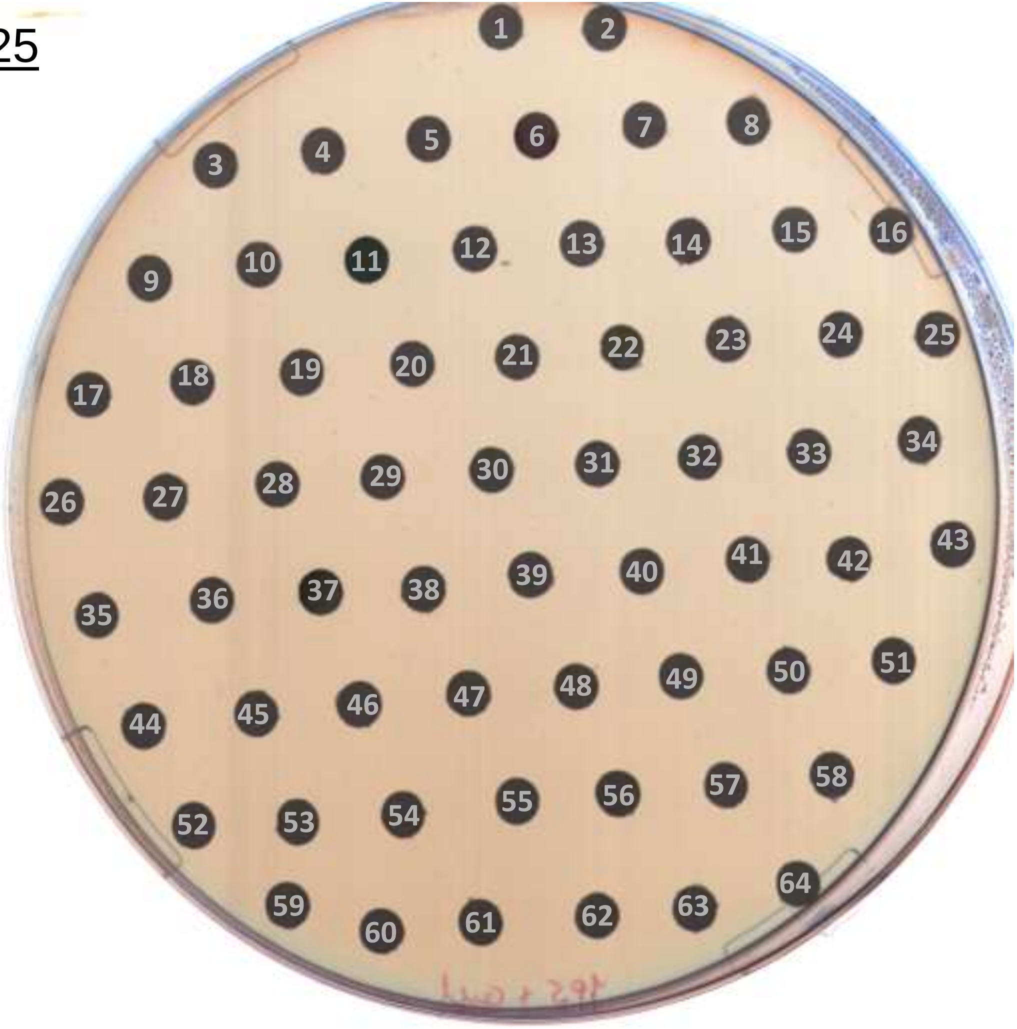

24 h

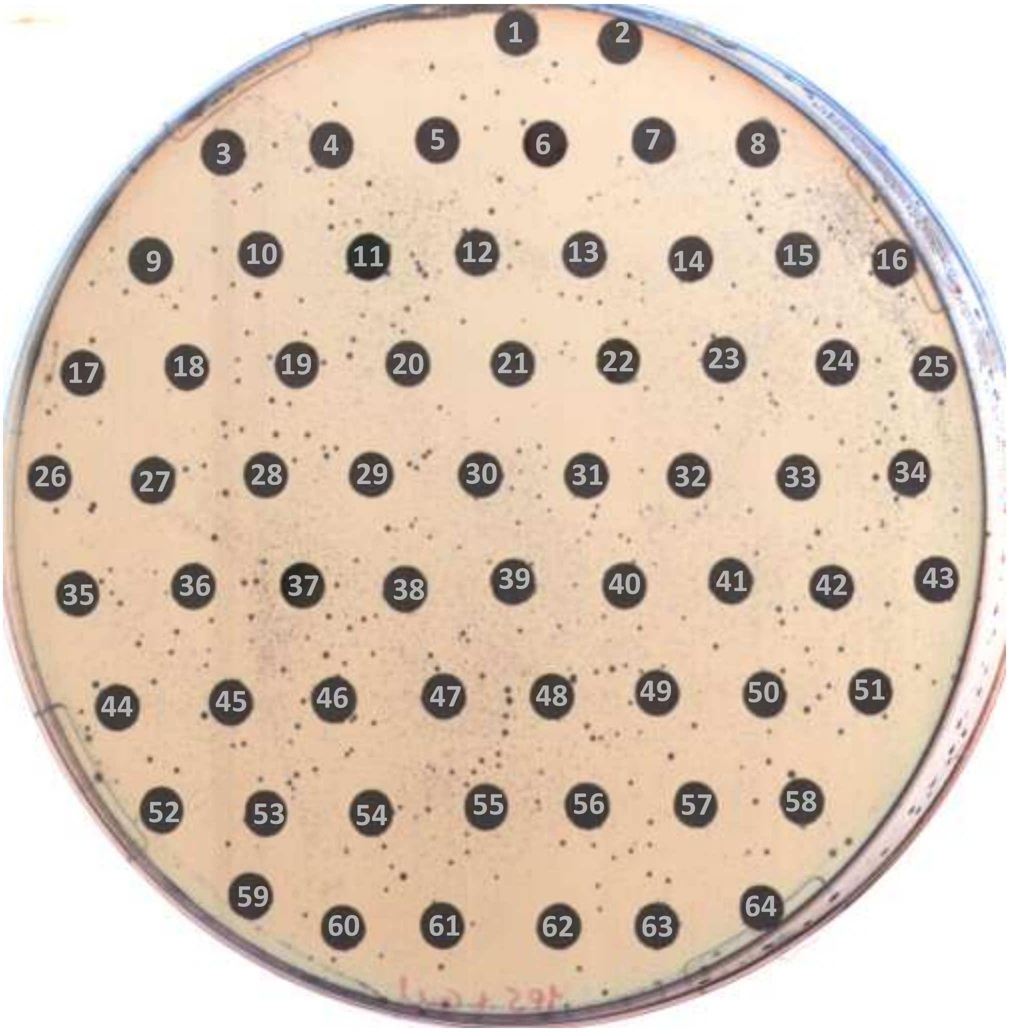

48 h

### PLATE 25

| no. | compound | no. | compound | no. | compound |
|---|---|---|---|---|---|
| - | - | 22 | Venetoclax | 44 | 10-Undecenoic acid (zinc salt) |
| 1 | Terbutaline (sulfate) | 23 | Cefotaxime (sodium salt) | 45 | Dexamethasone phosphate disodium |
| 2 | Limaprost | 24 | Niraparib | 46 | Gatifloxacin (hydrochloride) |
| 3 | Roxadustat | 25 | Uracil | 47 | L-Ascorbic acid sodium salt |
| 4 | Tolcapone | 26 | Gefarnate | 48 | Hydralazine (hydrochloride) |
| 5 | Indacaterol | 27 | Efloxate | 49 | Aceglutamide |
| 6 | Pirarubicin (Hydrochloride) | 28 | Indacaterol (maleate) | 50 | Sacubitril hemicalcium salt |
| 7 | Irinotecan (hydrochloride) | 29 | Donepezil | 51 | Clinofibrate |
| 8 | Taltirelin (acetate) | 30 | Acyclovir | 52 | Cephradine |
| 9 | Sildenafil | 31 | Tacrolimus (monohydrate) | 53 | Penfluridol |
| 10 | Chenodeoxycholic Acid | 32 | Eslicarbazepine | 54 | Penbutolol (sulfate) |
| 11 | Sodium copper chlorophyllin A | 33 | Fingolimod | 55 | Cephalexin (monohydrate) |
| 12 | Pretomanid | 34 | Sitagliptin | 56 | Dolutegravir (sodium) |
| 13 | Istradefylline | 35 | Desoximetasone | 57 | Bendazac L-Lysine |
| 14 | Bromocriptine (mesylate) | 36 | Liothyronine (sodium) | 58 | Vincamine |
| 15 | Sulfaphenazole | 37 | Verteporfin | 59 | Oxeladin (citrate) |
| 16 | Fidarestat | 38 | Dexamethasone | 60 | Rocuronium (Bromide) |
| 17 | (+)-Kavain | 39 | Flutamide | 61 | Candesartan |
| 18 | Quetiapine (hemifumarate) | 40 | Vincristine (sulfate) | 62 | Ziprasidone |
| 19 | Vandetanib | 41 | Flunisolide | 63 | Trihexyphenidyl (hydrochloride) |
| 20 | L-Leucine | 42 | Colchicine | 64 | Fondaparinux (sodium) |
| 21 | Nomegestrol acetate | 43 | Amiloride hydrochloride dihydrate | - | - |

PLATE 26

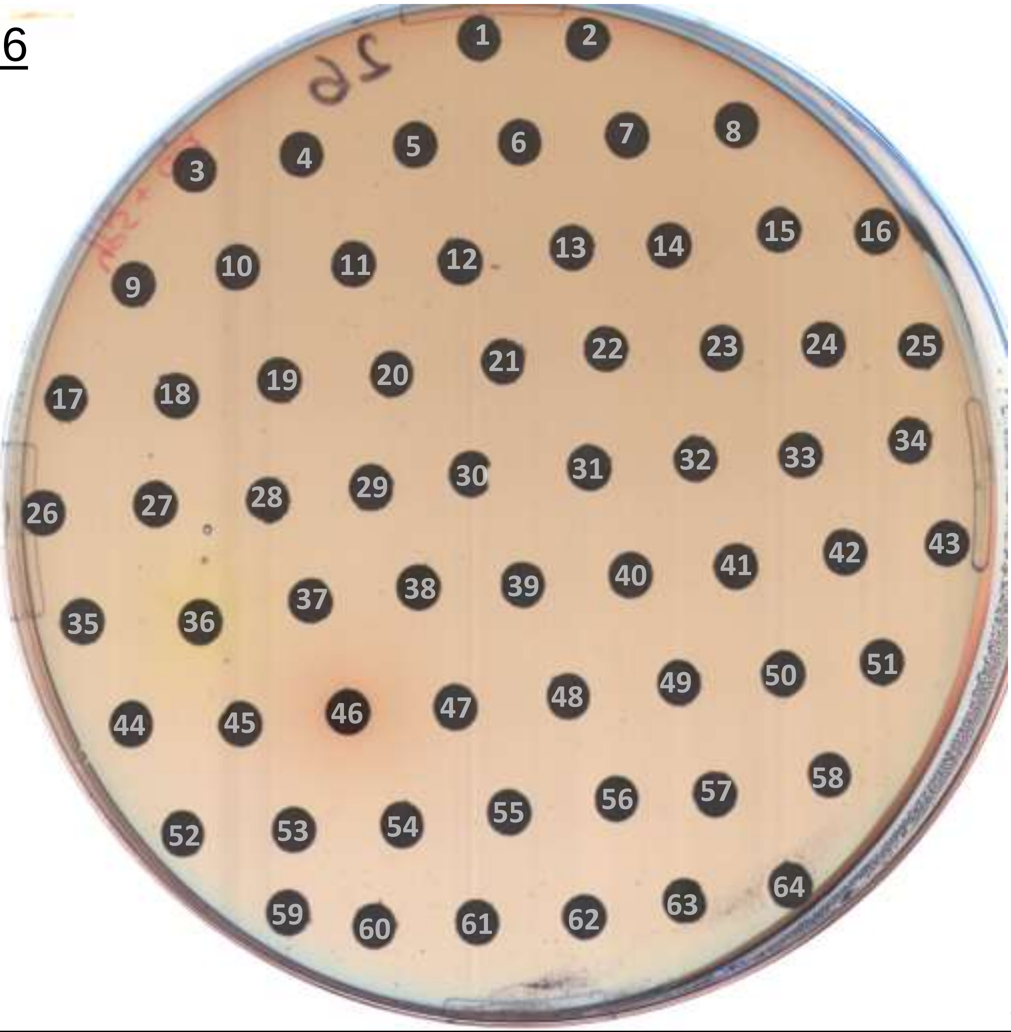

24 h

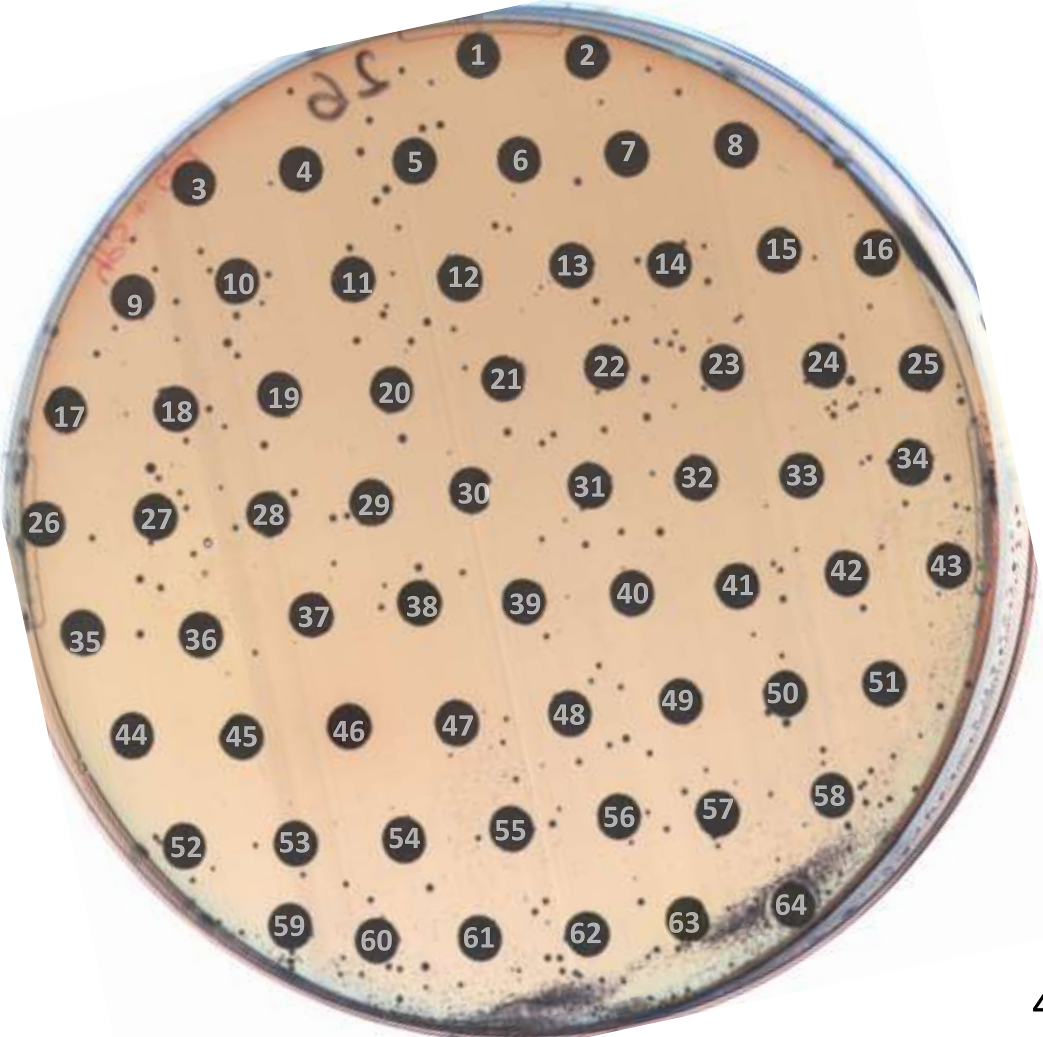

48 h

### PLATE 26

| no. | compound | no. | compound | no. | compound |
|---|---|---|---|---|---|
| - | - | 22 | Glycerol | 44 | Mequitazine |
| 1 | Imiquimod (hydrochloride) | 23 | Agomelatine (hydrochloride) | 45 | Ketorolac (tromethamine salt) |
| 2 | Imiquimod | 24 | Kasugamycin (hydrochloride hydrate) | 46 | Idarubicin (hydrochloride) |
| 3 | Gadoxetate (Disodium) | 25 | Proglumide (sodium) | 47 | Mepyramine maleate |
| 4 | Betamipron | 26 | α-Lipoic Acid | 48 | Liothyronine |
| 5 | Sulfalene | 27 | Alectinib | 49 | Norethindrone |
| 6 | Idramantone | 28 | Fumaric acid | 50 | Leuprolide (Acetate) |
| 7 | Rasagiline (mesylate) | 29 | Mebendazole | 51 | Acetylcholine (chloride) |
| 8 | Cephalexin | 30 | Molsidomine | 52 | Methyl Salicylate |
| 9 | sn-Glycero-3-phosphocholine | 31 | Pipemidic acid | 53 | Celecoxib |
| 10 | Tenofovir (Disoproxil Fumarate) | 32 | Fosphenytoin (disodium) | 54 | Clindamycin (phosphate) |
| 11 | Choline Fenofibrate | 33 | Ticagrelor | 55 | Vigabatrin (hydrochloride) |
| 12 | Palonosetron (Hydrochloride) | 34 | Cilazapril (monohydrate) | 56 | Pericyazine |
| 13 | Glucosamine (hydrochloride) | 35 | Trimebutine (maleate) | 57 | Clidinium (bromide) |
| 14 | Flopropione | 36 | Rifaximin | 58 | Gemcitabine (hydrochloride) |
| 15 | Acrivastine | 37 | Meclofenamate (sodium) | 59 | Felypressin |
| 16 | Octenidine (dihydrochloride) | 38 | Bazedoxifene (acetate) | 60 | Metipranolol hydrochloride |
| 17 | Neratinib | 39 | Digitoxin | 61 | Serotonin hydrochloride |
| 18 | Gefitinib (hydrochloride) | 40 | Fluocinolone (Acetonide) | 62 | Canrenone |
| 19 | Valsartan | 41 | Vernakalant (Hydrochloride) | 63 | Nintedanib |
| 20 | Ofloxacin | 42 | Ixabepilone | 64 | Pomalidomide |
| 21 | Risperidone | 43 | Lipoic acid | - | - |

PLATE 27

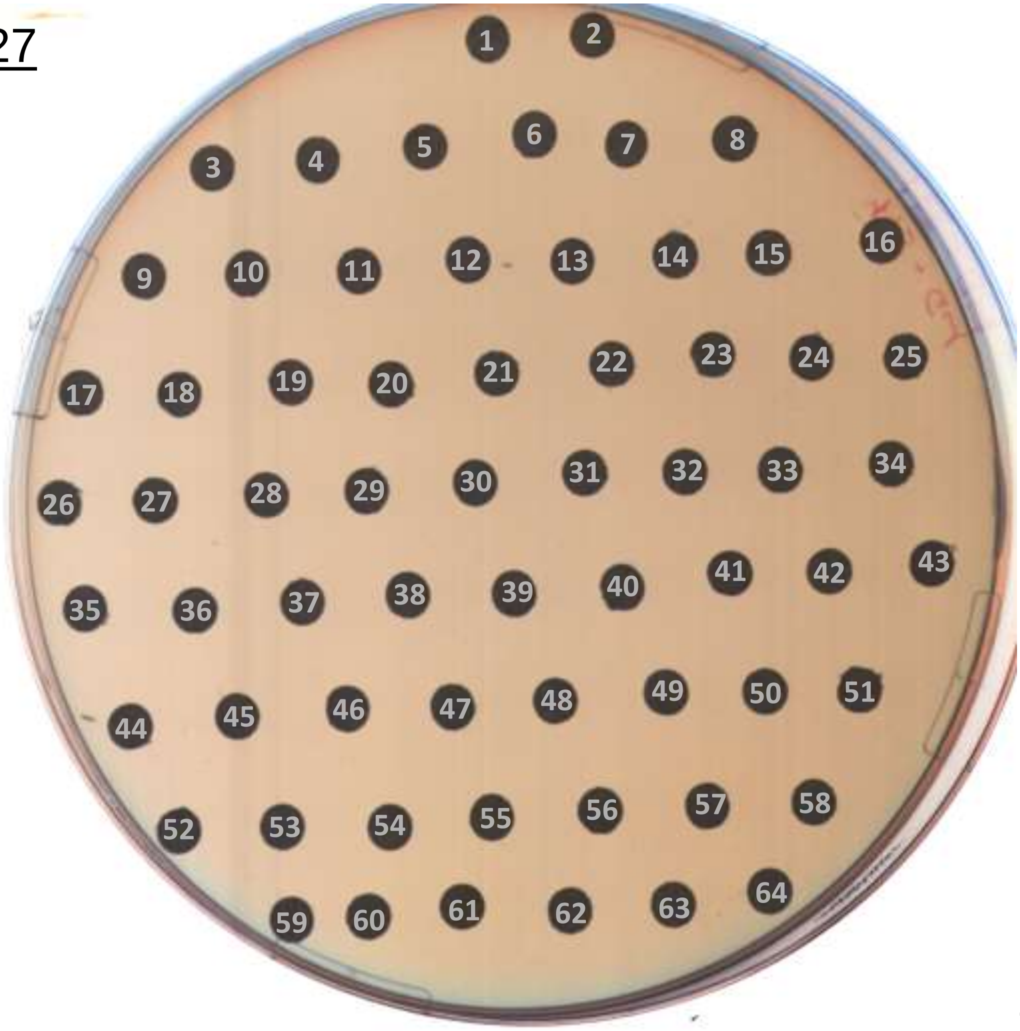

24 h

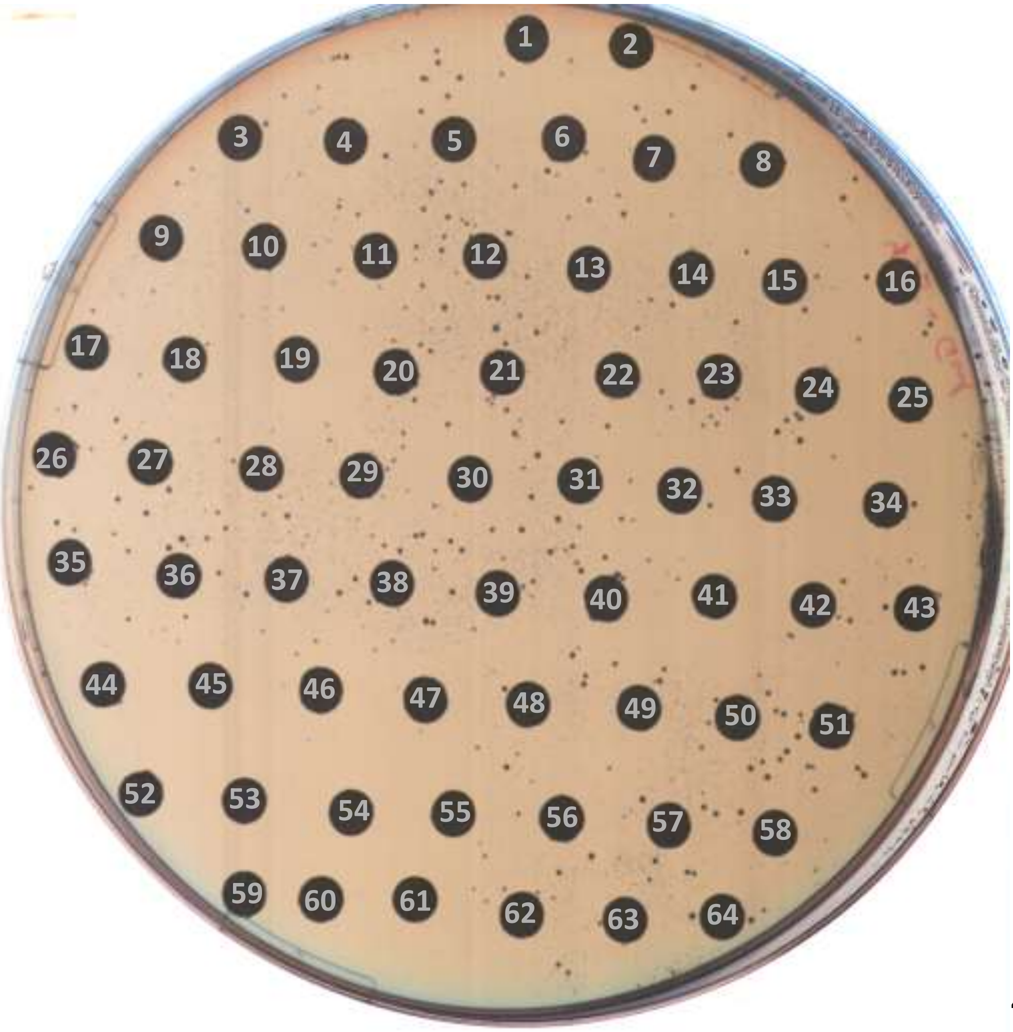

48 h

### PLATE 27

| no. | compound | no. | compound | no. | compound |
|---|---|---|---|---|---|
| - | - | 22 | Lomerizine dihydrochloride | 44 | Lapatinib |
| 1 | Pemetrexed | 23 | Almitrine mesylate | 45 | Fostamatinib Disodium |
| 2 | Lapatinib (ditosylate) | 24 | Tafluprost | 46 | Ribociclib succinate |
| 3 | Regadenoson | 25 | Tranylcypromine (hemisulfate) | 47 | Deprodone propionate |
| 4 | Furosemide (sodium) | 26 | Cobimetinib | 48 | Octreotide (acetate) |
| 5 | Bictegravir | 27 | Bezafibrate | 49 | Dacomitinib |
| 6 | Docusate (Sodium) | 28 | Amoxicillin | 50 | Docetaxel (Trihydrate) |
| 7 | Phenolphthalein | 29 | Rauwolscine (hydrochloride) | 51 | Oxethazaine |
| 8 | Cefathiamidine | 30 | Latanoprost | 52 | Terbinafine hydrochloride |
| 9 | Phillyrin | 31 | Pantoprazole (sodium hydrate) | 53 | Neostigmine (methyl sulfate) |
| 10 | Carcainium (chloride) | 32 | Delapril (hydrochloride) | 54 | Nifurtimox |
| 11 | Nafamostat (mesylate) | 33 | Perphenazine | 55 | Dantrolene (sodium hemiheptahydrate) |
| 12 | Mebhydrolin | 34 | Empagliflozin | 56 | Primaquine (Diphosphate) |
| 13 | Imatinib | 35 | Monocrotaline | 57 | Fluorescein |
| 14 | Protriptyline (hydrochloride) | 36 | Melatonin | 58 | Paclitaxel |
| 15 | Eltrombopag | 37 | Ropinirole (hydrochloride) | 59 | Rapamycin |
| 16 | Docetaxel | 38 | Doxycycline (hyclate) | 60 | Glucosamine |
| 17 | Abiraterone | 39 | Santonin | 61 | Levosimendan |
| 18 | Dimercaprol | 40 | Dasatinib (hydrochloride) | 62 | Carboxin |
| 19 | Artesunate | 41 | Metixene hydrochloride hydrate | 63 | Belinostat |
| 20 | Minocycline (hydrochloride) | 42 | Sodium Salicylate | 64 | Sulindac |
| 21 | Atropine methyl bromide | 43 | Aprotinin | - | - |

PLATE 28

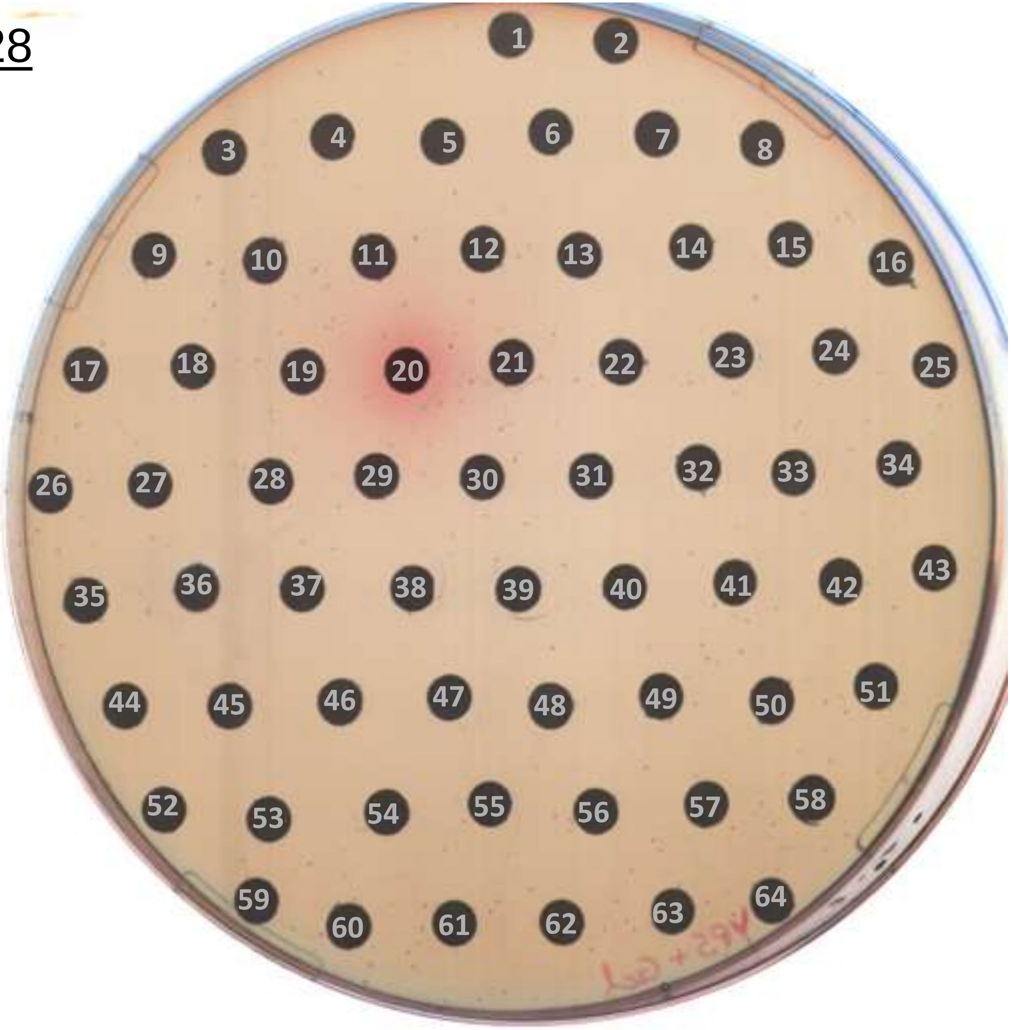

24 h

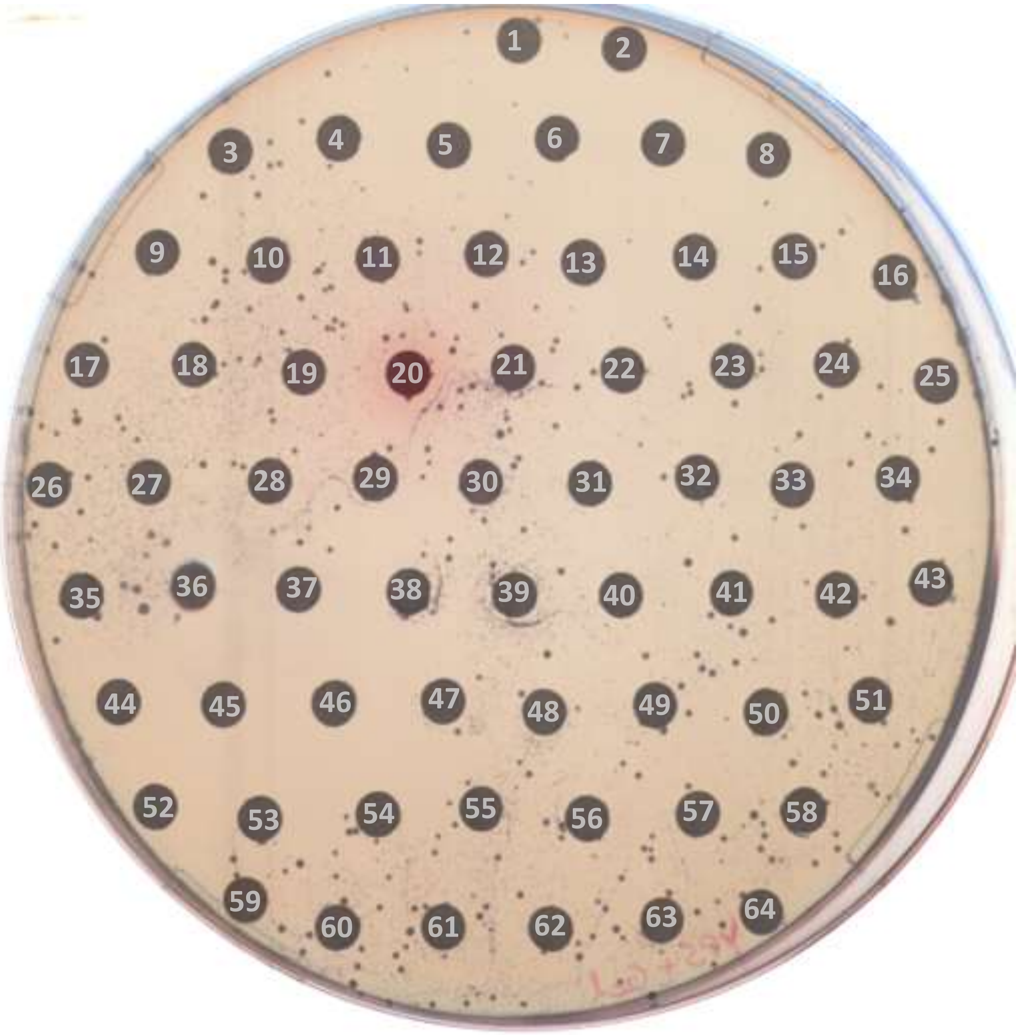

48 h

### PLATE 28

| no. | compound | no. | compound | no. | compound |
|---|---|---|---|---|---|
| - | - | 22 | L-Histidine | 44 | Amifostine |
| 1 | Cefazolin (sodium) | 23 | 4-(Aminomethyl)benzoic acid | 45 | Miltefosine |
| 2 | Catechin | 24 | Paromomycin (sulfate) | 46 | Alendronate (sodium hydrate) |
| 3 | Beclometasone | 25 | Gadobutrol | 47 | Zoledronic acid (monohydrate) |
| 4 | Hesperidin | 26 | Capreomycin (sulfate) | 48 | L-Ornithine |
| 5 | Dihydroartemisinin | 27 | L-Lysine hydrochloride | 49 | Gluconate (sodium) |
| 6 | Dolasetron | 28 | Guanethidine (sulfate) | 50 | DL-Arginine |
| 7 | Artemotil | 29 | Streptomycin (sulfate) | 51 | Deoxycholic acid sodium salt |
| 8 | Osimertinib mesylate | 30 | Palbociclib (hydrochloride) | 52 | Risedronate (sodium) |
| 9 | Citicoline (sodium) | 31 | Colistin (sulfate) | 53 | Sodium citrate (dihydrate) |
| 10 | Citric acid (trilithium salt tetrahydrate) | 32 | Levoleucovorin (Calcium) | 54 | Pamidronate (disodium pentahydrate) |
| 11 | L-Arginine | 33 | Gluconate (Calcium) | 55 | Plerixafor |
| 12 | L-Glutamic acid monosodium salt | 34 | Tenofovir (hydrate) | 56 | Bekanamycin |
| 13 | Plerixafor (octahydrochloride) | 35 | DL-Glutamine | 57 | Prednisolone (disodium phosphate) |
| 14 | Biapenem | 36 | Citicoline | 58 | L-Valine |
| 15 | L-Asparagine | 37 | L-Serine | 59 | Gastrodenol |
| 16 | Lisinopril (dihydrate) | 38 | Chloroquine (phosphate) | 60 | Amikacin (disulfate) |
| 17 | Netilmicin (sulfate) | 39 | Sugammadex (sodium) | 61 | 6-Aminocaproic acid |
| 18 | Tranexamic acid | 40 | L-Isoleucine | 62 | Peramivir (trihydrate) |
| 19 | L-Glutamine | 41 | L-Lysine | 63 | (R)-Baclofen |
| 20 | Merbromin | 42 | Ribostamycin (sulfate) | 64 | Glycine |
| 21 | L-Cysteine | 43 | Histamine (phosphate) | - | - |

PLATE 29

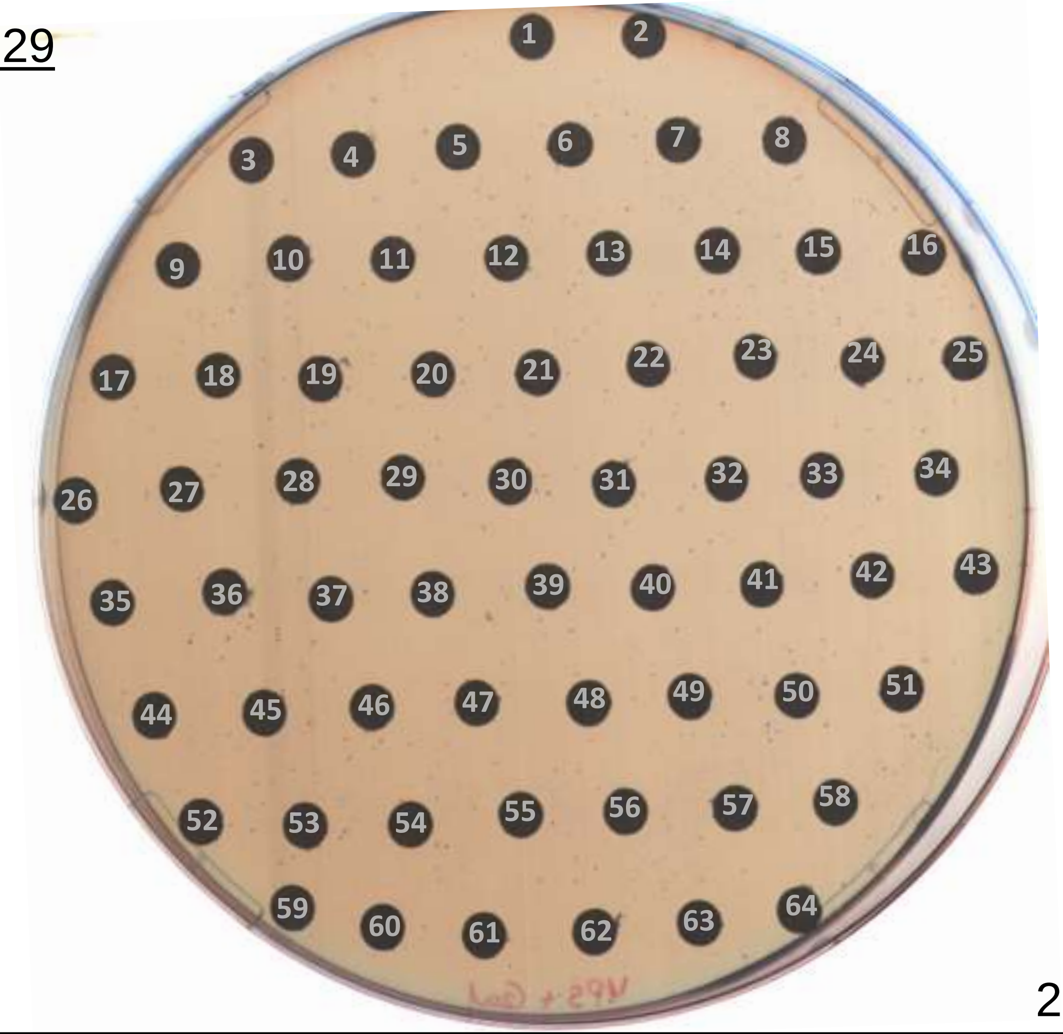

24 h

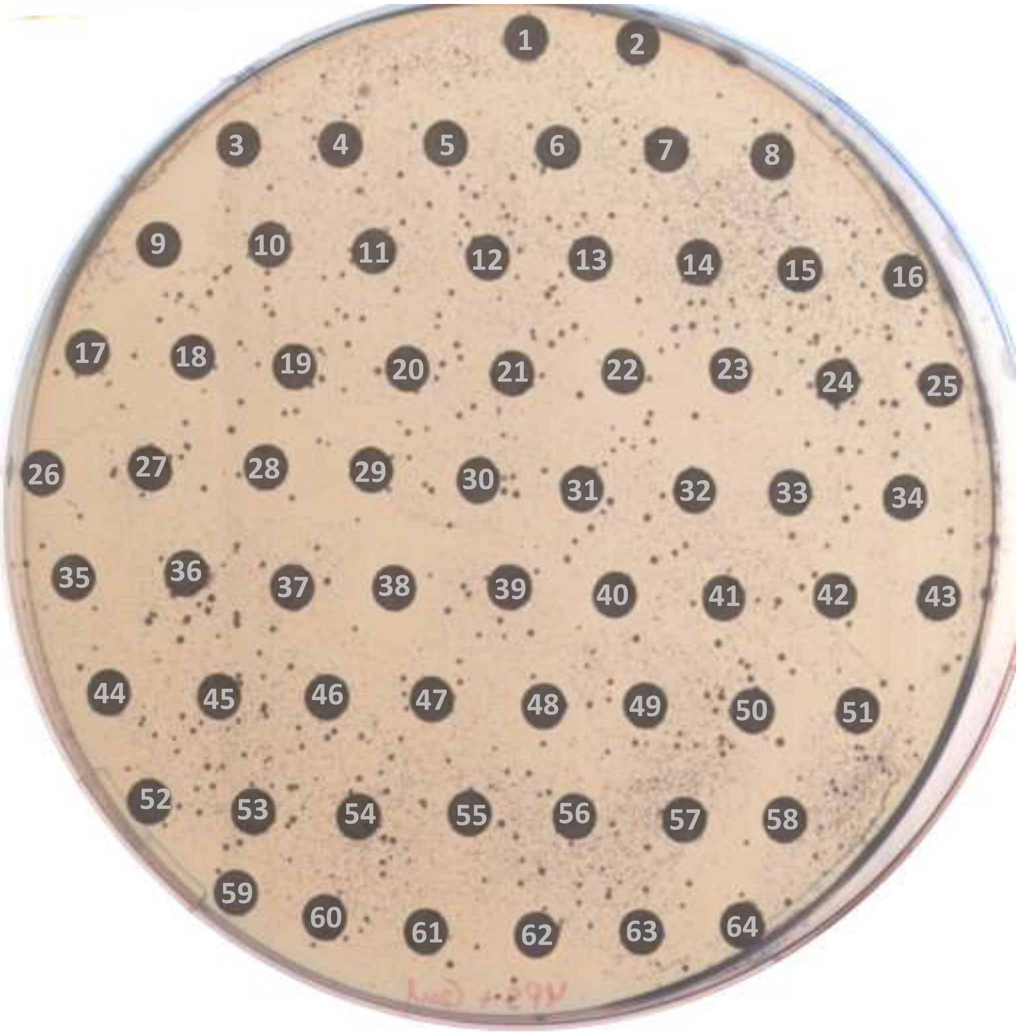

48 h

### PLATE 29

| no. | compound | no. | compound | no. | compound |
|---|---|---|---|---|---|
| - | - | 22 | Etofenamate | 44 | Flurandrenolide |
| 1 | Acamprosate (calcium) | 23 | Ruxolitinib | 45 | 6α-Methylprednisolone 21-hemisuccinate (sodium salt) |
| 2 | L-Arginine (hydrochloride) | 24 | Benzyl alcohol | 46 | Daidzin |
| 3 | Diquafosol (tetrasodium) | 25 | Boceprevir | 47 | Lobeline (hydrochloride) |
| 4 | Kanamycin (sulfate) | 26 | Vismodegib | 48 | Pregnenolone |
| 5 | Sisomicin (sulfate) | 27 | Tocofersolan | 49 | Cefetamet pivoxil (hydrochloride) |
| 6 | Hydroxychloroquine sulfate | 28 | Testosterone propionate | 50 | Tamsulosin |
| 7 | Neomycin (sulfate) | 29 | Almotriptan (malate) | 51 | Bupivacaine (hydrochloride) |
| 8 | Selenomethionine | 30 | Levobunolol (hydrochloride) | 52 | Oxymatrine |
| 9 | Mangafodipir (trisodium) | 31 | Trandolapril | 53 | Rifamycin (sodium) |
| 10 | Adefovir | 32 | Dicloxacillin (Sodium hydrate) | 54 | Phenacetin |
| 11 | Ethylenediaminetetraacetic acid (trisodium salt) | 33 | Minaprine (dihydrochloride) | 55 | Vitamin K4 |
| 12 | (R)-Serine | 34 | Histamine (dihydrochloride) | 56 | Levonorgestrel |
| 13 | Mildronate (dihydrate) | 35 | Ampicillin | 57 | (R)-(+)-Atenolol |
| 14 | L-Methionine | 36 | Dichlorphenamide | 58 | Vanillin |
| 15 | Mildronate | 37 | Sulfathiazole (sodium) | 59 | Quinestrol |
| 16 | Creatine | 38 | Inosine | 60 | Ampicillin (sodium) |
| 17 | (S)-Glutamic acid | 39 | Iloprost | 61 | Droperidol |
| 18 | Molindone (hydrochloride) | 40 | 17-Hydroxyprogesterone | 62 | Medroxyprogesterone |
| 19 | Topotecan (Hydrochloride) | 41 | Deslanoside | 63 | Acenocoumarol |
| 20 | Apixaban | 42 | Eptifibatide | 64 | Sultiame |
| 21 | Imidapril (hydrochloride) | 43 | Daidzein | - | - |

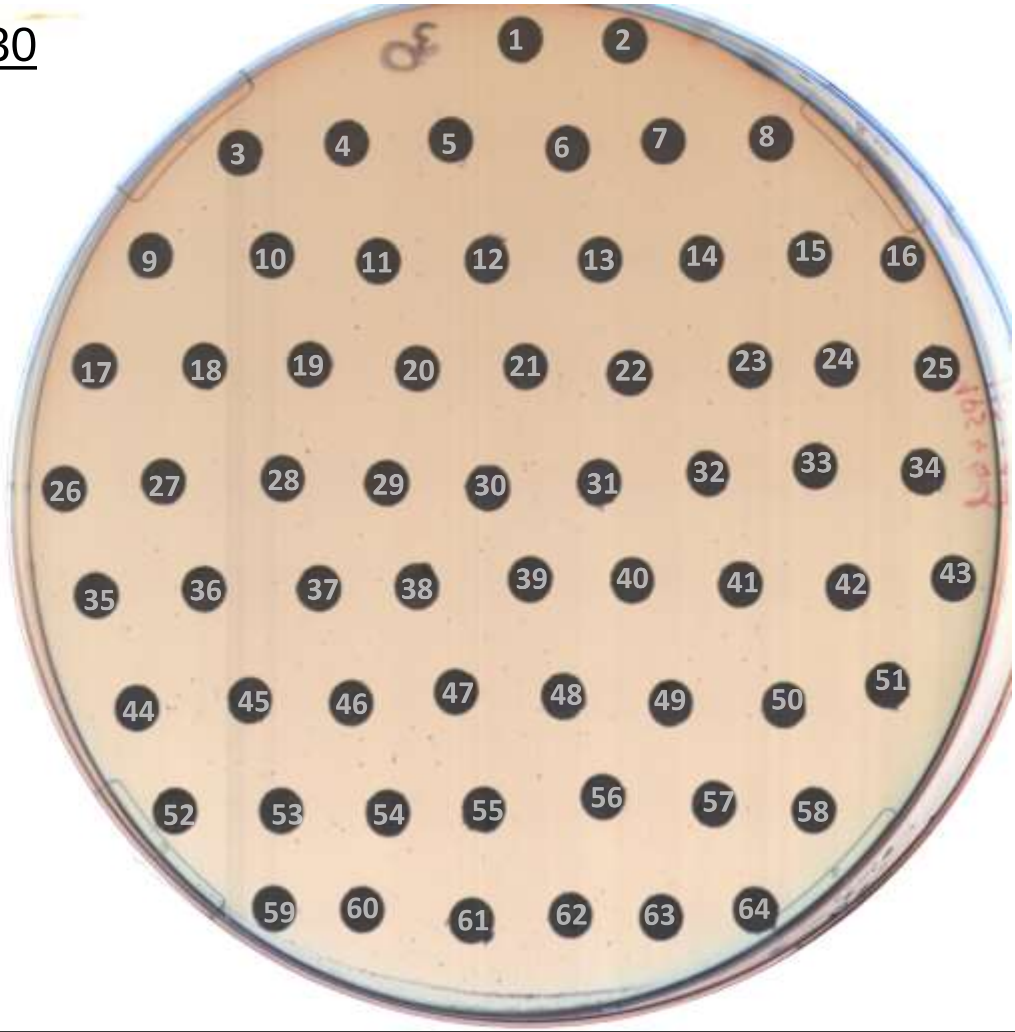

24 h

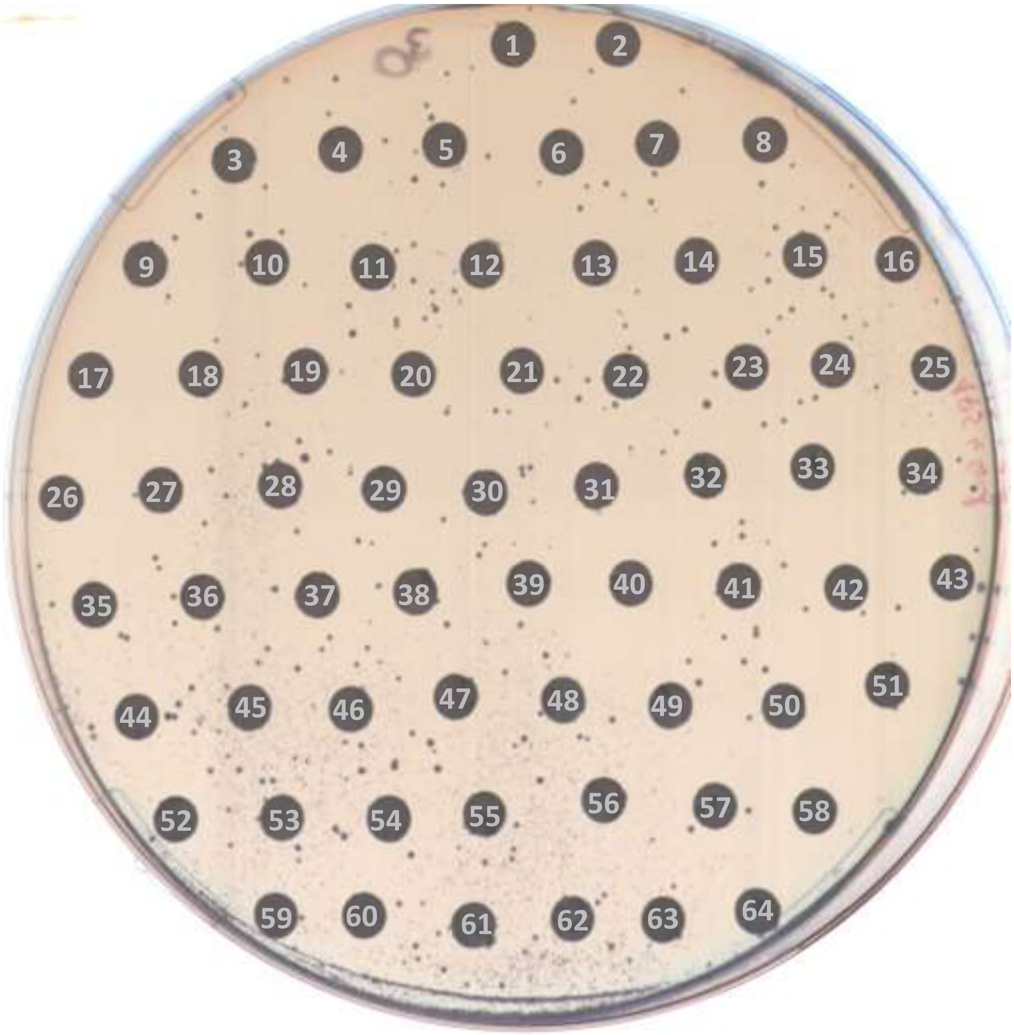

48 h

### PLATE 30

| no. | compound | no. | compound | no. | compound |
|---|---|---|---|---|---|
| - | - | 22 | Fluticasone furoate | 44 | Isosorbide dinitrate |
| 1 | Thonzonium (bromide) | 23 | Epristeride | 45 | (+)-Camphor |
| 2 | Rizatriptan (benzoate) | 24 | (-)-Limonene | 46 | Oleic acid |
| 3 | Cefotetan | 25 | Choline (bitartrate) | 47 | Abarelix |
| 4 | Mebrofenin | 26 | Mizolastine | 48 | Cabergoline |
| 5 | Lanreotide (acetate) | 27 | Paliperidone | 49 | Propoxycaine (hydrochloride) |
| 6 | Berbamine (dihydrochloride) | 28 | Bentiromide | 50 | Ethylparaben |
| 7 | 18β-Glycyrrhetinic acid | 29 | Nylidrin (hydrochloride) | 51 | Tandospirone (citrate) |
| 8 | Eugenol | 30 | Sivelestat (sodium) | 52 | Laurocapram |
| 9 | Indobufen | 31 | Levomepromazine | 53 | Dipotassium glycyrrhizinate |
| 10 | Pyrimethamine | 32 | Dihydroergocristine (mesylate) | 54 | Ondansetron |
| 11 | Triptonide | 33 | Dienestrol | 55 | Lurasidone (Hydrochloride) |
| 12 | Pipobroman | 34 | Cetrorelix (Acetate) | 56 | Lenampicillin (hydrochloride) |
| 13 | Fosfomycin (calcium) | 35 | Fenoterol (hydrobromide) | 57 | Squalene |
| 14 | Alizapride hydrochloride | 36 | Vinburnine | 58 | Sinomenine hydrochloride |
| 15 | Disodium succinate | 37 | Perospirone | 59 | Fenofibrate |
| 16 | Amcinonide | 38 | Ensulizole | 60 | Allylestrenol |
| 17 | Bromperidol | 39 | Piperidolate | 61 | Isocarboxazid |
| 18 | Cimetropium (Bromide) | 40 | Flupentixol dihydrochloride | 62 | Oleanolic Acid |
| 19 | Sophoridine | 41 | Teneligliptin | 63 | Argipressin |
| 20 | Harringtonine | 42 | Estradiol valerianate | 64 | (Z)-Thiothixene |
| 21 | Arotinolol | 43 | Linoleic acid | - | - |

### PLATE 31

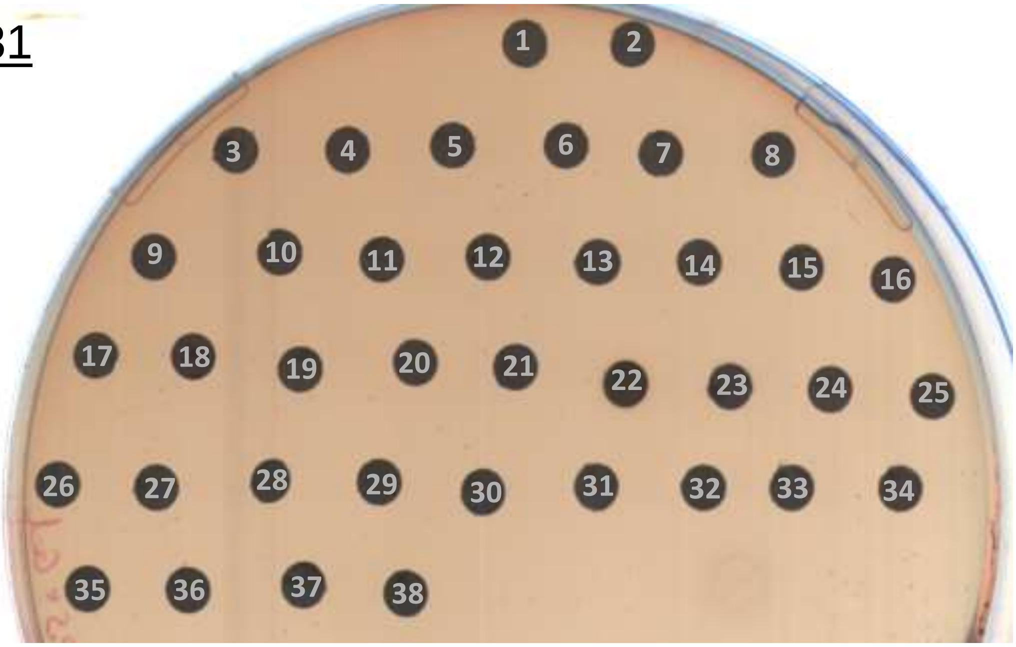

24 h

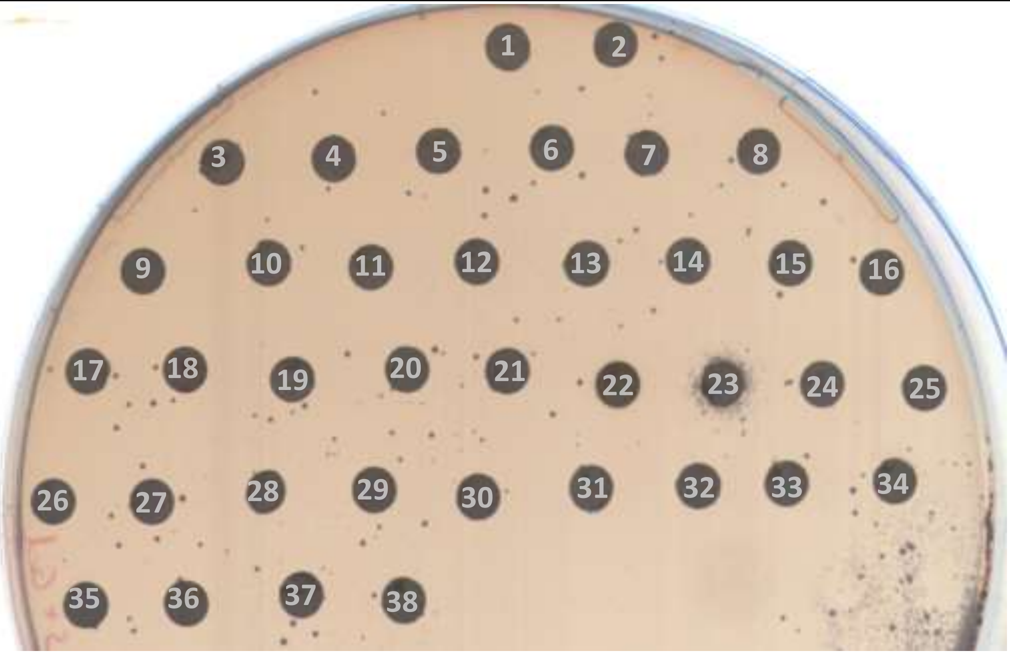

48 h

### PLATE 31

| no. | compound | no. | compound | no. | compound |
|---|---|---|---|---|---|
| - | - | 22 | Baicalein | - | - |
| 1 | Hyodeoxycholic acid | 23 | Meisoindigo | - | - |
| 2 | Treosulfan | 24 | Camptothecin | - | - |
| 3 | Metergoline | 25 | Bithionol | - | - |
| 4 | Thymol | 26 | N-Acetyl-L-tyrosine | - | - |
| 5 | Dehydroandrographolide succinate | 27 | L-Cysteine (hydrochloride) | - | - |
| 6 | Triptolide | 28 | Dimethyl fumarate | - | - |
| 7 | Treprostinil | 29 | L-Cycloserine | - | - |
| 8 | Abaloparatide (TFA) | 30 | Urapidil (hydrochloride) | - | - |
| 9 | Cefpodoxime Proxetil | 31 | Taltirelin | - | - |
| 10 | EC5026 | 32 | Afoxolaner | - | - |
| 11 | Isocorydine | 33 | Midodrine (hydrochloride) | - | - |
| 12 | Alfuzosin (hydrochloride) | 34 | Amiodarone (hydrochloride) | - | - |
| 13 | Panaxatriol | 35 | Levocetirizine (dihydrochloride) | - | - |
| 14 | Mizoribine | 36 | Perhexiline maleate | - | - |
| 15 | Diloxanide furoate | 37 | Gramicidin | - | - |
| 16 | Tetrahydropalmatine | 38 | Talaporfin (sodium) | - | - |
| 17 | S-Adenosyl-L-methionine (disulfate tosylate) | - | - | - | - |
| 18 | Indirubin | - | - | - | - |
| 19 | Fumagillin | - | - | - | - |
| 20 | Baicalin | - | - | - | - |
| 21 | Indigo | - | - | - | - |
